# AI-driven Discovery of Morphomolecular Signatures in Toxicology

**DOI:** 10.1101/2024.07.19.604355

**Authors:** Guillaume Jaume, Thomas Peeters, Andrew H. Song, Rowland Pettit, Drew F. K. Williamson, Lukas Oldenburg, Anurag Vaidya, Simone de Brot, Richard J. Chen, Jean-Philippe Thiran, Long Phi Le, Georg Gerber, Faisal Mahmood

**Affiliations:** Department of Pathology, Brigham and Women’s Hospital, Harvard Medical School, Boston, MA; Department of Pathology, Massachusetts General Hospital, Harvard Medical School, Boston, MA; Cancer Program, Broad Institute of Harvard and MIT, Cambridge, MA; Cancer Data Science Program, Dana-Farber Cancer Institute, Boston, MA; Institute of Animal Pathology, Vetsuisse, University of Bern, Switzerland; COMPATH, Institute of Animal Pathology, University of Bern, Switzerland; Bern Center for Precision Medicine, University of Bern, Switzerland; Department of Pathology & Laboratory Medicine, Emory University School of Medicine, Atlanta, GA; Health Sciences and Technology, Harvard-MIT, Cambridge, MA; Signal Processing Laboratory, EPFL, Lausanne, Switzerland; Harvard Data Science Initiative, Harvard University, Cambridge, MA

## Abstract

Early identification of drug toxicity is essential yet challenging in drug development. At the preclinical stage, toxicity is assessed with histopathological examination of tissue sections from animal models to detect morphological lesions. To complement this analysis, toxicogenomics is increasingly employed to understand the mechanism of action of the compound and ultimately identify lesion-specific safety biomarkers for which *in vitro* assays can be designed. However, existing works that aim to identify morphological correlates of expression changes rely on qualitative or semi-quantitative morphological characterization and remain limited in scale or morphological diversity. Artificial intelligence (AI) offers a promising approach for quantitatively modeling this relationship at an unprecedented scale. Here, we introduce GEESE, an AI model designed to impute morphomolecular signatures in toxicology data. Our model was trained to predict 1,536 gene targets on a cohort of 8,231 hematoxylin and eosin-stained liver sections from *Rattus norvegicus* across 127 preclinical toxicity studies. The model, evaluated on 2,002 tissue sections from 29 held-out studies, can yield pseudo-spatially resolved gene expression maps, which we correlate with six key drug-induced liver injuries (DILI). From the resulting 25 million lesion-expression pairs, we established quantitative relations between up and downregulated genes and lesions. Validation of these signatures against toxicogenomic databases, pathway enrichment analyses, and human hepatocyte cell lines asserted their relevance. Overall, our study introduces new methods for characterizing toxicity at an unprecedented scale and granularity, paving the way for AI-driven discovery of toxicity biomarkers.

**Live demo:** https://mahmoodlab.github.io/tox-discovery-ui/

## Introduction

Identifying and characterizing the potential toxicity of a drug early in its development is a major challenge for the pharmaceutical industry^1–3^. At the preclinical phase, toxicity is assessed in animal models through histological examination of tissue sections, in which pathologists report drug-induced lesions and abnormalities to determine the dose-response relationship of the compound (**Fig. 1a**). Despite advancements of *in vitro* assays for early toxicity detection, safety concerns remain the leading cause of drug attrition at the preclinical stage^4^. For these reasons, preclinical research increasingly relies on toxicogenomics^5–8^, such as gene expression profiling, to develop a mechanistic understanding of the drug action. By correlating changes in gene expression with specific morphological lesions, such as cellular necrosis, investigators can characterize the morphomolecular response of the compound. When validated across multiple studies and compounds, lesion-specific genetic biomarkers can serve as novel indicators for early toxicity detection from *in vitro* testing, overall enhancing the likelihood of successfully transitioning to early-stage clinical development^9^.

**Figure 1:**
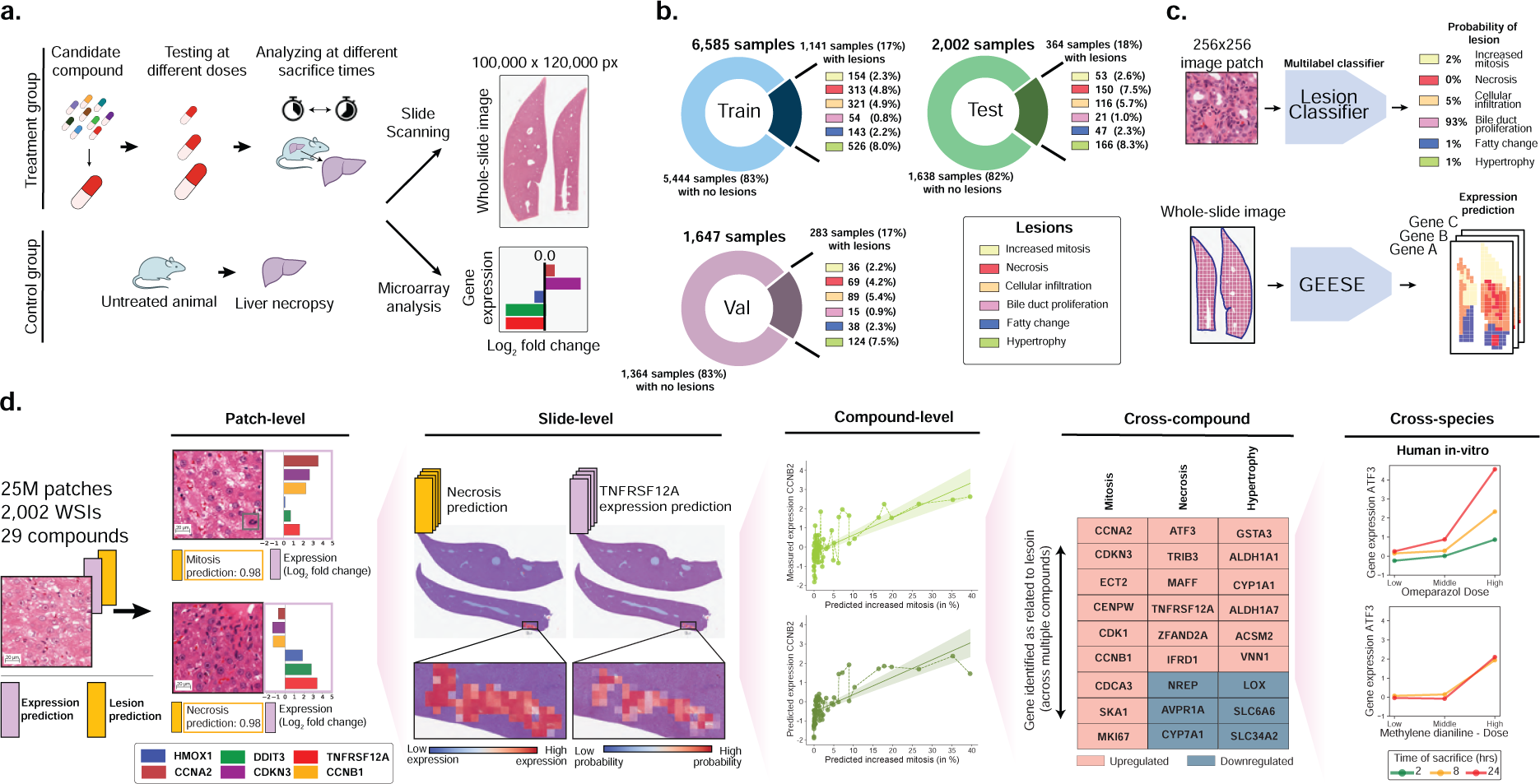
Study overview. **a.** At preclinical stage, drug candidates undergo toxicity assessment in animal models to characterize the dose-response relationship of the compound based on histological examination. Toxicogenomics can be employed to complement the compound characterization. **b.** Overview of TG-GATEs composed of 156 preclinical safety studies (and compounds) accounting for 10,234 pairs of hematoxylin and eosin (H&E) whole-slide images and gene expression profiles. TG-GATEs is split into a development set (127 studies, 8,232 slides) and a test set (29 studies, 2,002 slides). **c.** We developed two independent prediction models: (1) a morphological lesion prediction model (denoted as Lesion classifier), which classifies 256*×*256 pixels (or 128 *µ*m) image patches into six lesions, and (2) a gene expression regression model (GEESE), which predicts bulk expression of 1,536 gene targets from an input tissue section. Feature attribution enables GEESE to derive patch-level expression profiles to yield pseudo-spatially resolved expression maps. **d.** The resulting output forms a dataset of 25 million predicted patch-level morphology-expression pairs, which we use for inferring and validating morphomolecular signatures across several scales, from patches (small regions of interest) to slides (entire tissue sections) to compounds (can include dozens of slides), then across several compounds, and finally across species (rat *in vivo* to human *in vitro*).

However, prior studies investigating relationships between gene expression changes and morphological lesions have relied on pathology reports for morphological characterization of the tissue, which provides limited information compared to the tissue itself^10–14^. Therefore, these assessments remain qualitative and semi-quantitative, and may be subject to high inter-observer variability, especially for severity scoring of lesions, which limits the ability to detect subtle associations. In addition, existing works and toxicity databases aggregating findings, such as the Comparative Toxicogenomics Database^15–17^ are limited to reporting biomarkers linked to drug-induced injury rather than linking specific lesions, such as fatty change, to expression changes.

To overcome these limitations, artificial intelligence (AI) and computational pathology offer a promising approach for quantitatively and spatially modeling the relationship between morphology and gene expression changes at scale^18–20^. Multiple works have shown the ability to predict molecular profiles, such as gene mutations^21–27^, microsatellite instability ^28^, and gene expression ^29–32^, directly from whole-slide images (WSIs). This direction holds promise for identifying morphological correlates of molecular alterations and ultimately for biomarker discovery^25, 33^. Existing works in this area have primarily focused on cancer cohorts, typically with sample sizes under 1,000 cases per disease, such as those from The Cancer Genome Atlas Program (TCGA). Moreover, most studies have been limited to qualitative analyses, such as examining attention weights or gradient attributions, rather than quantitative assessments. Consequently, there remains an unmet need for quantitative, objective, and scalable methods to analyze morphological correlates of gene expression.

Here, we introduce the first AI model designed to identify and impute morphomolecular signatures in toxicology data by connecting specific morphologies to expression changes. Our model, named Gene Expression Regressor (GEESE), is a deep learning architecture that predicts bulk expression levels of 1,536 selected gene targets from digitized H&E-stained liver sections (whole-slide images, WSIs). GEESE employs a weakly supervised training approach to predict slide-level labels (i.e., gene expression) without requiring patch-level annotations and enabling scalable training on large datasets. We trained GEESE on 8,231 hematoxylin and eosin (H&E) WSIs from *Rattus norvegicus* liver, spanning 127 preclinical drug safety studies. The model was evaluated on an independent test set of 29 studies comprising 2,002 WSIs and expression profile pairs (**Fig. 1b**). GEESE’s unique architecture enables fine-grained attribution of gene expression predictions, yielding pseudo-spatially resolved gene expression maps for all test samples (**Fig. 1c**). To connect gene expression with morphological lesions, we additionally developed a morphological classification model to identify six common drug-induced liver injuries (DILI), including necrosis, fatty change, and increased mitosis. By correlating the pseudo-spatially resolved expression predictions from GEESE with the lesion predictions, we generated a dataset of 25 million morphology-expression pairs.

From this analysis, we established a robust and quantitative relation between up and down-regulated genes and morphological lesions, with multiple associations being preserved across multiple compounds (**Fig. 1d**). We curated lists of genes linked to each of the studied lesions and identified biomarkers that were corroborated with public databases, such as the Comparative Toxicogenomics Database^15–17^, and pathway-enrichment analyses. We further validated these gene signatures against *in vitro* primary human hepatocyte cell lines, providing an initial assessment of translatability to humans. Overall, our study introduces new methods to understand toxicity and its underlying morphomolecular mechanisms at an unprecedented scale and granularity, paving the way for enhanced prediction and mechanistic understanding of compounds.

## Results

### Study overview

Our study leverages the publicly available Toxicogenomics Project-Genomics Assisted Toxicity Evaluation System (TG-GATEs) dataset^34^, a collection of preclinical drug safety studies (**Materials and Methods**, section **TG-GATEs protocol** and **table S1**). TG-GATEs studies were acquired as part of the Japanese Toxicogenomics Project consortium designed to test the hepatotoxicity of known drugs and chemicals after *in vivo* compound exposure on *Rattus norvegicus*. Here, we collected data from 156 drug safety studies accounting for 10,234 pairs of haematoxylin and eosin (H&E) whole-liver tissue sections (20*×* magnification, 0.49*µ*m/px) with the corresponding gene expression profile measured with mRNA microarrays^35^. Each slide represents the morphological changes caused by administering a specific dose of a compound at a particular time post-administration. In addition, each slide was annotated with morphological lesions identified by toxicologic pathologists, such as reporting the presence of hepatocellular hypertrophy – with lesions that might be drug-induced or spontaneous (**Materials and Methods**, section **Histopathology acquisition and annotation**).

Analogously, bulk gene expression profiles was performed to encode the molecular landscape from the tissue section, which the action of the compound could have altered. Gene expression was measured with mRNA microarrays, which provide a bulk whole-transcriptome description post-drug administration. We limit our study from 31,042 probes to a set of 1,536 genes selected to cover a large biological space relevant to toxicology (**Materials and Methods**, section **Gene expression of *in vivo* rat studies**). Specifically, through complementary *a priori* knowledge-driven and data-driven approaches, we filtered the full set of genes to a subset that satisfies either of the two conditions: (1) genes that are biologically relevant to toxicology (linked to liver metabolism and liver response to injury), as identified through works such as T1000^36^; and (2) genes whose measured expression levels were highly correlated with the presence of drug-induced lesions were included (e.g., *TMBIM1*, a gene involved in death receptor binding activity and necrosis).

To rigorously assess the generalizability of our model to unseen studies, we split TG-GATEs into a set for training and validation that comprises 127 studies and 8,232 slides, and a test set that comprises 29 studies and 2,002 slides (**Fig. 1b** and **Materials and Methods**, section **Dataset split**). Overall, 1,788 slides (17% of the TG-GATEs dataset) report morphological lesions: 1,141 (17%) in train, 283 (17%) in validation, and 364 (18%) in test. In this study, we focus our analysis on compounds that induce the following six commonly found morphological lesions: abnormal increases in mitotic figures (2.4% of slides in TG-GATEs), necrosis (5.2% of slides), cellular infiltration (5.1% of slides), bile duct proliferation (0.9% of slides), fatty change (2.2% of slides), and hepatocellular hypertrophy (8.0% of slides).

### Weakly-supervised expression prediction

We introduce the gene expression regressor (GEESE) method, a deep learning model based on multiple instance learning (MIL)^37–39^ that can predict the expression profile associated with an input whole-slide image (WSI). Following the MIL paradigm, we tessellate the slide into 256*×*256 pixels patches. We use a pre-trained vision encoder to extract patch embeddings from each patch. Here, to reduce the domain gap between the source training domain (such as natural images^40^ or human histology^41–43)^ and the target domain (rodent histology), we trained a Vision Transformer from scratch with iBOT^44–46^, a self-supervised learning (SSL) model^47, 48^. Our SSL model was trained on 15 million patches extracted in 46,734 slides from TG-GATEs (**Materials and Methods**, section **iBOT pretraining**). We then trained a MIL regression model predicting gene expression from patch embeddings within the slide. Our method, GEESE, can extract patch-level gene expression scores using a multilayer perceptron (MLP) mapping the patch embedding to the expression profile. By summing patch-level predictions, GEESE can derive slide-level expression levels, which represent the predicted log2 fold change expression quantifying expression changes between the control and the tested configuration (zero means no change). We employed the mean squared error (MSE) between the predicted and measured expression to train GEESE end-to-end with the patch embeddings. This MIL formulation naturally yields patch-level expression predictions, thereby enabling a pseudo-spatially resolved expression map, where the resolution is given by patch size^29^. This differs from widely employed attention-based methods^38, 39^ or gradient attribution methods^49^, which rely on surrogate attribution. Additional information is provided in the **Materials and Methods**, section **Gene expression regressor (GEESE)**.

We trained GEESE on a set of 127 studies (8,232 slides) and evaluated it on 29 held-out studies (2,002 slides) (**fig. S1 and S2**). The model prediction is assessed using the Pearson correlation and area under the ROC curve (AUC) adapted to regression (**Materials and Methods**, section **Evaluation and implementation**). Predicted expression levels of several genes, such as *TNFRSF12A* (involved in inflammation and cell death) and *SLC10A1* (part of the sodium/bile acid co-transporter family), show a high correlation with observed expression levels: r=0.722 (95% CI: [0.675, 0.762]) and r=0.688 (95% CI: [0.630, 0.733]), respectively. When considering all selected genes, the average correlation is 0.29 and increases to 0.63 when considering the top 100 best-predicted genes. We attribute these large variations in prediction to the fact that (1) some genes (expressed or not) might not be reflected in the tissue morphology making them undetected by GEESE, (2) other genes might have low expression in the test studies rendering detection harder due to noise, and (3) some learned expression profiles associated with certain morphologies in training might not generalize to test studies.

We additionally investigated model performance stratified by compound. Specifically, we identified the top 100 best-predicted genes and computed the average Pearson correlation for each study (**Fig. 2a**). Overall, we observe large variations from one study to another. In some studies, such as thioacetamide and methylene dianiline, predictions have high correlations with ground truth (*r >*0.8), while in others, such as carboplatin, correlations are smaller (*r <*0.5). To understand such discrepancies, we investigated the percentage of slides per study that reported on the six considered lesions (background or drug-induced). We observe that studies with poorly predicted expression usually corresponded to compounds with little to no reported lesions. This suggests that the model uses the presence of morphological lesions to predict expression.

**Figure 2:**
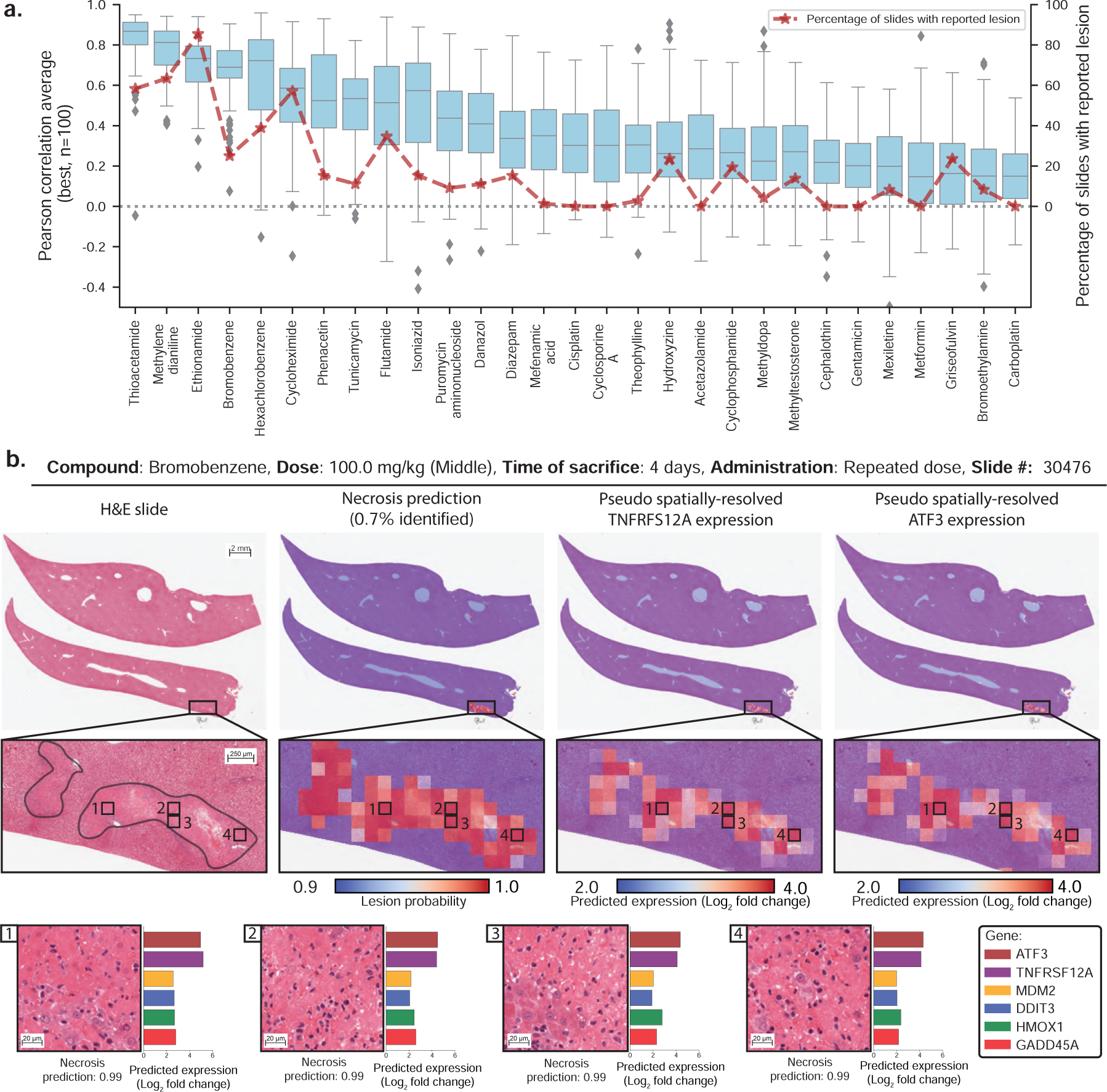
Gene expression profiling from whole-slide images using GEESE. **a.** Gene expression prediction of the top 100 best-predicted genes evaluated using Pearson correlation and stratified by compound from the test set. The percentage of slides with lesions is displayed in red. Boxes indicate quartile values of gene-level Pearson correlation, with the center line indicating the 50th percentile. Whiskers extend to data points within 1.5*×* the interquartile range. **b.** Example of a liver section after exposure to bromobenzene (left). Overlay of patch-level necrosis prediction. Predictions below 90% confidence are represented in blue, and high-probability predictions are represented in red (center-left). Pseudo spatially-resolved gene expression heatmaps of genes *TNFRSF12A* (center-right) and *ATF3* (right). Examples of high-probability necrotic patches from the slide pseudo gene expression of *ATF3*, *TNFRSF12A*, *MDM2*, *DDIT3* (also known as CHOP), *HMOX1*, and *GADD45A* (bottom).

We further inspected the slide-level expression prediction and ground truth in thioacetamide and methylene dianiline, the two best-predicted compounds for the genes *SLC10A1*^50^ and *TNFRSF12A*^51^. Notably, we observe a substantial Pearson correlation (*r ∈* [0.8, 0.9]), where the presence of a lesion (as per TG-GATEs annotations) appears to be linked to *SLC10A1* downregulation and *TNFRSF12A* upregulation (**fig. S1c**). Yet, samples with lesions highlight different expression levels (from 0 to 4 log2 fold change in *TNFRSF12A* expression in methylene dianiline), which require quantification methods for deeper analyses.

### Analysis of morphological correlates of expression changes

As part of routine histological assessment, identified morphological lesions are reported with a score describing the extent and severity of the lesion (in TG-GATEs, minimal, slight, moderate, severe). However, lesion scoring remains based on qualitative or semi-quantitative assessment and, as such, lacks consistency within and across studies, making it impractical for robust quantification analyses. Instead, we leverage our in-domain vision encoder as a foundation for training a lesion classifier. We gathered patch-level annotations from six commonly found lesions: increased mitosis, necrosis, cellular infiltration, bile duct proliferation, fatty change, and hypertrophy (**Materials and Methods**, section **Patch encoding and lesion classification**, **table S3** and **table S4**). In total, we acquired 24,631 patch annotations with lesions extracted from 3,458 liver slides and 13,888 normal patch annotations from 3,531 slides, accounting for a total of 38,519 patch annotations. The vision encoder was then fine-tuned on these annotations and trained to minimize a multilabel binary cross-entropy objective on the six lesions of interest (patches can include multiple lesions). Our fine-tuned lesion classifier reaches an average performance of 98.9% macro-AUC across all lesions, making it a reliable and robust predictive tool for subsequent analysis.

By running lesion classification on all 2,002 test slides, we can obtain high-quality patch-level predictions on 25 million patches, where each prediction describes the likelihood that a given lesion is present in the patch. Subsequently, we can compare the lesion prediction with the pseudo-spatially-resolved gene expression obtained with GEESE (**Fig. 2b**). For instance, when analyzing a sample from the bromobenzene study (administration of 100mg/kg for four days, each day), we observe that the gene expression heatmap of *TNFRSF12A*^51^ and *ATF3*^52^ focuses on the same regions. In addition, both regions largely overlap with the prediction of necrosis. This suggests that GEESE uses the presence of necrosis to predict the increased expression of both genes. This finding is not limited to *TNFRSF12A* and *ATF3*, as other genes highlight the same trend such as *MDM2*, *DDIT3*, *HMOX1*, and *GADD45A*, genes involved in a diverse array of processes often related to inflammation, oxidative stress, and cell death such as positive regulation of apoptotic process or response to endoplasmic reticulum (ER) stress.

When conducting a similar analysis across several test studies, such as on thioacetamide, ethionamide, methylene dianiline, and methyldopa, we observe that the presence of necrosis aligns with the upregulation of *TNFRSF12A* and *ATF3*, suggesting that this finding is compound-agnostic – or that the mechanism of action of the compound is linked to both these genes (**fig. S2**).

While this analysis was conducted with necrosis, similar behavior is observed with other lesions. The upregulation of *CCNA2*, *KNSTRN*, *CDKN3*, *CCNB1*, *ARHGAP11A*, and *HMMR* aligns with the presence of increased mitosis (**fig. S3**); the upregulation of *CXCL1*, *BCL2A1*, *S100A4*, *CXCL10*, *EVI2A*, and *FILIP1L* aligns with the presence of cellular infiltration (**fig. S4**); the upregulation of *SERPINA7*, *BEX4*, *CDH13*, *CLDN7*, *CD24* and downregulation of *SERPINA4* aligns with the presence of bile duct proliferation (**fig. S5**); the upregulation of *ACOT1*, *GSTP1*, *ACOT2*, *CYP1A1*, *HID1* and the downregulation of *SLC6A6* aligns with the presence of fatty change (**fig. S6**); and the upregulation of *GSTA3*, *ALDH1A1*, *ADGRG2*, *RGD1559459*, and the downregulation of *PLVAP* and *LOX* aligns with the presence of hypertrophy (**fig. S7**). Overall, by comparing patch-level lesion predictions with the pseudo-gene expression map from GEESE, strong relationships are observed between the presence of each of the six lesions and the upregulation or downregulation of specific genes.

### Single-study morphomolecular analysis

We expanded the analysis to all WSIs within a study to refine and characterize the morpho-molecular relationships previously identified. We rely on three quantitative measures for indepth investigation: 1) the gene expression (normalized as log2 fold change) as measured with mRNA microarray, 2) the predicted pseudo-spatially-resolved expression map, and 3) the predicted size, type, and location of each lesion. We emphasize that the latter two measures are enabled by our proposed lesion classifier and GEESE architecture.

By measuring the Pearson correlation between the measured slide-level expression and the predicted lesion size, we observe a subset of genes with high correlation (*r >*0.7, **Fig. 3a**). This is exemplified with increased mitosis after administration of danazol (**Fig. 3**), cellular infiltration in methylene dianiline (**fig. S8**), bile duct proliferation in methylene dianiline (**fig. S9**), fatty change in ethionamide (**fig. S10**), and hypertrophy in hexachlorobenzene (**fig. S11**). This analysis further shows that genes showing a high correlation between measured expression and predicted lesion size overlap with the genes best predicted by GEESE. In addition, we observe that certain genes are not associated with any of the considered lesions.

**Figure 3:**
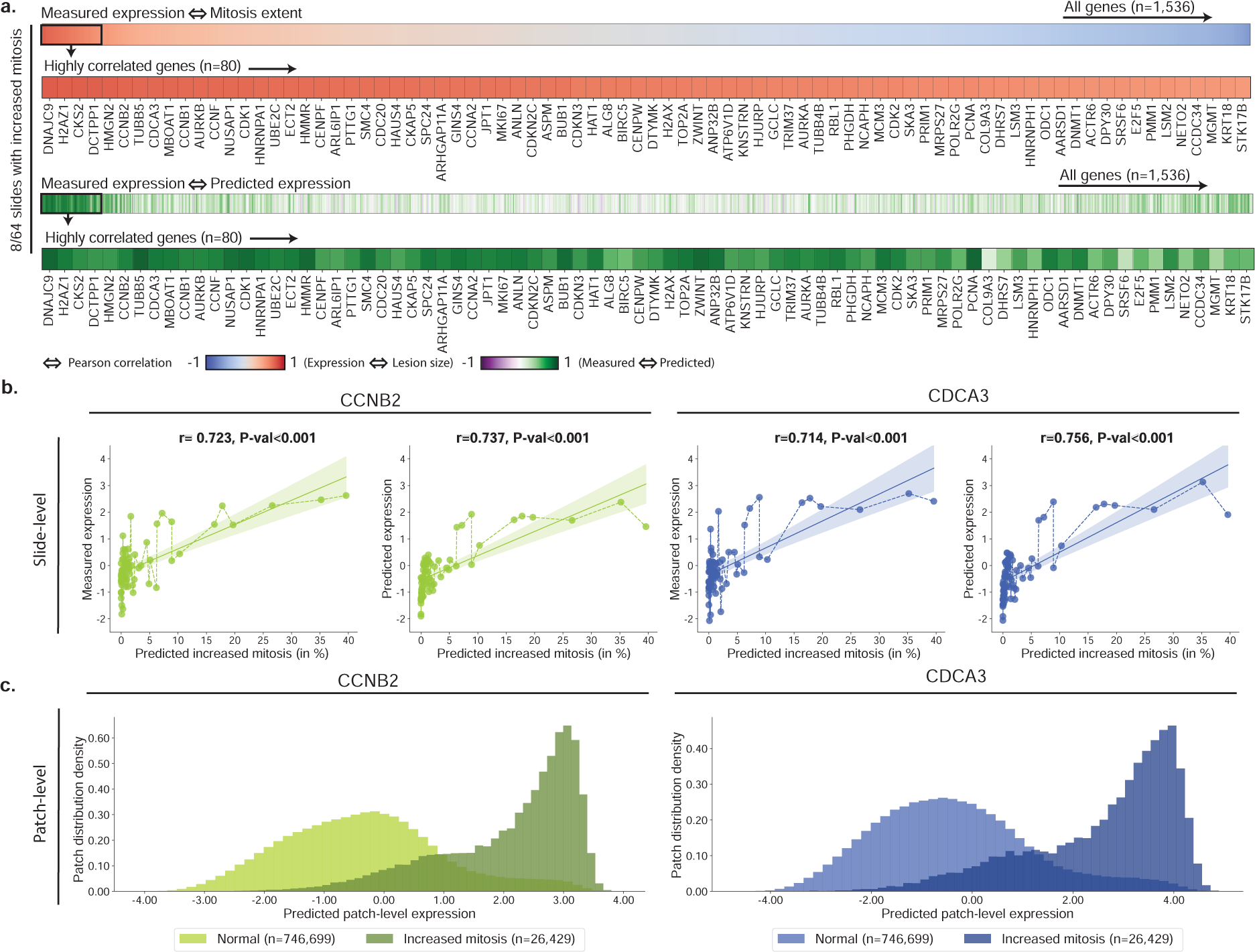
Morphomolecular analysis of increased mitosis in danazol. **a.** For each slide-expression pair, we correlate the predicted percentage of mitosis found in the slide with the measured gene expression. Genes with a high correlation between measured expression and predicted lesion extent can be seen as part of the morphomolecular signature associated with the compound. TG-GATEs original annotations report 8/64 slides with increased mitosis in danazol. **b.** Correlation between the estimated percentage of the mitosis in a slide and the gene expression of *CCNB2* and *CDCA3* (left: measured, right: predicted). P-value derived from testing the two-sided null hypothesis of non-correlation. **c.** Density distribution of patch-level expression for patches predicted as normal by the lesion classifier (n=746,699 patches) and patches predicted as containing mitosis (n=26,429). Patches were extracted from the 64 WSIs of the danazol study.

When focusing on specific genes, such as *CCNB2* (Cyclin B2) and *CDCA3* (Cell Division Cycle Associated 3) in danazol, we observe how the measured expression varies with the predicted lesion size (**Fig. 3b**). We find *CCNB2* and *CDCA3* to be increasingly upregulated as the predicted lesion size increases (r*>*0.7). The same observation holds when analyzing the slide-level gene prediction (r*>*0.7 between predicted expression and predicted mitosis). We further analyzed the distribution of the predicted patch-level expression for these two genes. To this end, we assign a label to each patch (lesion or normal), according to whether the predicted lesion probability crosses lesion-specific thresholds (**Materials and Methods**, section **Lesion classifier**). We observe a bi-modal distribution, where normal patches (*i.e.*, no sign of lesion) have an average predicted expression centered around zero and patches with mitosis are predicted as upregulated (**Fig. 3c**). Overall, by comparing gene expression in patches with and without lesions, we can establish precise and quantitative relationships between up and downregulation of genes and the extent of lesions.

### Cross-study analysis of morphomolecular signatures

The next aim is to study whether these signatures are preserved across studies, for instance, if the molecular correlates of mitosis identified in response to danazol are also found in other compounds. This allows us to determine if observed genetic markers linked to particular morphologies remain consistent across compounds, which would be suggestive of specific toxicological mechanisms. To this end, we compiled a list of studies where a given lesion is observed (**table S5** and **table S6**), for instance, the eight studies from the TG-GATEs test set with mitosis (**fig. S12**). For each slide from each study, we extracted all patches with lesions (for instance, 157,414 patches predicted with mitosis). A detailed description of the number of patches extracted for each study can be found in **table S7**. We then computd the macro-average of the predicted expression for the selected patches across all studies and rank them from the highest to lowest. Genes with the highest (absolute) expression are expected to be the most linked to the lesions (**Materials and Methods**, section **Post-hoc morphomolecular signature analysis**). We conducted similar analyses for all lesions: necrosis (four studies, **fig. S13**), cellular infiltration (four studies, **fig. S14**), bile duct proliferation (two studies, **fig. S15**), fatty change (four studies, **fig. S16**), and hypertrophy (eleven studies, **fig. S17**).

An overview of the analyses is shown in **Fig. 4** where we highlight the 40 most upregulated (in red) and most downregulated (in blue) genes for all lesions. Some genes are consistently upregulated (such as *ABCC3* and *GPX2*) or downregulated (such as *NOX4*, *CAR3*, and *OAT*) for multiple lesions (**table S8**). These genes do not appear to be lesion-specific but may rather be indicative of general toxic exposure (for instance, *ABCC3* is involved in transporting organic anions and drugs out of cells). Interestingly, some of these genes are those best predicted by GEESE, such as *KLF6*, *NOX4*, and *CAR3*, that have Pearson correlations of 0.717 (95% CI: [0.672, 0.759]), 0.639 (95% CI: [0.605, 0.672]), 0.698 (95% CI: [0.659, 0.736]), respectively, between predicted and observed expression on all test studies (**fig. S1**). A large number of the identified genes (both up and downregulated) are linked to drug-induced liver injury according to the Comparative Toxicogenomics Database (CTD)^15–17^, either through direct therapeutic or mechanistic evidence (genes with filled black star in **Fig. 4**) or via inferred evidence from network-based analysis (star with black contour in **Fig. 4**). For instance, 23/40 of the most upregulated genes associated with hypertrophy are referenced in CTD as direct or inferred evidence (inference score*>*400), such as CYP1A1, which encodes a member of the cytochrome P450 superfamily of enzymes known for catalyzing reactions involved in drug metabolism^53, 54^. This analysis validates the ability of GEESE to identify highly relevant gene sets connected to toxicity.

**Figure 4:**
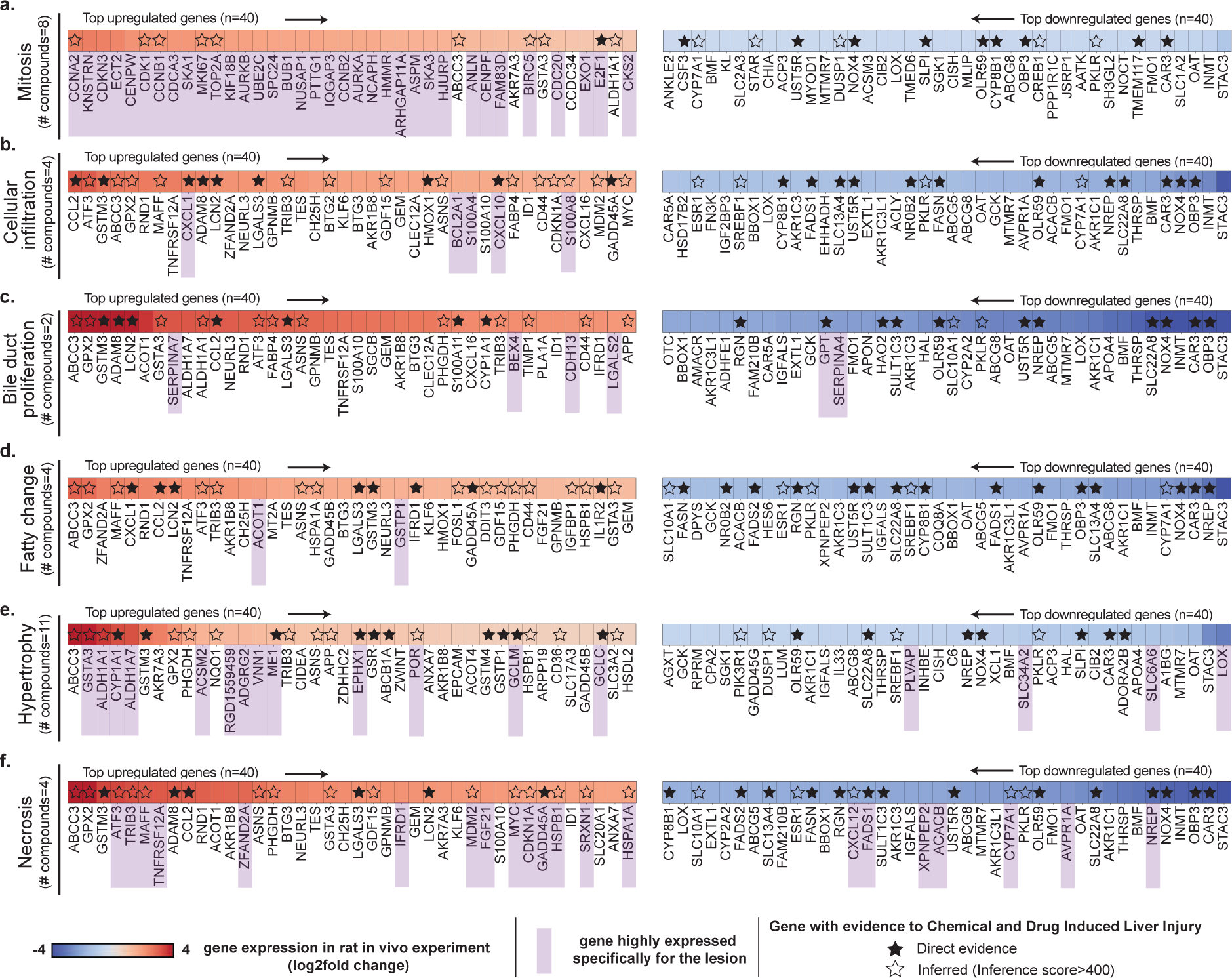
Cross-study morphomolecular analysis. Heatmap illustrating the mean expression of genes in patches displaying specific lesions, comparing the top 40 upregulated and top 40 downregulated genes for each lesion type. Panels **a.** mitosis, **b.** cellular infiltration, **c.** bile duct proliferation, **d.** fatty change, **e.** hypertrophy, and **f.** necrosis details the gene expression dynamics, where each analysis spans across multiple compounds (**table S5**). Genes are ranked by their absolute expression, with upregulated genes indicated in red (descending order) and downregulated genes in blue (ascending order). Genes expressed for each specific lesion are highlighted in purple. Genes with known connections to drug-induced liver injury (DILI) are marked with a star (either as direct or inferred evidence). Direct evidence refers to known mechanistic and/or therapeutic connections between the gene and DILI. Inferred evidence with DILI refers to genes with an inference score*>*400 as per the Comparative Toxicogenomics Database (CTD)^15–17^ (inference score measured with the similarity between CTD chemical–gene–disease networks and a similar scale-free random network).

Certain genes exhibit changes in expression only for a specific lesion, indicating a closer association with the lesion itself rather than general toxic exposure. To identify these lesion-specific genes, we compared the average predicted patch-level expression containing a lesion of interest against the expression of all other patches with lesions. A gene was then considered lesion-specific if its expression was significantly larger than the ones of the other five lesions (**Materials and Methods**, section **Post-hoc morphomolecular signature analysis**). For each lesion, we identified a varying number of lesion-specific genes (genes marked in purple in **Fig. 4**). For instance, mitosis exhibits a distinct molecular signature, likely due to its unique nature compared to other lesions, such as cellular infiltration and fatty change, that can co-occur with other more prominent lesions in our dataset. To assess the relevance of the identified lesion-specific molecular signatures, we conducted a pathway enrichment analysis using the Rat Genome Database^55^ that aggregates previously established biological processes in rats and humans (**Fig. 5a,b,c** and **fig. S12b, S13b, S14b, S15b, S16b and S17b** for lesion-wise analysis). GEESE-identified gene sets are significantly enriched for pathways linked to different lesions. For instance, out of the 51 genes uniquely linked to mitosis (**Fig. 5a and table S10**), such as *CDK1*^56^ and *CCNB1*, 40 are involved in the cell cycle pathway in rats (p-value=6.53E-34), with 41 also involved in the equivalent human pathway (p-value=5.31E-33), 29 genes are involved in the chromosome segregation pathway in rats and humans (p-value=2.07E-36 and p-value=1.69E-33, respectively), and 25 genes are involved in cell division (p-value=2.09E-26), with 34 also involved in the equivalent human pathway (p-value=1.96E-37). When conducting a similar analysis in necrosis (**Fig. 5b and table S10**), we found that out of the 33 upregulated identified genes, such as *ATF3*^52^, *TNFRSF12A* ^51^, *DDIT3*^57–59^ and *TRIB3*^14^, 21 genes are involved in the apoptotic process in rats and humans (p-value=1.30E-14 and p-value=1.25E-13, respectively), and 20 genes are involved in the cellular response to stress in rats (p-value=1.24E-13), with 21 also involved in humans (p-value=3.18E-14). The 18 upregulated genes linked to hypertrophy (e.g., *ALDH1A1*, *ACSM2*^60^ and *VNN1*^61^) are also significantly enriched for several metabolic pathways such as fatty acid, lipid, and glucose metabolism. Similar analyses conducted on the three other lesions further assert the relevance of GEESE-identified genes. This illustrates the ability of GEESE to identify relevant lesion-specific genetic biomarkers with promising transferability to humans.

**Figure 5:**
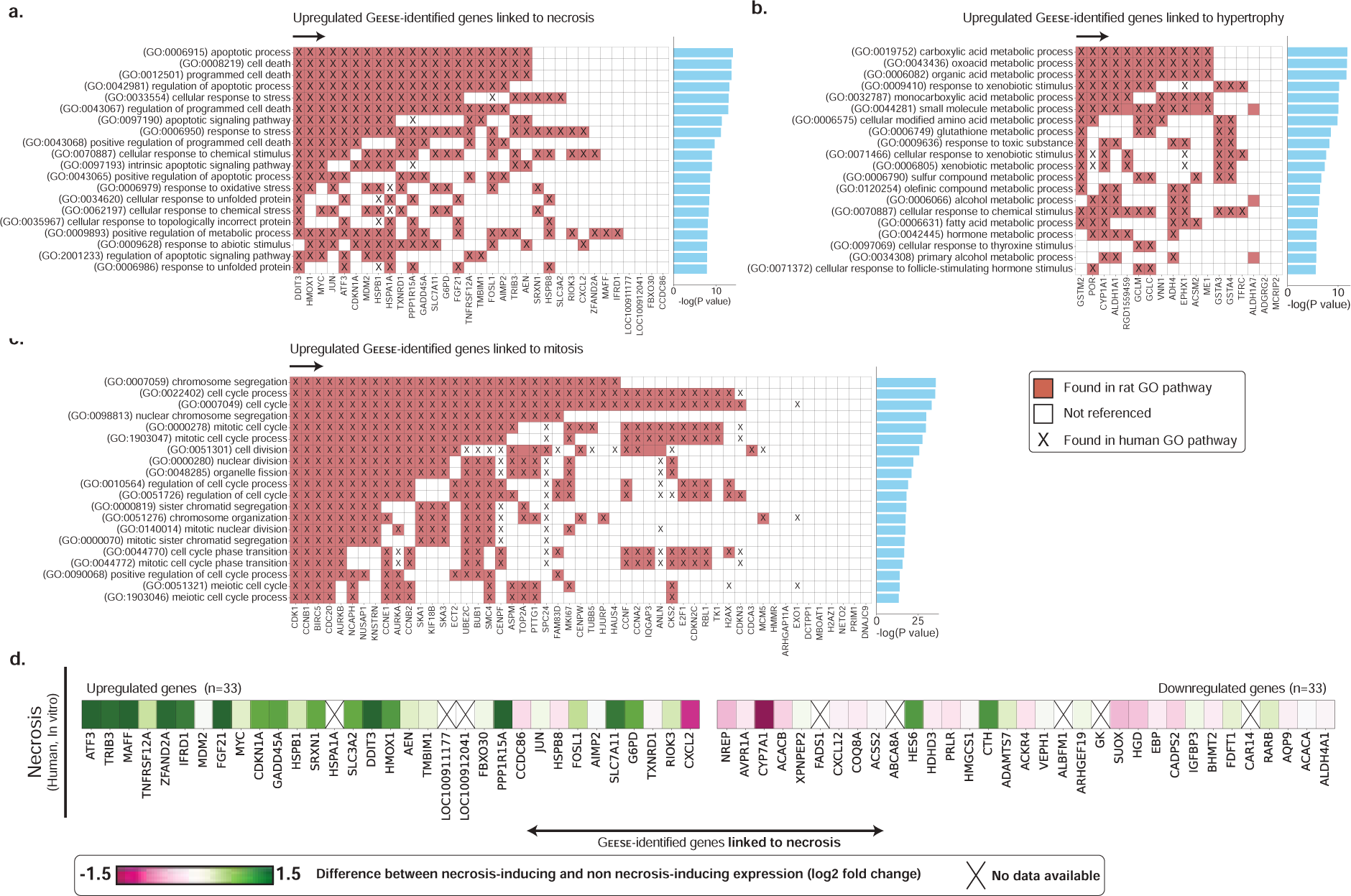
Pathway enrichment analysis and human *in vitro* validation. **a.** Pathway enrichment analysis of GEESE-identified genes uniquely linked to necrosis in rat and human biological processes. **b,c.** Enrichment analysis conducted for hypertrophy and increased mitosis. Additional analysis is provided in **fig. S12,13,14,15,16, and 17**. **d.** Translation to *in vitro* primary human hepatocyte cell lines in the gene set uniquely identified as linked to necrosis (*in vivo*) with a focus on the 33 upregulated and 33 downregulated genes identified as related to necrosis.

### Translation to *in vitro* human cell lines

We further assessed the translatability of the GEESE-identified genes associated with necrosis to human biology. To this end, we leveraged data from *in vitro* primary human hepatocytes (PHH) cell lines collected as part of TG-GATEs on a subset of 140 compounds. For each tested compound, expression changes were measured using mRNA microarrays after fixed time intervals (**Materials and Methods**, section **Gene expression of *in vitro* human studies**). Specifically, from the *in vivo* rat experiments, we defined two groups of compounds: (1) a group with 10 compounds where at least five necrotic slides were found (corresponds to 42 high-dose samples *in vitro*), and (2) a group with 83 compounds without any slides with necrosis, which act as a control group (corresponds to 331 high-dose samples *in vitro*). For each group, we computed the average gene expression (log2 fold change) of the high-dose samples, specifically targeting genes identified by GEESE *in vivo* (rat). We then reported the expression difference between the first and second groups.

Several GEESE-identified genes were associated with necrosis, including *ATF3*, *TRIB3*, and *MAFF*, which were also differentially expressed in PHH treated with necrosis-inducing compounds (**Fig. 5d and fig. S18**). This suggests that these genes could be conserved markers of necrosis across species and experimental systems. These findings align with our pathway enrichment analysis and comparison against the CTD database. Other genes, such as *MDM2* and *AVPR1A*, did not show the same differential expression patterns in PHH as observed in rat livers, which several factors could explain: (1) the doses and time points used in the PHH experiments may not elicit the full extent of necrosis-related gene expression changes observed *in vivo*; (2) PHH cell lines have known limitations in fully recapitulating the complexity of the intact liver, such as its 3D architecture, zonation, and cross-talk with non-parenchymal cells and other organs^62^; and (3) inherent differences between rat and human hepatocytes could result in a lack of direct translation for some genes. Despite the translational gap across species and systems, GEESE successfully identified several genes that exhibit evidence of conservation as biomarkers of liver lesions and could be further investigated.

## Discussion

In this study, we demonstrated that large deep learning models can be used to elucidate morphological correlates of molecular changes in toxicity studies. To this end, we built GEESE, a weakly-supervised regression model (GEESE) trained on over 8,000 liver tissue sections from 129 preclinical safety studies to predict bulk gene expression changes of 1,536 gene targets. In addition to slide-level profiling, GEESE can derive local gene expression changes by predicting pseudo-spatially-resolved gene expression maps of 1,536 genes. By combining GEESE with a morphological lesion classifier that can precisely locate and quantify six commonly found morphological lesions in liver, we extracted from 29 held-out safety studies a dataset of 25 million image patches, each associated with a pseudo-expression profile and lesion labels. This analysis enabled us to identify various morphomolecular associations within and across multiple compounds. For example, the upregulation of 33 genes such as *ATF3*, *TNFRSF12A*, *DDIT3*, known for their involvement in stress response pathways and apoptotic signaling^63^, was associated with the presence of hepatocellular necrosis in multiple studies. Similarly, the presence of mitosis was consistently linked with the upregulation of 52 genes such as *CCNA2*, *CCNB2*, *KNSTRN* and *CDKN3*, which are involved in the regulation of the cell cycle, DNA replication, and mitotic spindle formation. This analysis allowed the curation of comprehensive gene sets associated with each of the six studied lesion types that we further validated against public toxicogenomic databases, pathway enrichment analyses, and *in vitro* human cell line data. Overall, GEESE enables the discovery of subtle and robust morphomolecular associations within and across compounds at an unparalleled scale.

Even though the size of the cohorts used in this study is of unprecedented size in toxicology, our study has limitations. First, our analysis is focused on rat liver tissue, which limits our findings to a single organ. Analyzing other tissues and confirming the conservation of signatures across species would strengthen the translational relevance. While we validate our analysis with *in vitro* primary human hepatocyte cell lines, these data remain limited in representing the complexity of an intact liver. Future studies, which employ more advanced *in vitro* models that better mimic the *in vivo* liver microenvironments such as high-content imaging^64^, 3D spheroids^65^ or organ-on-a-chip systems^66^, can accelerate such translational efforts. Additionally, the number of compounds considered in the downstream analysis is limited to 29 *in vivo* rat studies (156 total, with 127 for model training) and 140 *in vitro* human studies. Therefore, our analysis cannot encompass the morphomolecular diversity that the administration of any compound can induce. Scaling to thousands of preclinical studies (and millions of slides) is needed to increase the diversity of the discovered morphomolecular signatures. Lastly, validating the accuracy of GEESE pseudo-spatially resolved expression profiles remains challenging. Immunohistochemistry (IHC) is not routinely performed in toxicology studies, and spatial transcriptomic (ST) data remains scarce due to high cost.

We envision the incorporation of GEESE into the preclinical workflow, providing a valuable tool for toxicogenomic profiling from histology and imputation of spatially-resolved gene expression maps. In addition, GEESE can be used to automatically identify, quantify, and characterize the relationships between morphological lesions and gene expression changes. These capabilities are crucial as toxicity remains a major cause of drug attrition, with preclinical toxicology studies as a critical threshold for the $1.8 billion total cost necessary to bring a new molecular entity to market^2, 67, 68^. Given the high attrition rate of drug candidates due to toxicity during preclinical testing with only about 5% of compounds that enter preclinical studies ultimately receiving approval^1^, GEESE can contribute to streamlining toxicity assessment by reducing manual semi-quantitative evaluations and providing molecular insights cost-effectively without requiring specialized techniques like ST or IHC.

While additional validations will be required to ascertain some of our findings, our approach to morphomolecular signature discovery can seamlessly scale to more compounds, gene targets, and species. In addition, joint efforts from the pharmaceutical industry and academia, such as the BigPicture initiative^69^, will gather large cohorts of preclinical studies, which can be harnessed as additional training data or validation. Furthermore, the integration of additional clinical data, such as from the DrugMatrix^10^ and ToxCast^70^ programs, could help bridge the translational gap between rodent studies and human toxicity. Finally, establishing community-wide standards and infrastructure for structuring toxicological data for AI method incorporation and foundation model development, such as through the eTRANSAFE^71^ consortia, will bring us closer to translating AI methods into practical tools for drug and biomarker discovery. Overall, our study lays the foundations for several promising avenues in AI-driven toxicology research and preclinical drug safety assessment.

## Materials and Methods

### Ethics statement

The study involves a retrospective examination of previously collected tissue samples of *Rattus Norvegicus* liver sections, which are part of a public archive. Examination of the original study’s documentation confirms that the experimental protocol was subject to an ethical review and subsequently received approval from both the Ethics Review Committee for Animal Experimentation at the National Institute of Health Sciences (NIHS) and the relevant contract research organizations.

### Study design

#### TG-GATEs protocol

Four contract research organizations conducted animal experiments on male Crl:CD Sprague-Dawley (SD). Animals were allocated into groups of 20, each using a computerized stratified random grouping method based on body weight^34^. Two types of dose administration were conducted: single-dose and repeated-dose. In single-dose experiments, groups of 20 animals were administered a compound, and then five animals were sacrificed 3, 6, 9, and 24 hours after administration. In repeated-dose experiments, groups of 20 animals received a dose every day, and five animals were sacrificed 4, 8, 15, and 29 days after administration. For each sample group (unique compound, dose, sacrifice time), three animals underwent a toxicogenomic analysis with mRNA microarrays. Animals were not fasted before being sacrificed. The compounds examined (as detailed in **table S1**) were chosen through literature reviews and agreement among toxicologists from the pharmaceutical industry and the Japanese government. In most compounds, three dose levels were tested with a dose ratio between the low, middle, and high levels of 1:3:10.

#### Histopathology acquisition and annotation

All liver sections were stained with H&E (hematoxylin and eosin) and mounted on glass slides. Tissue sections were converted into digital pathology images using a ScanScope AT scanner (Aperio Technologies Inc., CA, USA) at 20*×*magnification (0.49 *µ*m/px). The histopathology data from TG-GATEs include annotations that detail the lesions observed in the slides. These annotations are unnormalized, with various studies employing differing terminologies and taxonomies to describe identical findings. In total, 66 different lesion types were identified across 23,136 liver sections. However, many of these are either synonyms or more specific classifications of broader lesion categories. In our analysis, we grouped related lesions into six lesions of interest: increased mitosis, necrosis, cellular infiltration, bile duct proliferation, fatty change, and hypertrophy. A description of each lesion is provided in **table S4**.

#### Gene expression of in vivo rat studies

The raw transcriptomic data consists of microarrays (Affymetrix Rat Genome 230 2.0 Array GeneChip) with 31,042 probes. All data followed probe-wise normalization using log2 fold change with respect to a control group. Log2 fold change quantifies the proportional difference, on a logarithmic scale, between the expression levels of a particular probe under two conditions: a control group (on average 22 slides per study in TG-GATEs) and a sample group (a defined set of compound, time and sacrifice). The log2 fold change gene expression changes were not further normalized before processing by our models. Each probe was mapped to a unique gene name identifier, resulting in 13,404 gene expression measurements per sample. From there, we reduced the number of genes analyzed to (1) discard genes unrelated to liver metabolism, drug administration, and toxicity, (2) simplify training of the gene expression prediction model (GEESE), and (3) simplify the post-hoc analysis. Here, we selected genes based on two strategies to ensure the use of a biologically diverse set. Firstly, we included the T1000 gene set^36^, a set of 1,000 genes responsive to chemical exposures from which we retrieved 867 genes. Second, we used a data-driven approach, where we computed the Pearson correlation between each measured gene expression and slide-level lesion labels (as reported in TG-GATEs annotations) in the train studies. We then retained genes with a Pearson correlation larger than a threshold set to 0.15. The threshold was decided arbitrarily to include promising genes that may be morphologically expressed while keeping the total number of genes analyzed around 1,500. The integration of genes from distinct methodologies results in a consolidated subset of 1,536 gene targets.

#### Gene expression of in vitro human studies

In-vitro human experiments were conducted using the Affymetrix human U133 Plus assay with 54,613 probes on primary human hepatocytes (PHH) cell lines^34, 72^. This assay was conducted on a subset of 140 compounds with three dose levels: low, medium, and high, followed by sample collection after 2h, 8h, and 24h.

#### Dataset split

To avoid compound-specific information leakage when training the gene expression regressor, we extracted 29 studies for testing (N=2,002 slides) and kept 127 studies from 8,232 pairs for training and validation. We further split training and validation slides to obtain a train (N=6,585 slides) and validation set (N=1,647 slides). From the 6,585 samples in the training set, 1,141 (17%) were annotated as containing one or multiple lesions: 154 with increased mitosis (2.3%), 314 with necrosis (4.8%), 321 with cellular infiltration (4.9%), 54 with bile duct proliferation (0.8%), 143 with fatty change (2.2%), and 526 with hypertrophy (8.0%). From the 2,002 samples from the testing set, 364 samples were annotated as containing one or multiple lesions: 53 with increased mitosis (2.6%), 150 with necrosis (7.5%), 116 with cellular infiltration (5.7%), 21 with bile duct proliferation (1.0%), 47 with fatty change (2.3%), and 166 with hypertrophy (8.3%). The complete distribution of lesions in test studies is provided in **table S6**.

### Deep learning modeling

All slides are preprocessed by first segmenting tissue regions and then tesselating the slide into patches (see **Tissue segmentation and patching**). Then, we learn a vision encoder based on self-supervised learning to derive a compressed representation of image patches (see **iBOT pre-training**). The learned patch embeddings serve as input to the expression regressor (GEESE) following the Multiple Instance Learning (MIL) paradigm^38, 39^ (see **Gene expression regressor**). In addition, patch-level annotations are used to fine-tune the patch encoder and extract pseudo-lesion labels (see **Lesion classifier**).

#### Tissue segmentation and patching

Before MIL training, each slide was segmented using the CLAM toolbox^39^ that includes modules for automatic detection of tissue vs. background. After segmentation, non-overlapping 256 *×* 256-pixel patches were extracted at 20*×* magnification (0.5 *µ*m/px) and then resized to 224*×*244 pixels image patches.

#### iBOT pretraining

We employed the iBOT framework^46^, a state-of-the-art approach in self-supervised learning for building compressed morphological descriptors (patch embeddings) of image patches. iBOT employs a student-teacher knowledge distillation strategy designed for pretraining Vision Transformer (ViT)^45^. iBOT uses two main objectives: self-distillation loss^47, 73^, which aims to align the representations of a student and teacher network, and masked image modeling loss^74^, which aims to reconstruct the original image from partially observed inputs. We trained a ViT-Base model that yields 768-dimensional embeddings on 15 million patches extracted from 46,734 WSIs. We trained the network for 1,176,640 iterations (or 80 epochs). The specific hyperparameters used for training are listed in **table S15**.

#### Gene expression regressor (GEESE)

We cast gene expression prediction as a weakly supervised regression task, where we learn a pooling function to aggregate the iBOT patch embeddings into a slide-level gene expression prediction. We propose the gene expression regressor, denoted as GEESE, that enables joint derivation of patch and slide prediction scores using slide-level supervision only. Namely, each patch embedding, denoted as 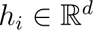, is passed to a patch regressor network *f*(.). Then, the slide-level regressor is built by taking the arithmetic mean over all patch-level regression scores, resulting in a slide prediction. Formally, we define it as:

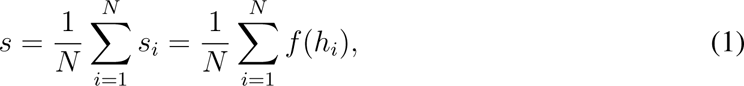

where 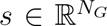 denotes the slide-level log2 fold change gene expression scores. As the slide prediction is directly defined as the mean of the individual patch contributions, denoted as 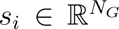, the gene-wise patch importance can be readily obtained without the need for analyzing attention scores^39^ or gradient attribution^33, 75^. The resulting patch attribution *s_i_* can, therefore, be seen as a pseudo-spatially-resolved gene expression, where the resolution is given by the patch resolution (for the patch of 256*×*256 pixels at 0.5*µ*m/px, the resolution is 128*µ*m).

Here, *f*(.) is implemented using a 4-layer MLP patch regressor with LayerNorm, dropout (0.1) between all layers, and GELU activation (see **table S16**). This formulation has connections with AdditiveMIL^76^.

#### Lesion classifier

We use the iBOT model as a foundation for classifying six common liver lesions at patch-level (each patch is 256*×*256 pixels or 128*µ*m). As the TG-GATEs cohort only includes slide-level labels, we curated a set of patch annotations. To this end, we employed four different approaches: 1) **Public annotations** We used publicly available annotations provided by Bayer Pharmaceuticals and Aignostics GmbH https://zenodo.org/record/7541930. These annotations consist of polygonal annotations within 230 whole slide images from TG-GATEs. These polygonal annotations were subsequently converted into patch annotations, with patches retained based on their overlap with annotations. 2) **Human-in-the-loop annotation** Semi-annotated human-in-the-loop annotations were generated using a weakly supervised slide classification system. A subsequent manual review led to the selection of true positive examples. 3) **Normal patches** To include normal patches, we extracted ten random patches from lesion-free slides, each thoroughly examined to exclude small lesions such as mitosis or single-cell necrosis. 4) **Manual annotation** Human annotations were performed using the QuPath software^77^ to extract missing lesions such as fatty change. The process yielded 24,631 patch annotations with lesions extracted from 3,458 slides, and 13,888 normal patch annotations from 3,531 slides (see **table S3**).

The pretrained iBOT vision encoder was fine-tuned on these annotations and trained to minimize classification loss, defined as a multilabel binary cross entropy objective on all six classes. Note that each patch can either be normal (no lesion detected) or include one or multiple lesions. Here, we used a class-stratified 80/20% train/validation split. The network was finetuned for 20 epochs using the AdamW optimizer with an initial learning rate of 4e-4 and layerwise learning decay of 0.65. Basic patch augmentations were performed during fine-tuning, based on random color jittering, mirroring, and rotation. The lesion classifier provides the likelihood that each patch contains one of the six lesions (expressed as a probability post-Sigmoid activation). To ensure that we only include positive patches, we use conservative classification thresholds set to 0.95 for cellular infiltration, 0.9 for necrosis, 0.9 for bile duct proliferation, 0.99 for fatty change, and 0.9 for increased mitosis. Each threshold was determined using an independent set of patches for each lesion.

### Post-hoc morphomolecular signature analysis

Using GEESE, we inferred the patch-level pseudo-expression on the 2,002 slides from TG-GATEs test set. We proceeded analogously to extract patch-level lesion prediction using our morphological lesion classifier. Each patch becomes assigned to 1,536 gene expression scores (expressed as a log2 fold change), and a single or multiple morphological labels, such as necrosis, cellular infiltration and fatty change, or normal if no lesion was detected. In total, this operation yielded 25 million lesion-expression pairs. We used all pairs for the downstream quantitative analyses.

#### Gene identification

We present the detailed steps for the post-hoc analysis of necrosis. A similar process is conducted for all other five lesions. We start by selecting compounds associated with necrosis (for instance, we selected 4 out of 29 studies from the TG-GATEs test set, see **table S5**). Out of the 25 million lesion-expression pairs, we subsequently selected patches that contain necrosis in the selected studies, yielding 53,542 patches (**fig. S13**). Using the corresponding patch-level expression of the selected patches, we extract the most upregulated and the most downregulated genes out of the 1,536 gene targets. Specifically, we compute the average patch expression per selected compound and further average across all selected compounds. We then extract genes with a mean expression above 1 log2 fold change (upregulated genes), and with a mean expression below −1 (downregulated genes). We proceed similarly for other lesions using a threshold of −0.5 and 0.5 (instead of 1 and −1) for increased mitosis, fatty change, cellular infiltration, and hypertrophy. These thresholds were set arbitrarily so that a pool of genes could be identified for further analysis and validation. Varying this threshold controls the number of genes used for additional investigation.

We refine the gene selection as a final step to focus on genes that are only differentially expressed for a single lesion (e.g., *CCNA2* is only upregulated in the presence of mitosis, whereas *ABCC3* is associated with multiple lesions, such as necrosis, cellular infiltration, and fatty change). A gene is selected as specific to necrosis if it satisfies the following three conditions simultaneously: (1) no other lesion is more differentially expressed than necrosis, (2) its absolute average expression across the five other lesions is at least two times lower, and (3) the absolute Pearson correlation between measured and predicted slide-level gene expressions, for slides identified as containing necrosis, is above 0.3. Formally, we express these three conditions as,

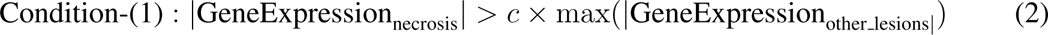

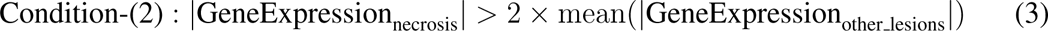

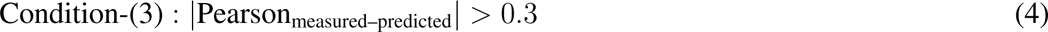

where the constant *c* is set to 1.5. Intuitively, Condition-(1) screens an initial subset of genes associated with necrosis. Condition-(2) serves to refine this subset, where the constant *c* is set arbitrarily to 2 to identify genes with the largest correlation to necrosis while removing those primarily indicative of generic toxic exposure effects. Condition-(3) ensures that only the genes that can be satisfactorily predicted from tissue morphology, with GEESE, are retained. We follow the same process for other lesions besides cellular infiltration, where the first condition is relaxed such that no other lesion is more differentially expressed than 75% of its value (c=0.75). This relaxation reflects the fact that cellular infiltration often co-appears with necrotic patches and bile duct proliferation. For the same reason, in the analysis of bile duct proliferation and necrosis, we exclude cellular infiltration from Condition-(1) and Condition-(2).

#### Pathway analysis

To validate our findings, we identify biological pathways related to previously identified genes. The goal of this analysis is two-fold: (1) confirm the biological relevance of the identified molecular signatures and (2) identify new biomarkers that have previously been poorly explored and characterized. To this end, we utilized the Rat Genome Database^11, 55^, a state-of-the-art public resource for multi-species pathway enrichment analysis that includes both rats and human pathways. Specifically, we employed the Multi-Ontology Enrichment Tool, MOET, available within the Rat Genome Database to identify the most relevant biological processes based on the Gene Ontology database. For this analysis, overlaps were computed on the child terms of the term *biological process* (GO:0008150), which include 20,292 process sets for rats and 19,761 process sets for humans. This allowed the identification of biological pathways in which the genes from each lesion’s gene list were most involved.

### Evaluation and implementation

#### Training details

All weakly supervised expression regression models are trained using the AdamW optimizer with an initial learning rate of 1e-04, a mean-squared error objective, a maximum of 40 epochs with early stopping (patience set to 10) with respect to the validation loss.

#### Metrics

GEESE predictive performance is evaluated using the Pearson correlation, Area under the ROC Curve (AUC), log2 fold change, R2, and Mean Squared Error.

**Pearson correlation** describes the linear relationship between two sets of scalars. It varies between −1 and +1, with 0 implying no correlation. The corresponding p-value (2-tailed p-value) represents the two-sided null hypothesis of non-correlation. We employ the pearsonr implementation from the Python package Scipy version 1.13.0.

**AUC** is the area under the receiver operating curve plotting the true positive rate against the false positive rate as the classification threshold is varied. This metric is mainly used for classification tasks but can be adapted for regression. Formally, we define AUC for regression as,

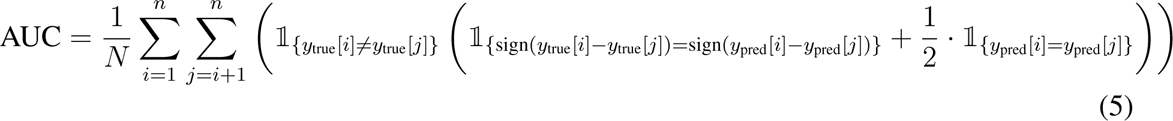

where 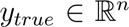 and 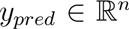 are vectors with the true and predicted regression values, *N* is the number of valid pairs (i,j).

**Log2 fold change** is a measurement commonly used to quantify the relative change between two experimental conditions. It is calculated by taking the base 2 logarithm of the ratio between the percentage of a certain lesion under some conditions (such as high dose, sacrifice time of 29 days) and the percentage of that same lesion in the control group.

**R2** is a metric used to assess the quality of a regression model, where 1 indicates perfect regression, and 0 random regression. R2 measures the goodness of the fit and represents the proportion of variance in the dependent variable that is explained by the model. We employ the metrics.r2 score implementation from the python package scikit-learn version 1.2.1.

**Mean Squared Error (Standardized)** is computing the average of the squares of the errors and is used to quantify how far model predictions are from the actual values. Before computing the mean square error, the gene expression measured and predicted was standardized (subtracted by the mean and divided by the standard deviation of the gene expression measured). This standardization is made to make this metric consistent across genes. We employ the metrics.mean squared error implementation from the python package scikit-learn version 1.2.1.

#### Statistical analysis

The reported error bars correspond to 95% confidence intervals derived using non-parametric bootstrapping using 100 bootstrap iterations.

### Computing hardware and software

In this study, all coding was conducted using Python version 3.9. The neural networks were implemented with PyTorch version 2.1.0 with CUDA version 11.7. For whole slide image (WSI) pre-processing and manipulation, we utilized OpenSlide version 4.3.1 and openslidepython version 1.2.0. Metrics were implemented using Scikit-learn version 1.2.1. and Scipy version 1.13.0 Data processing tasks were performed using Pandas version 1.4.2, Numpy version 1.21.5, Pillow version 9.3.0 and OpenCV-python version 3.3.1. Matplotlib version 3.7 was employed for generating plots. The training of the patch encoder was based on the original iBOT implementation, which is available at ^1^. The pretraining of iBOT was carried out on 8 *×* 80GB NVIDIA A100 GPUs, configured for multi-GPU training using distributed data parallelism. Downstream experiments were conducted on 3 *×* 24GB NVIDIA 3090 GPUs. Slide annotation and visualization were done using QuPath version 0.4.3. Finally, rat microarray probes were converted using SynGoPortal, accessible at ^2^, and human microarray probes were converted using the python API of PythonBio, version 1.83. The viewer used for the online demo is based on OpenSeadragon (version 4.1.0) and JavaScript (version ES13). The GO processes for rats and humans were queried using the MOET tool (Multi Ontology Enrichment Tool) from the Rat Genome Database accessible at ^3^. Genes linked to chemical and drug-induced liver injury were retrieved using the Comparative Toxicology Database accessible at ^4^.

## Data availability

The TG-GATEs data, which includes histopathology whole-slide images and labels, is openly accessible on the National Institute of Biomedical Innovation portal at ^5^. A subset of 230 TG-GATEs with pixel annotations can be freely accessed from Zenodo at ^6^. Patch annotations, as well as pseudo-patch annotations generated by the fine-tuned patch encoder, are available on a case-by-case basis, depending on specific needs. The microarray data, part of The Japanese Toxicogenomics Project, were obtained from the Toxigates portal, accessible at ^7^.

## Code availability

Upon publication, the authors will release code and pre-trained models for extracting patch-level embeddings and lesion classification, performing weakly-supervised gene expression regression, and analyzing gene expression predictions and lesion predictions.

## Author contributions

G.J and T.P conceived the study and designed the experiments. G.J, T.P and L.O performed data collection and data cleaning. G.J, T.P, A.S, L.O, R.P, D.W, R.J.C, A.V performed model development and experimental analysis. G.J, T.P, A.S, R.P, D.W, S.B, J.T, L.L, and G.G. interpreted experimental results and provided feedback on the study. G.J, T.P, A.S, R.P prepared the manuscript with input from all co-authors. F.M. supervised the research.

## Acknowledgements

This work was supported in part by BWH & MGH Pathology, BWH President’s Fund, Massachusetts Life Sciences Center, NIGMS R35GM138216 (F.M.), and BWH President’s Scholar fund (G.G.) and NIGMS R35GM149270 (G.G.). R.J.C. was also supported by the NSF Graduate Fellowship. L.O. was supported by the German Academic Exchange (DAAD) Fellowship. We thank Dr. Pierre Moulin for the early-stage discussions on toxicity assessment and computational toxicology.

---

**Figure S1:**
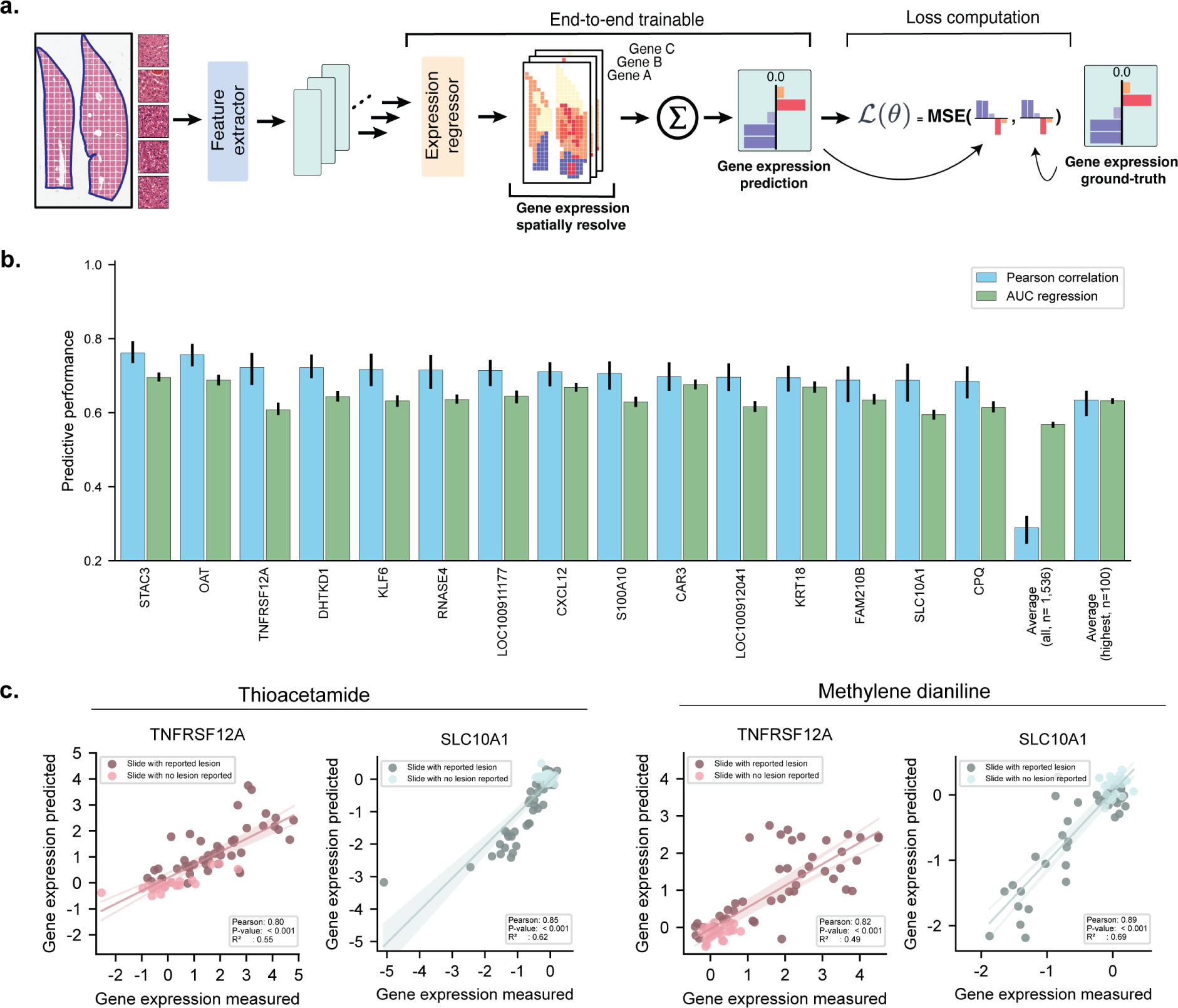
GEESE model architecture and performance. **a.** Overview of the gene expression regressor (GEESE) architecture, which can predict expression changes (log2 fold change) from an input tissue section (a whole-slide image). Patch attribution enables deriving a pseudo-spatially-resolve expression map, where each patch becomes associated with a pseudo-expression profile for all predicted genes (n=1,536). **b.** Slide-level gene expression prediction performance evaluated using Pearson correlation and macro-AUC regression on TG-GATEs test set (N=2,002 slides). We report predictive performance for the top 10 predicted genes, the average predictive performance across all genes (n=1,536), and the top 100 best-predicted. Error bars represent 95% confidence intervals using non-parametric bootstrapping (100 iterations). **c.** Expression prediction of *TNFRSF12A* and *SLC10A1* genes on compounds thioacetamide and methylene dianiline. Each dot represents a sample. Samples without lesions are clustered around the origin (small log2 fold change), while samples with lesions are either downregulated (*SLC10A1*) or upregulated (*TNFRSF12A*). P-value derived from testing the two-sided null hypothesis of non-correlation.

**Figure S2:**
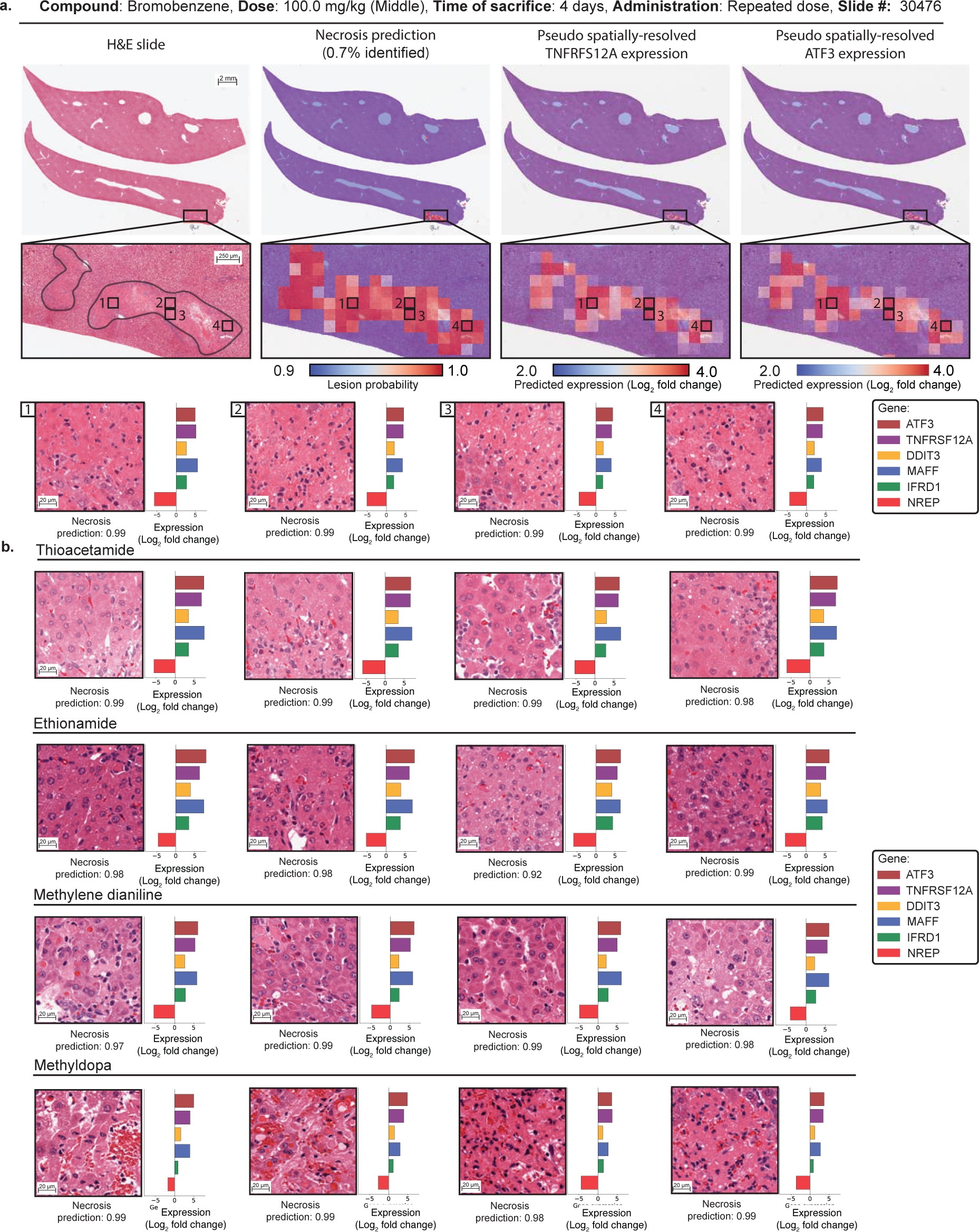
Pseudo spatially-resolved gene expression and visualization of necrosis. **a.** H&E liver section after exposure to bromobenzene (left). Overlay of patch-level necrosis prediction. Predictions below 90% confidence are represented in blue and high-probability predictions are represented in red (center-left). Pseudo-spatially-resolved gene expression heatmaps of genes *TNFRSF12A* (center-right) and *ATF3* (right). **b.** Additional examples of necrotic patches from four studies (thioacetamide, ethionamide, methylene dianiline, and methyldopa) highlight similar upregulation of *TNFRSF12A*, *ATF3*, *DDIT3*, *MAFF* and *IFRD1*, and downregulation of *NREP*.

**Figure S3:**
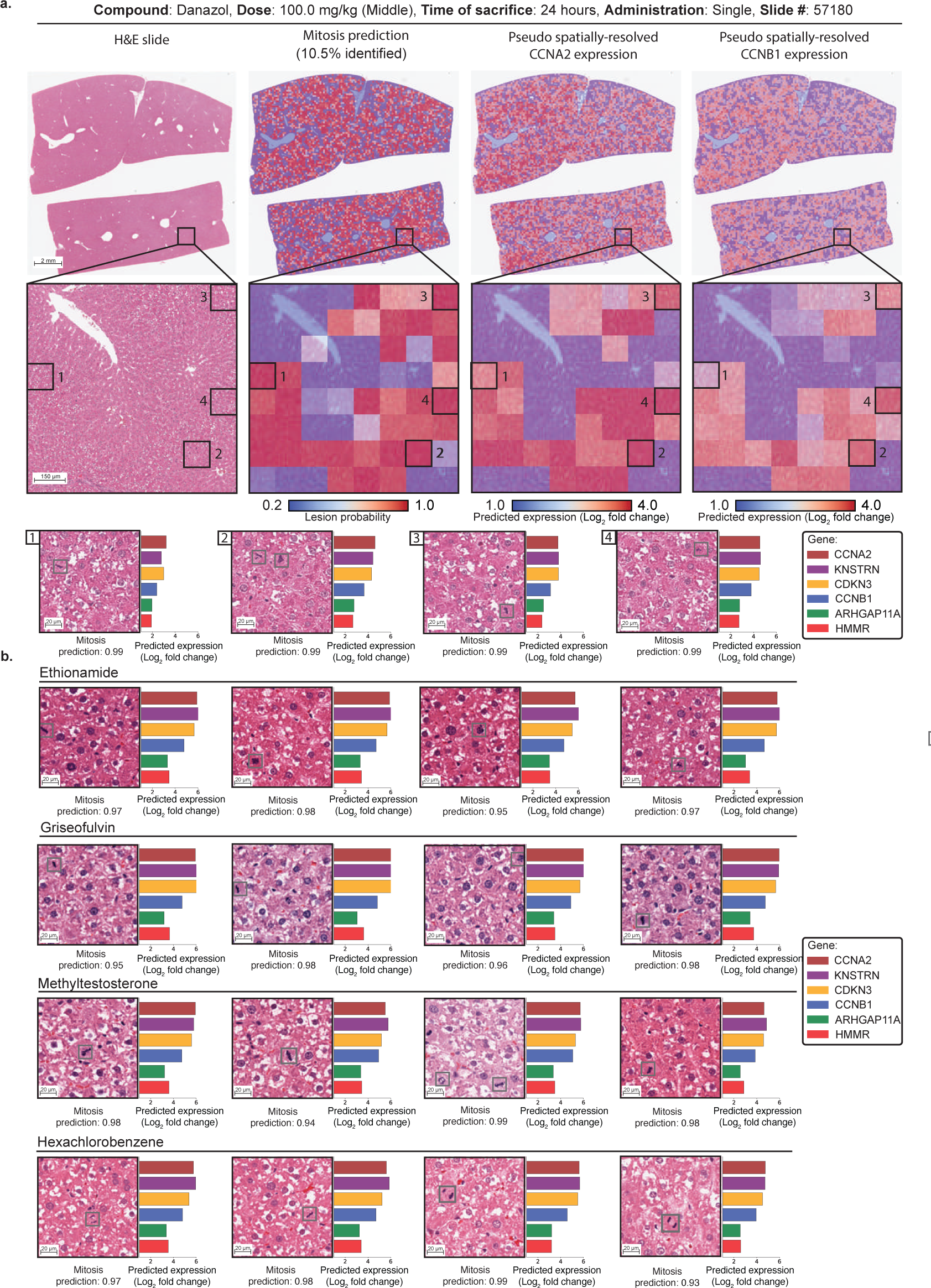
Pseudo spatially-resolved expression and visualization of mitosis. **a.** H&E liver section after exposure to danazol (left). Overlay of patch-level bile duct proliferation prediction. Predictions below 20% confidence are represented in blue, and high-probability predictions are represented in red (center-left). Pseudo-spatially-resolved gene expression heatmaps of *CCNA2* and *CCNB1* (center-right and right). **b.** Additional examples of patches with mitosis from four studies (ethionamide, griseofulvin, methyltestosterone, and hexachlorobenzene) highlight similar upregulation of *CCNA2* and *CCNB1*, *CDKN3*, *KNSTRN*, *ARHGAP11A* and *HMMR*.

**Figure S4:**
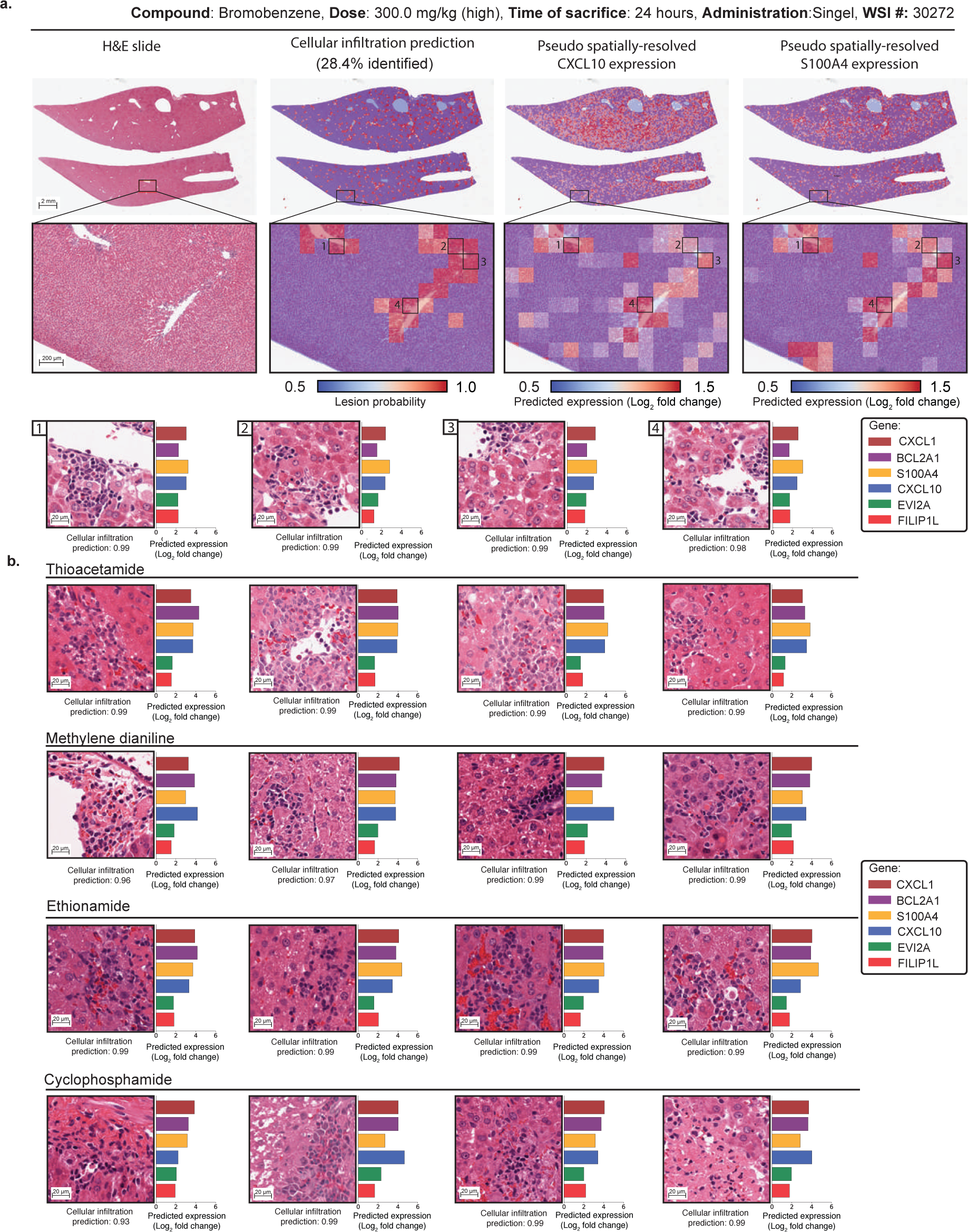
Pseudo spatially-resolved expression and visualization of cellular infiltration. **a.** H&E liver section after exposure to bromobenzene (left). Overlay of patch-level cellular infiltration prediction. Predictions below 50% confidence are represented in blue and high-probability predictions are represented in red (center-left). Pseudo-spatially-resolved gene expression heatmaps of *CXCL10* and *S100A4* (center-right and right). **b.** Additional examples of patches with cellular infiltration from four other studies (thioacetamide, methylene dianiline, ethionamide, and cyclophosphamide) highlight similar upregulation of *CXCL10*, *S100A4*, *CXCL1*, *BCL2A1*, *EVI2A* and *FILIP1L*.

**Figure S5:**
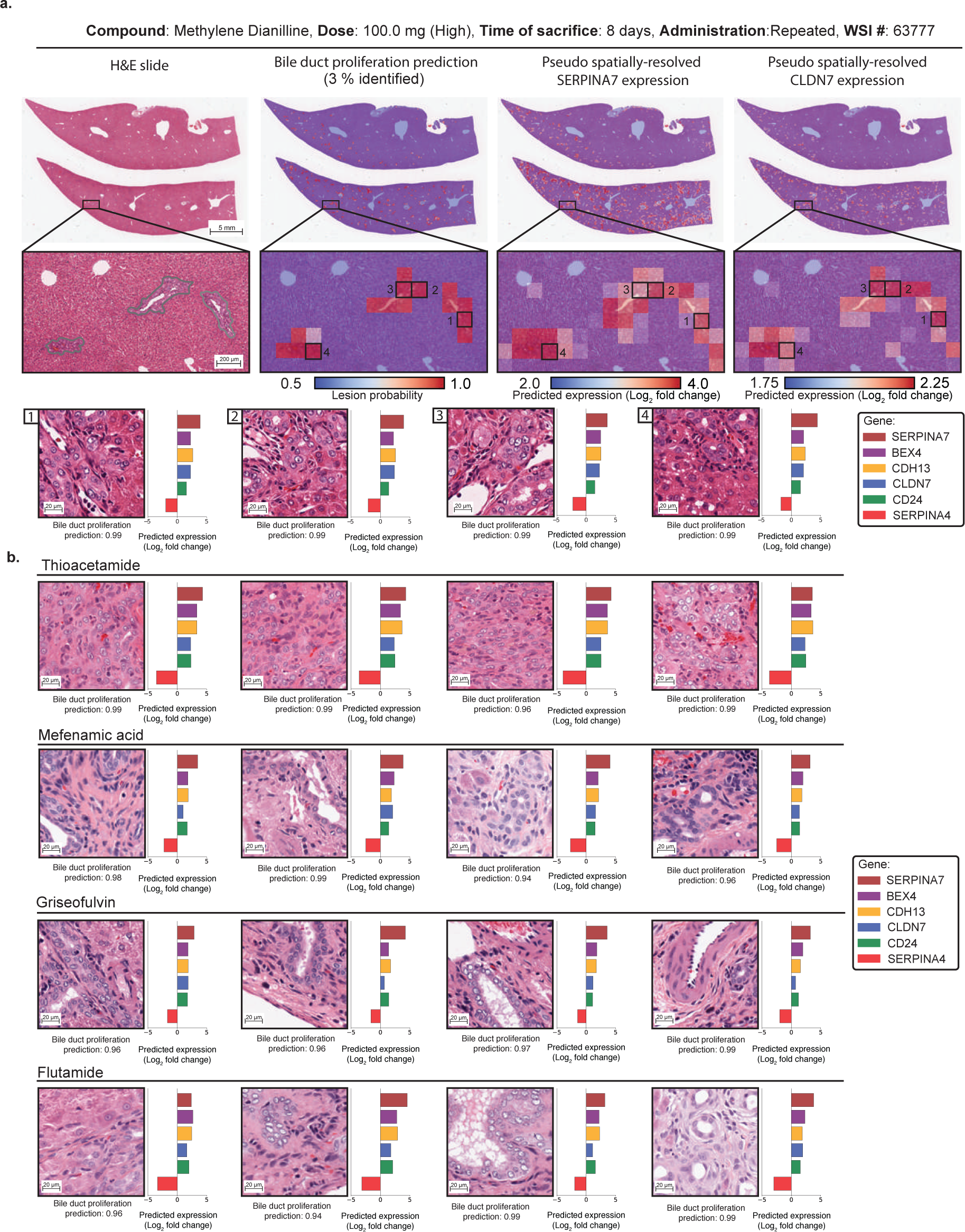
Pseudo spatially-resolved expression and visualization of bile duct proliferation. **a.** H&E liver section after exposure to methylene dianiline (left). Overlay of patch-level bile duct proliferation prediction. Predictions below 50% confidence are represented in blue and high-probability predictions are represented in red (center-left). Pseudo-spatially-resolved gene expression heatmaps of *SERPINA7* and *CLDN7* (center-right and right). **b.** Additional examples of patches with bile duct proliferation from four other studies (thioacetamide, mefenamic acid, griseofulvin and flutamide) highlight similar upregulation of *SERPINA7*, *CLDN7*, *BEX4*, *CDH13* and *CD24*, and downregulation of *SERPINA4*.

**Figure S6:**
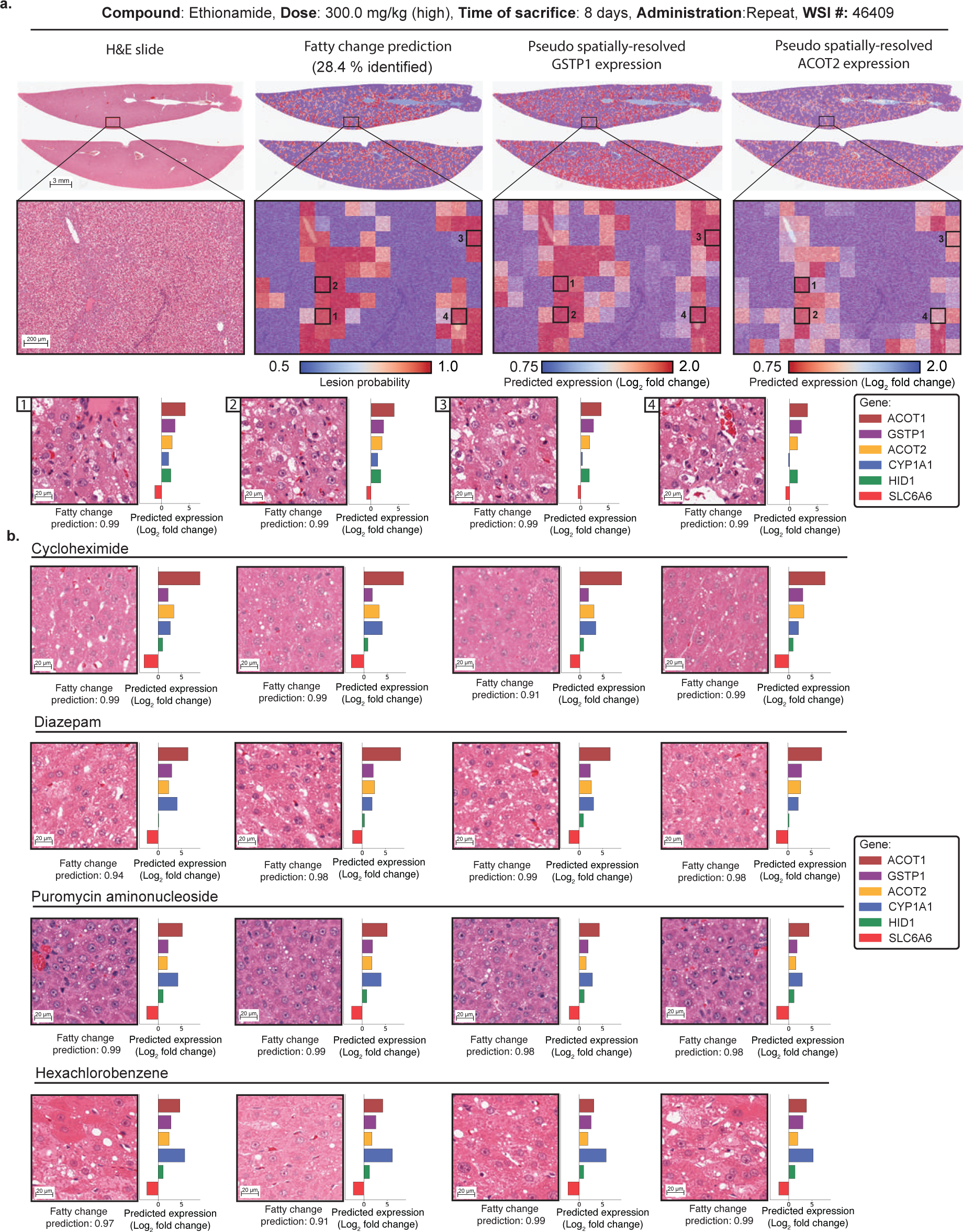
Pseudo spatially-resolved expression and visualization of fatty change. **a.** H&E liver section after exposure to ethionamide (left). Overlay of patch-level fatty change prediction. Predictions below 50% confidence are represented in blue and high-probability predictions are represented in red (center-left). Pseudo-spatially-resolved gene expression heatmaps of *GSTP1* and *ACOT2* (center-right and right). **b.** Additional examples of patches with fatty change from four other studies (cycloheximide, diazepam, puromycin aminonucleoside and hexachlorobenzene) highlight similar upregulation of *GSTP1*, *ACOT2*, *ACOT1*, *HID1* and *CYP1A1* and down-regulation of *SLC6A6*.

**Figure S7:**
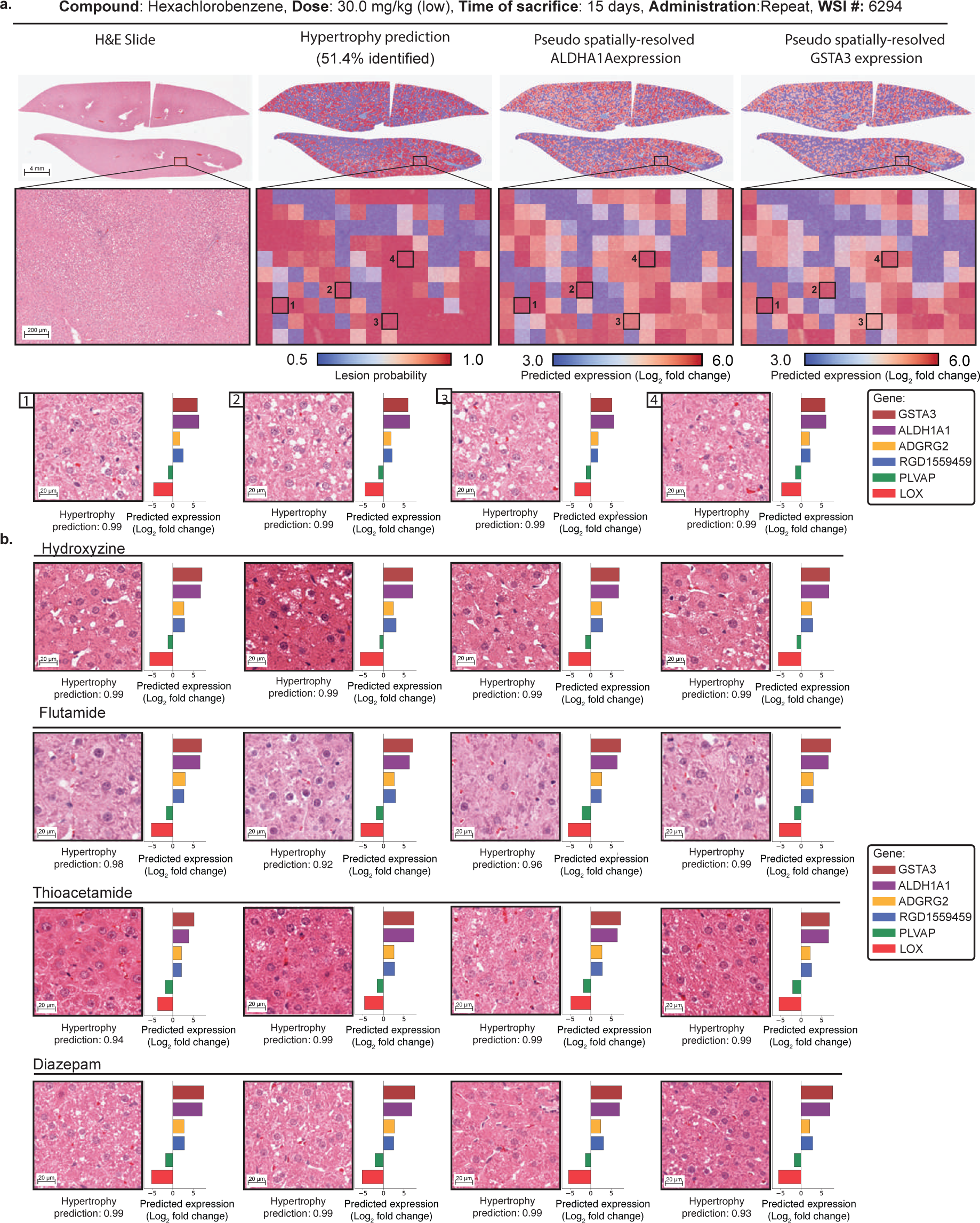
Pseudo spatially-resolved expression and visualization of hypertrophy. **a.** H&E liver section after exposure to hexachlorobenzene (left). Overlay of patch-level hypertrophy prediction. Predictions below 50% confidence are represented in blue and high-probability predictions are represented in red (center-left). Pseudo-spatially-resolved gene expression heatmaps of ALDHA1A and *GSTA3* (center-right and right). **b.** Additional examples of patches with hypertrophy from four other studies (hydroxyzine, flutamide, thioacetamide, and diazepam) highlight similar upregulation of *GSTA3*, *ALDH1A1*, *ADRG2* and *RGD1559459*, and downregulation of *PLAVP* and *LOX*.

**Figure S8:**
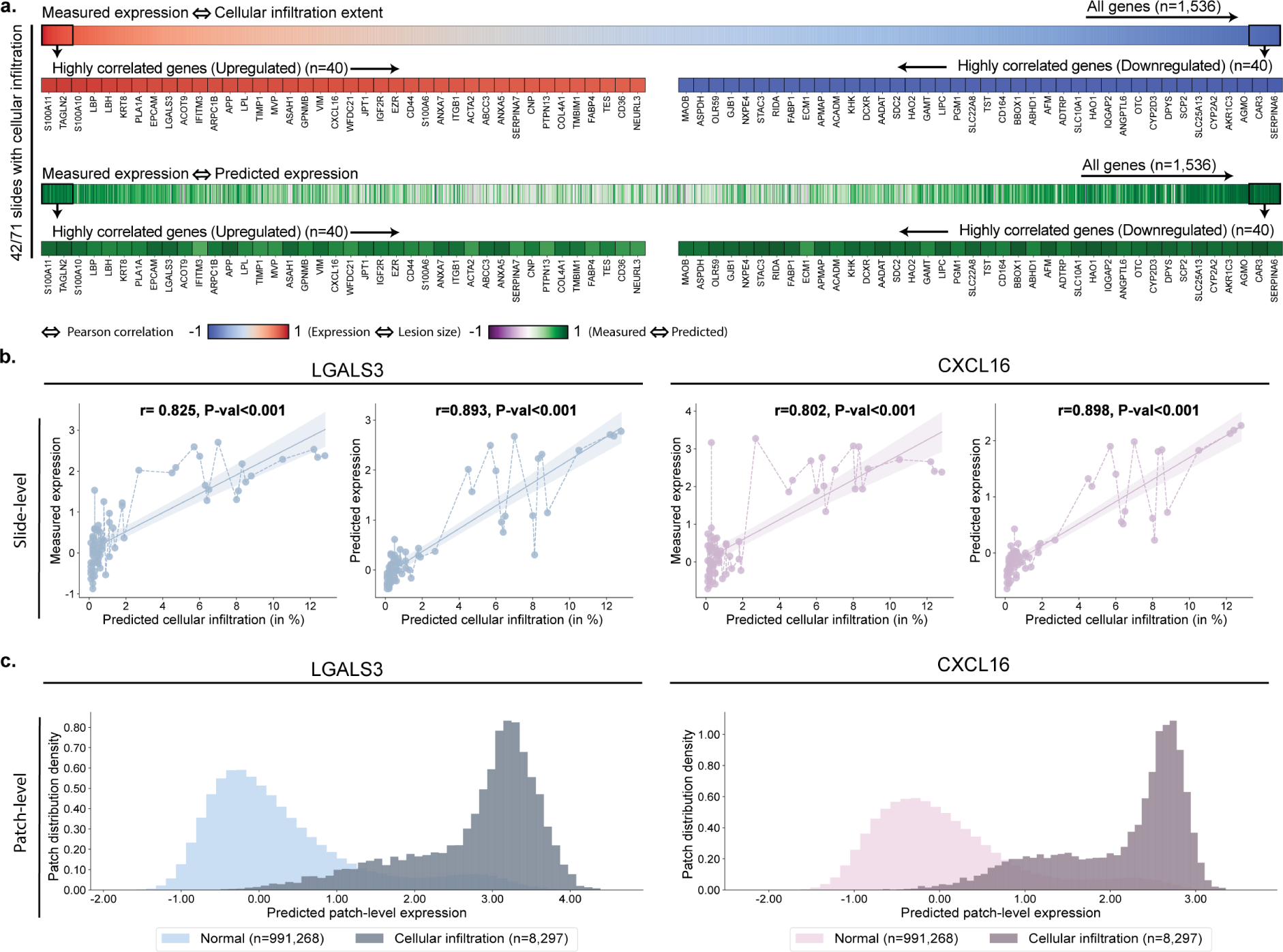
Morphomolecular analysis of cellular infiltration in methylene dianiline. **a.** For each slide-expression pair, we correlate the percentage of cellular infiltration predicted by the lesion classifier with the measured gene expression. Genes with a high correlation between measured expression and lesion size define morphomolecular signatures associated with the compound. **b.** Correlation between the estimated percentage of the slide containing cellular infiltration and the gene expression of *LGALS3* and *CXCL16* (measured with microarray and predicted with GEESE). P-value derived from testing the two-sided null hypothesis of non-correlation. **c.** Distribution of patch-level expression for patches predicted as normal (n=991,268) and patches predicted as containing cellular infiltration (n=8,297).

**Figure S9:**
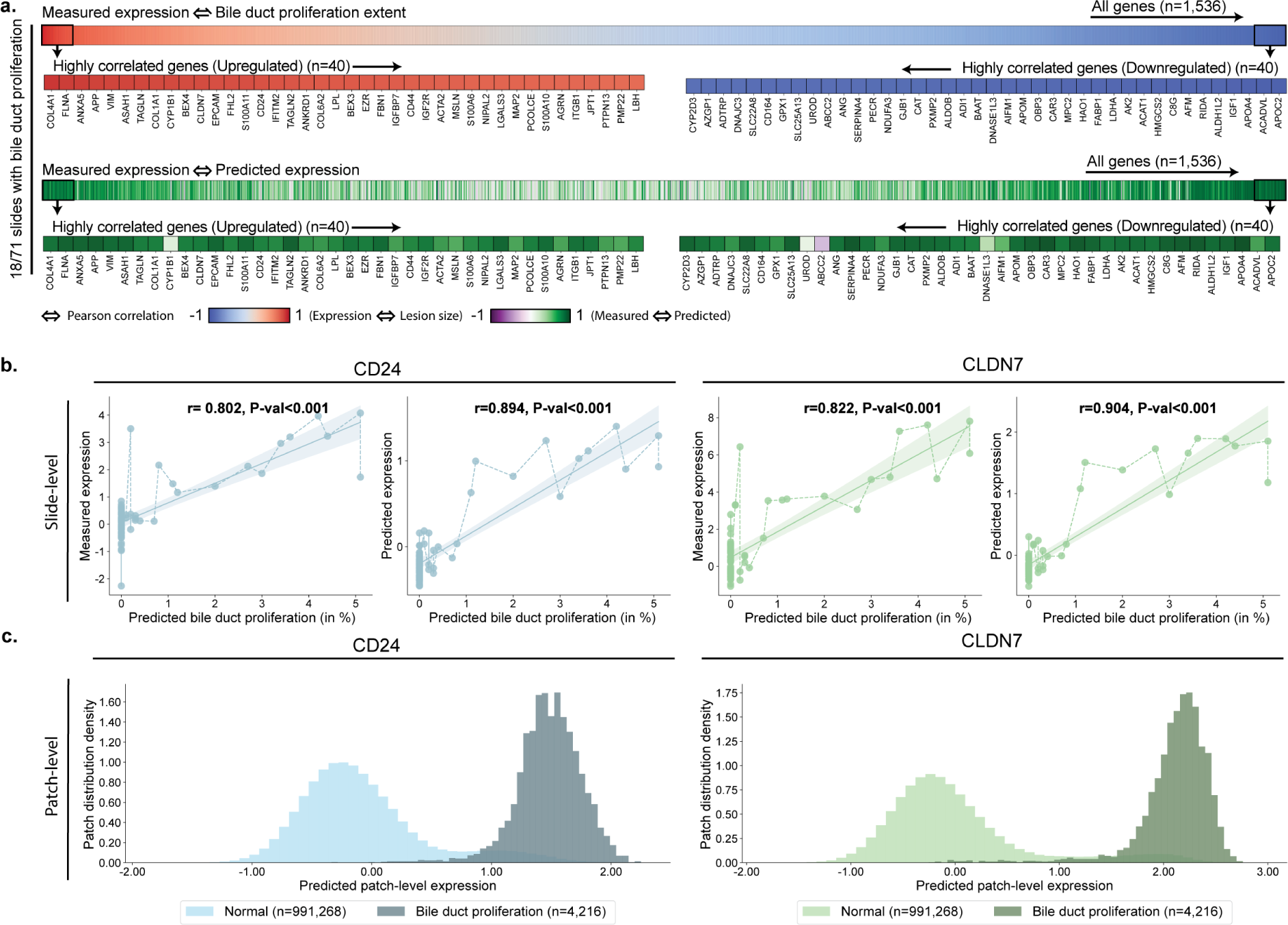
Morphomolecular analysis of bile duct proliferation detection in methylene dianiline. **a.** For each slide-expression pair, we correlate the percentage of bile duct proliferation predicted by the lesion classifier with the measured gene expression. Genes with a high correlation between measured expression and lesion size define morphomolecular signatures associated with the compound. **b.** Correlation between the estimated percentage of the slide containing bile duct proliferation and the gene expression of *CD24* and *CLDN7* (measured with microarray and predicted with GEESE). P-value derived from testing the two-sided null hypothesis of non-correlation. **c.** Distribution of patch-level expression for patches predicted as normal (n=991,268) and patches predicted as containing bile duct proliferation (n=4,216).

**Figure S10:**
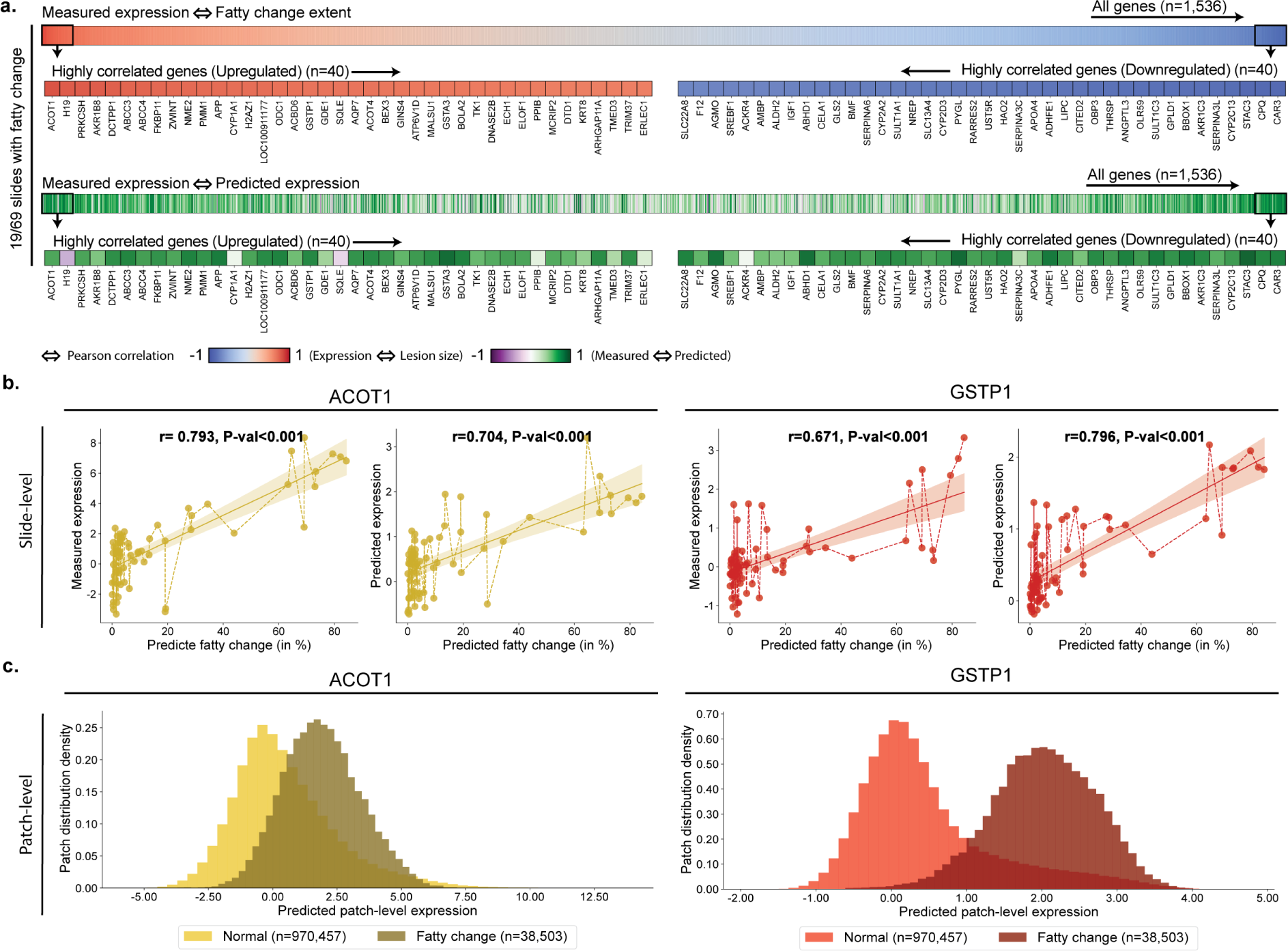
Morphomolecular analysis of fatty change detection in ethionamide. **a.** For each slide-expression pair, we correlate the percentage of fatty change predicted by the lesion classifier with the measured gene expression. Genes with a high correlation between measured expression and lesion size define morphomolecular signatures associated with the compound. **b.** Correlation between the estimated percentage of the slide containing fatty change and the gene expression of *ACOT1* and *GSTP1* (measured with microarray and predicted with GEESE). P-value derived from testing the two-sided null hypothesis of non-correlation. **c.** Distribution of patch-level expression for patches predicted as normal (n=970,457) and patches predicted as containing fatty change (n=38,503).

**Figure S11:**
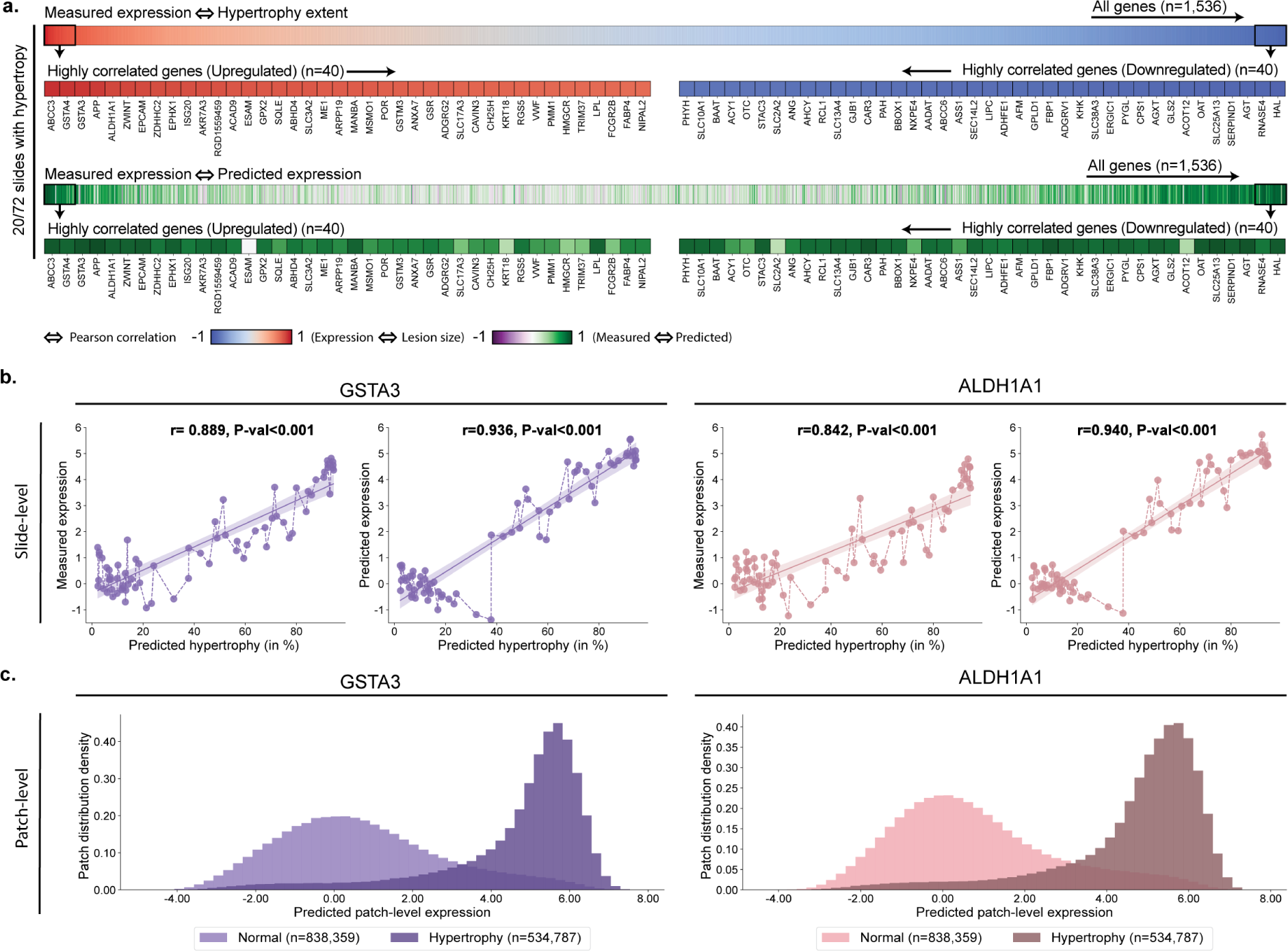
Morphomolecular analysis of hypertrophy detection in hexachlorobenzene. **a.** For each slide-expression pair, we correlate the percentage of hypertrophy predicted by the lesion classifier with the measured gene expression. Genes with a high correlation between measured expression and lesion size define morphomolecular signatures associated with the compound. **b.** Correlation between the estimated percentage of the slide containing hypertrophy and the gene expression of *GSTA3* and *ALDH1A1* (measured with microarray and predicted with GEESE). P-value derived from testing the two-sided null hypothesis of non-correlation. **c.** Distribution of patch-level expression for patches predicted as normal (n=838,359) and patches predicted as containing hypertrophy (n=534,787).

**Figure S12:**
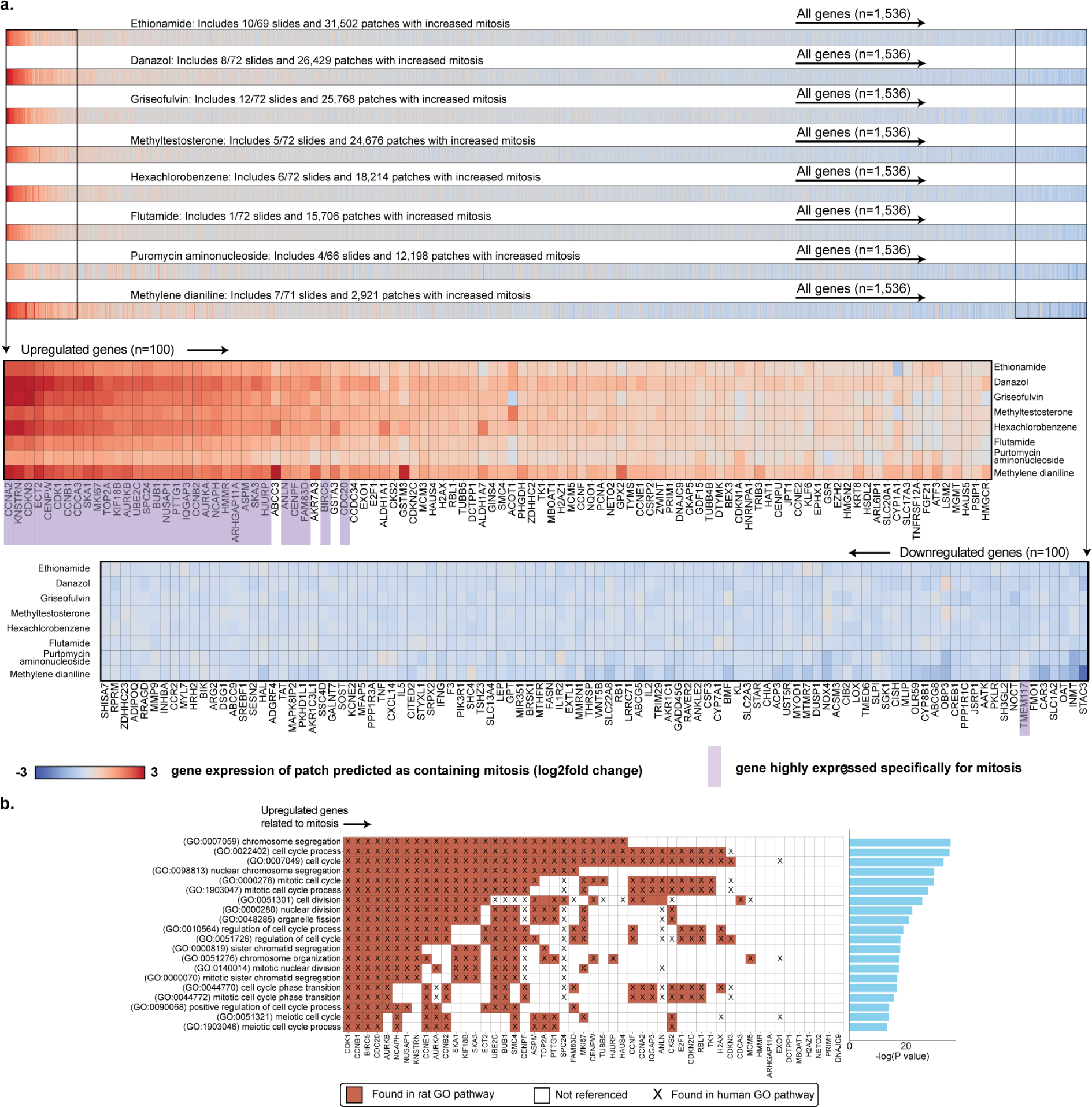
Cross-study analysis of the molecular signature of mitosis. **a.** Average predicted gene expression of patches with mitosis in eight different studies from TG-GATEs test set. Zoom of the top 100 upregulated (top, red) and downregulated genes (bottom, blue). Genes are sorted in decreasing order by their absolute mean gene expression across all considered studies. **b.** Top 20 biological pathways (ranked by highest p-values) of the 51 upregulated genes identified as specific to increased mitosis (**table S9**). The analysis was conducted using the rat genome database (RGD). Identified pathways, including cell cycle, mitotic nuclear division, and DNA replication, are highly relevant to mitosis. Most of the genes found, such as *CCNA2*, *CCNB1*, and *CCNB2* from the cyclin family, are known to play crucial roles in cell division and reproduction pathways are closely related to mitosis. Other genes, such as *HMMR* (involved in microtubule stabilization) and *ARHGAP11A* (associated with Rho GTPase signaling), demonstrate a less direct linkage but are still involved in related processes such as cytoskeletal organization. A lower p-value, obtained using hypergeometric distribution test, signifies a higher degree of over-representation.

**Figure S13:**
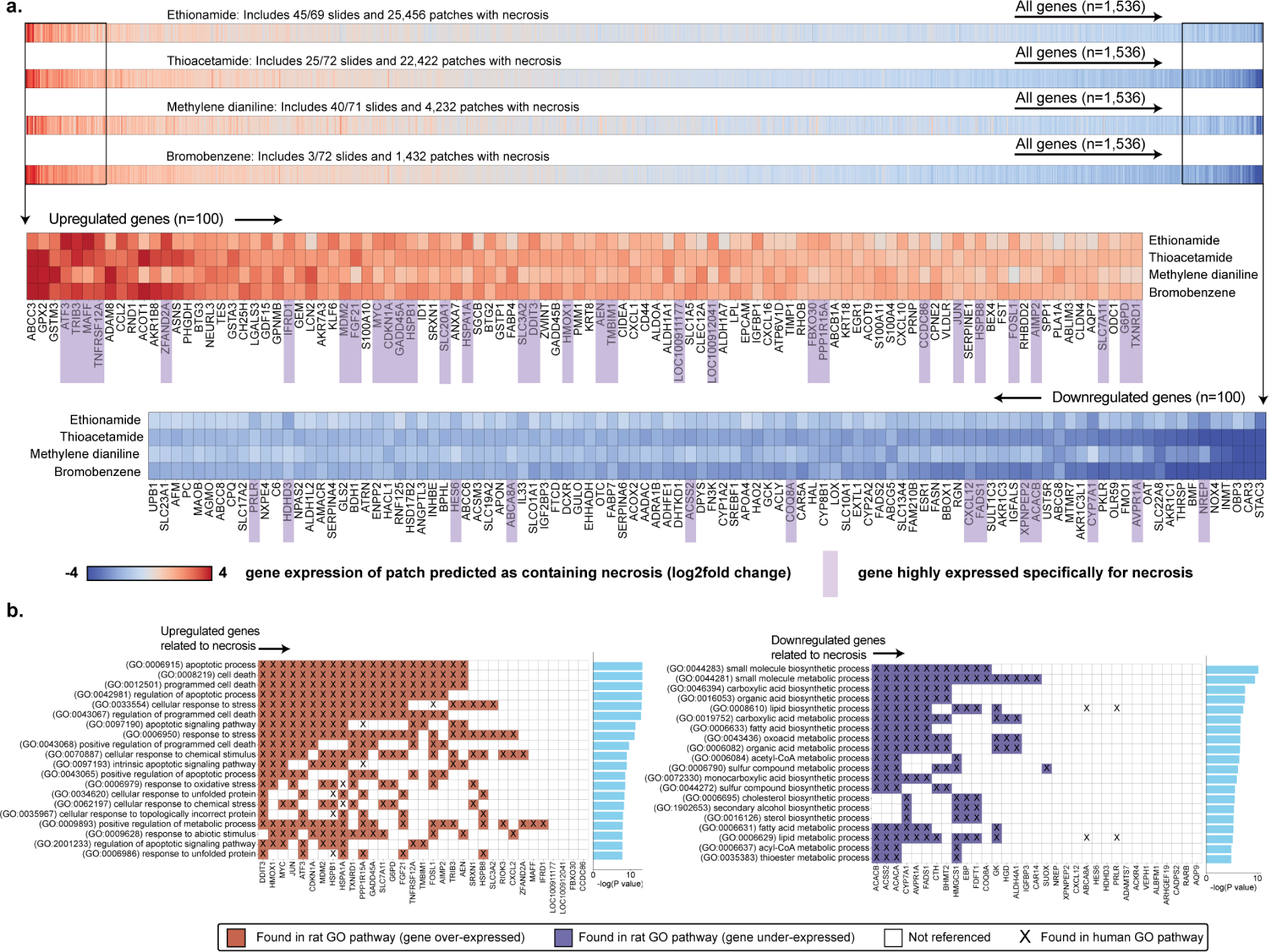
Cross-study analysis of the molecular signature of necrosis. **a.** Average predicted gene expression of patches with necrosis in four different studies from TG-GATEs test set. Zoom of the top 100 upregulated (top, red) and downregulated genes (bottom, blue). Genes are sorted in decreasing order by their absolute mean gene expression across all considered studies. **b.** Top 20 biological pathways (ranked by highest p-values) of the 33 upregulated genes identified as specific to necrosis (left), and the top 20 biological pathways (ranked by highest p-values) of the 33 downregulated genes identified as specific to necrosis (right) (**table S10**). The analysis was conducted using the rat genome database (RGD). Most identified biological pathways, including the apoptotic process, cell death, and cellular response to stress, are highly relevant to cellular necrosis. We observe known genes such as *ATF3* (known for modulating the stress response), *TNFRSF12A* (involved in inflammation and cell death), *HMOX1* (associated with the oxidative stress response), and *MYC* (playing roles in the cell cycle, apoptosis, and DNA damage response). Other genes like *DDIT3*, *TRIB3*, *FOSL1*, and *CXCL2* are also upregulated and associated with processes like response to oxidative stress, cellular response to unfolded protein, and cellular response to chemical stress. Conversely, genes such as *ACSS2*, *ACACA*, *CYP7A1*, and *AVPR1A* are downregulated and linked to small molecule biosynthetic processes, lipid biosynthetic processes, and fatty acid metabolic processes. Other genes like *MAFF* (a gene implicated in transcriptional regulation) or *NREP* (a gene known for its involvement in neural development and tissue re_4_p_7_air) demonstrate a less known direct linkage to necrosis despite showing high correlation with necrosis in our analysis. A lower p-value, obtained using hypergeometric distribution test, signifies a higher degree of over-representation.

**Figure S14:**
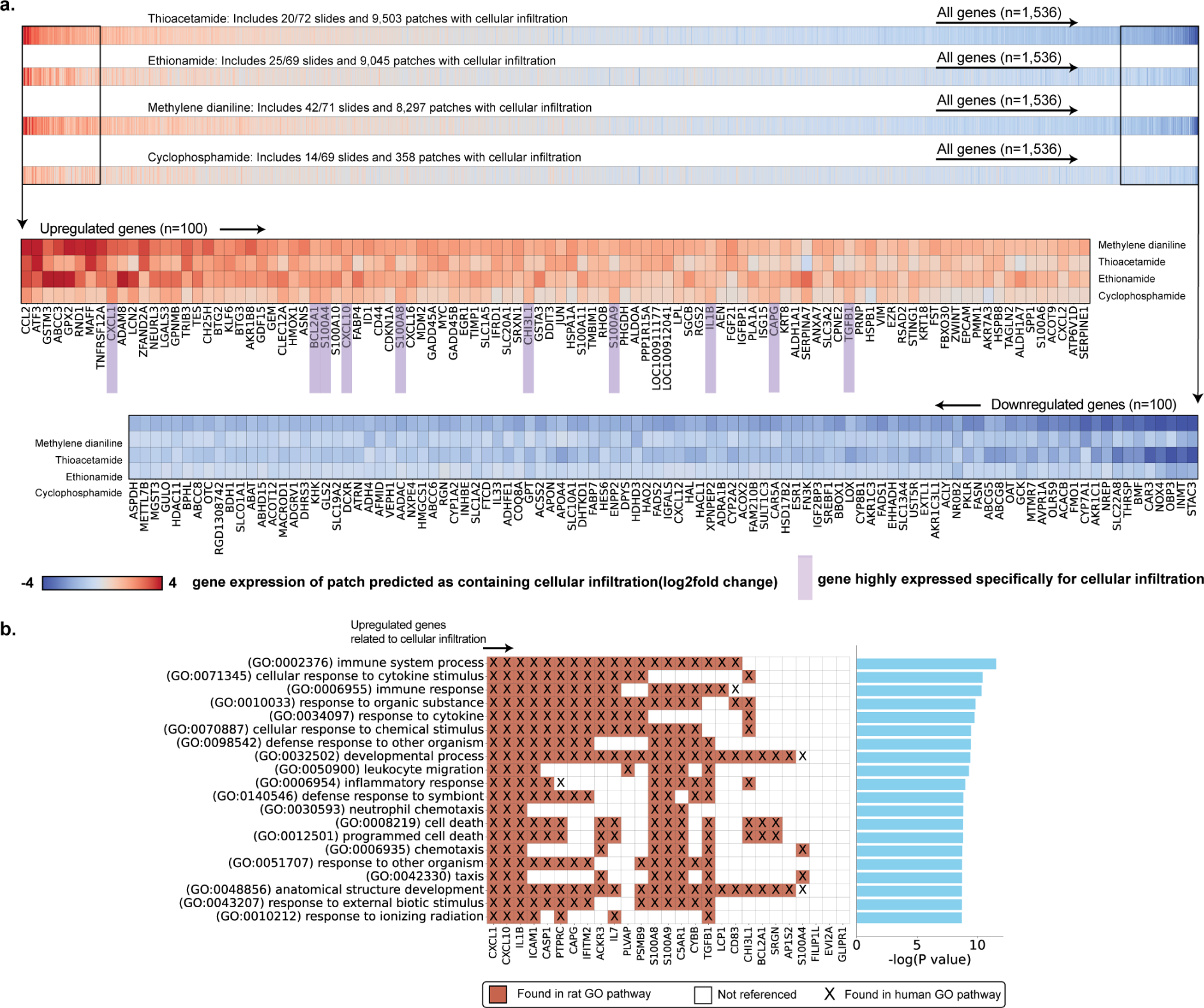
Cross-study analysis of the molecular signature of cellular infiltration. **a.** Average predicted gene expression of patches with cellular infiltration in four different studies from TG-GATEs test set. Zoom of the top 100 upregulated (top, red) and downregulated genes (bottom, blue). Genes are sorted in decreasing order by their absolute mean gene expression across all considered studies. **b.** Top 20 biological pathways (ranked by highest p-values) of the 27 upregulated genes identified as specific to cellular infiltration (**table S11**). Analysis was conducted using the rat genome database (RGD). Genes found such as *CXCL1*, *CXCL10*, noted for their roles in directing leukocytes to the site of infiltration, *BCL2A1*, a gene known for its role in immunity response and the anti-apoptotic role and *S100A4*, involved in modulating the inflammatory response and attracting leukocytes have a biological pathway with links to cellular infiltration. Other genes such as *FILIP1L* and *EVI2A* (gene not currently linked to any specific biological processes in humans and rats according to Gene Ontology Biological Process annotations) demonstrate a less direct linkage to cellular infiltration. A lower p-value, obtained using hypergeometric distribution test, signifies a higher degree of over-representation.

**Figure S15:**
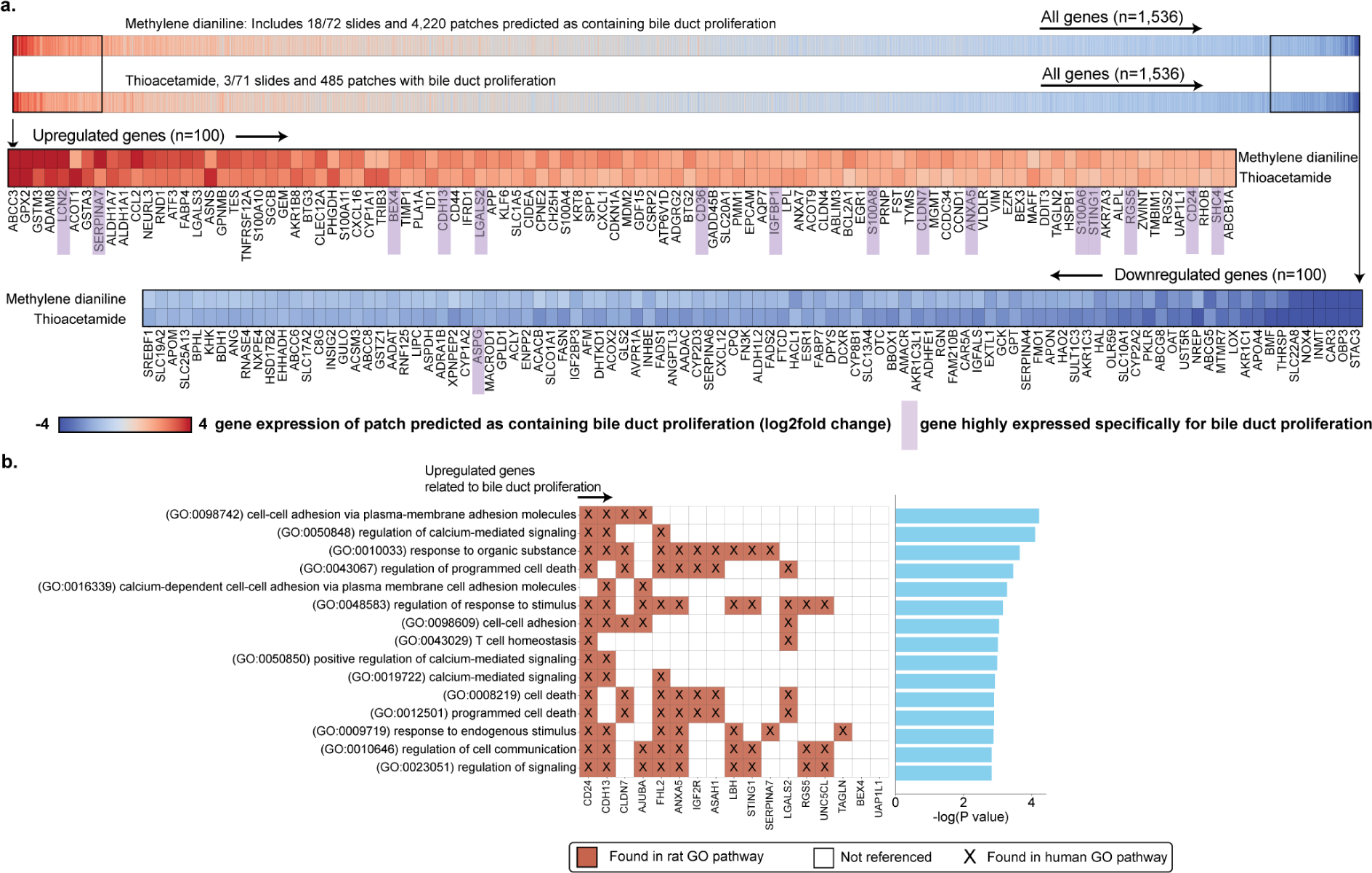
Cross-study analysis of the molecular signature of bile duct proliferation. **a.** Average predicted gene expression of patches with cbile duct proliferation in two studies from TG-GATEs test set. Zoom of the top 100 upregulated (top, red) and downregulated genes (bottom, blue). Genes are sorted in decreasing order by their absolute mean gene expression across all considered studies. **b.** Top 15 biological pathways (highest p-values) of the 17 upregulated genes identified as specific to bile duct proliferation (**table S12**). Analysis was conducted using the rat genome database (RGD). Several genes identified are involved in biological pathways relevant to bile duct proliferation, such as epithelial cell proliferation, extracellular matrix organization, and response to wounding. For instance, genes from the SERPINA family (*SERPINA7* and *SERPINA4*), known for their roles in protease inhibition, may impact inflammation and tissue remodeling processes associated with bile duct proliferation. Other genes like *LGALS2*, involved in cell-cell adhesion, protein-carbohydrate interactions, and immune response modulation, or *CDH13*, known for its role in cell adhesion, also show relevance to the molecular mechanisms underlying bile duct expression. A lower p-value, obtained using hypergeometric distribution test, signifies a higher degree of over-representation.

**Figure S16:**
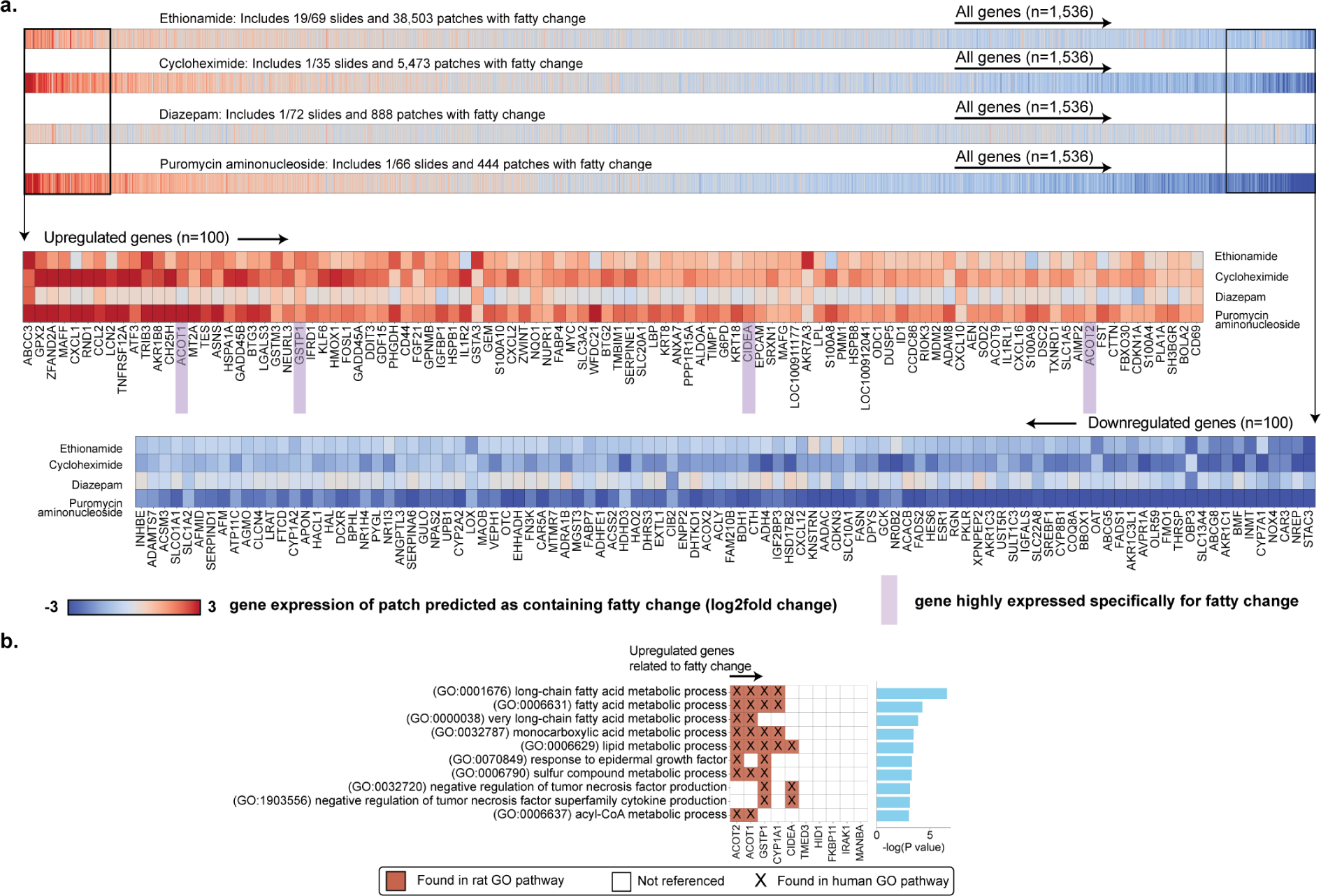
Cross-study analysis of the molecular signature of fatty change. **a.** Average predicted gene expression of patches with fatty change in four different studies from TG-GATEs test set. Zoom of the top 100 upregulated (top, red) and downregulated genes (bottom, blue). Genes are sorted in decreasing order by their absolute mean gene expression across all considered studies. **b.** Top 10 biological pathways (ranked by highest p-values) linked to the 10 upregulated genes identified as specific to fatty change (**table S13**). Analysis was conducted using the rat genome database (RGD). Some identified pathways, including fatty acid metabolic process, lipid metabolic process, and response to oxidative stress, support the relevance of the identified genes to fatty change. Genes such as *ACOT1* and *ACOT2* from the ACOT family, which are involved in the hydrolysis of acyl-CoA thioester compounds, a key process in lipid metabolism are highly relevant to fatty change. Additionally, *GSTP1*, a gene implicated in detoxification and the response to oxidative stress, is also upregulated in fatty change. A lower p-value, obtained using hypergeometric distribution test, signifies a higher degree of over-representation.

**Figure S17:**
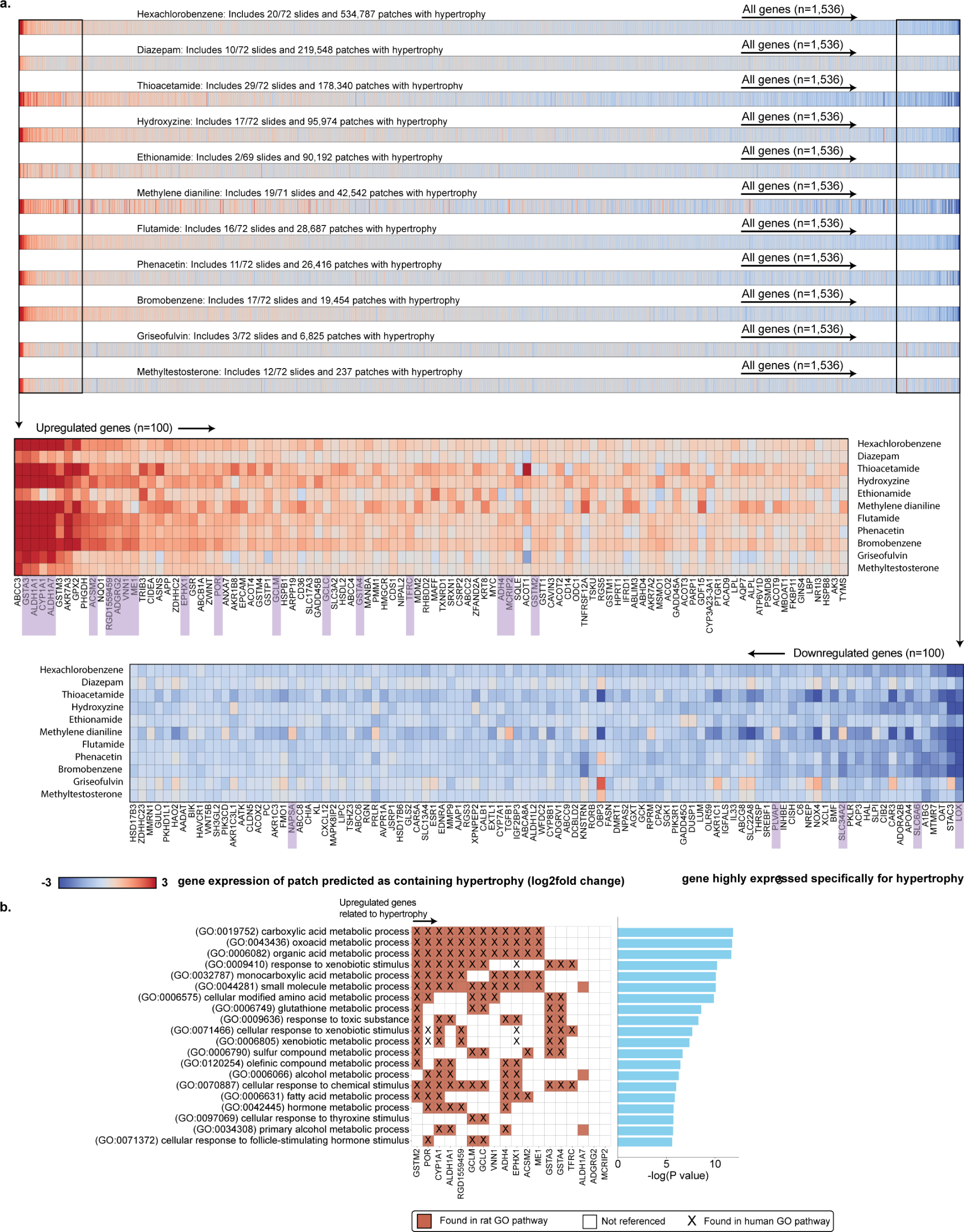
Cross-study analysis of the molecular signature of hypertrophy. **a.** Average predicted gene expression of patches with hypertrophy in eleven different studies from TG-GATEs test set. Zoom of the top 100 upregulated (top, red) and downregulated genes (bottom, blue). Genes are sorted in decreasing order by their absolute mean gene expression across all considered studies. **b.** Top 20 biological pathways (ranked by highest p-values) linked to the 18 genes upregulated genes identified as specific to hypertrophy (**table S14**). Analysis was conducted using the rat genome database (RGD). Genes involved in managing increased metabolic demands, such as *ACSM2*, *VNN1*, and *ME1*, are relevant in different aspects of metabolism—fatty acid, lipid, and glucose metabolism, respectively. Genes like *GSTA3*, *ALDH1A1*, and *ALDH1A7* are involved in detoxifying oxidative and aldehyde by-products, which are more prevalent as metabolic activities intensify in hypertrophic cells. The enrichment of pathways such as fatty acid metabolic process, oxidation-reduction process, and extracellular matrix organization further supports the relevance of these genes to the cellular adaptations occurring during hypertrophy. Additionally, *LOX*’s role in collagen synthesis and extracellular matrix assembly is necessary for the structural adaptation of tissues undergoing hypertrophy. A lower p-value, obtained using hypergeometric distribution test, signifies a higher degree of over-representation.

**Figure S18:**
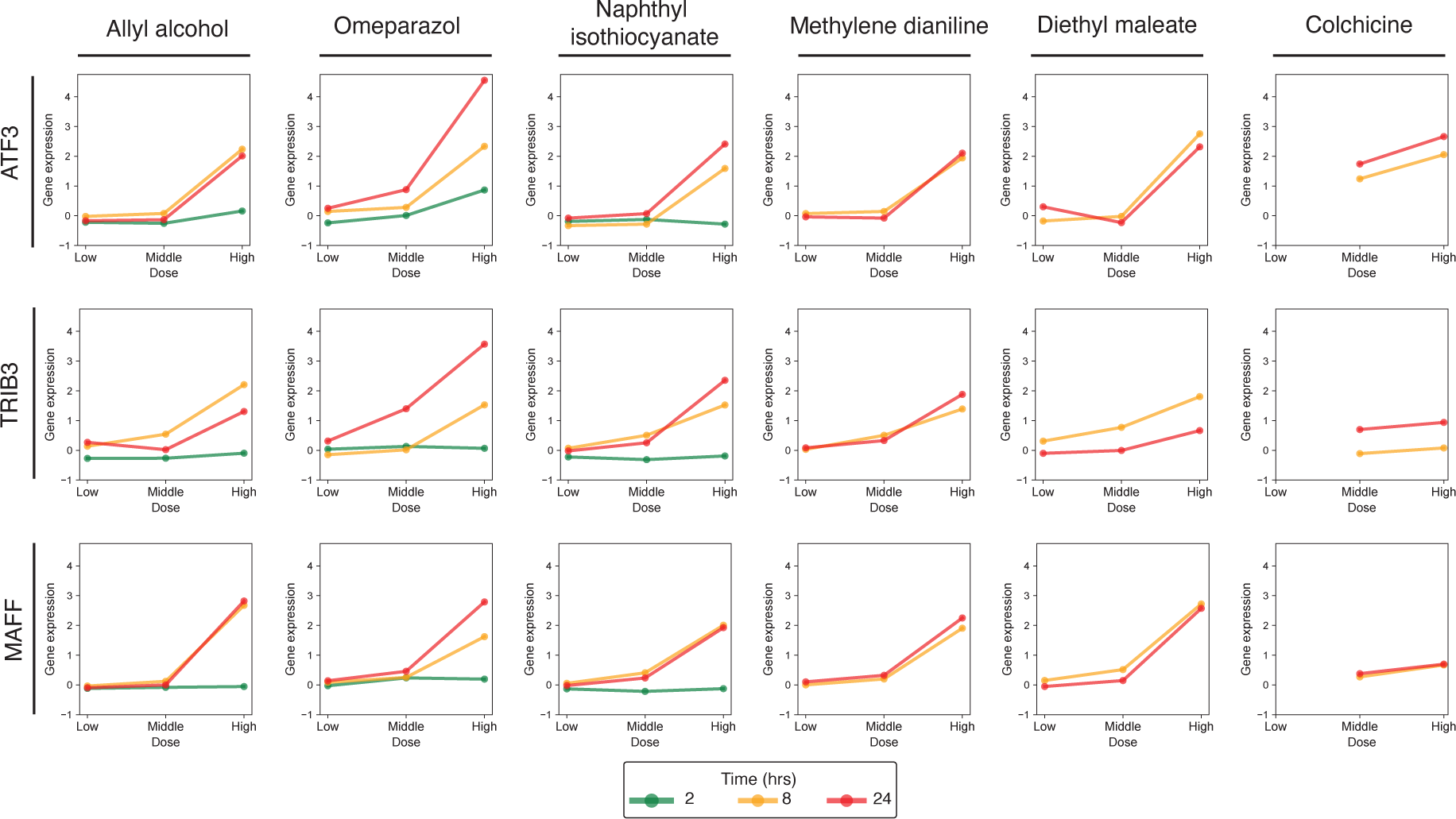
Translation of necrosis genetic biomarkers to human in vitro primary human hepatocytes cell lines. Dose-response relationship of six necrosis-inducing selected compounds of in vitro human cell lines at three different time points. Genes tested (*ATF3*, *TRIB3*, and *MAFF*) were identified as connected to necrosis in the in vivo rat experiments.

**Table S1:**
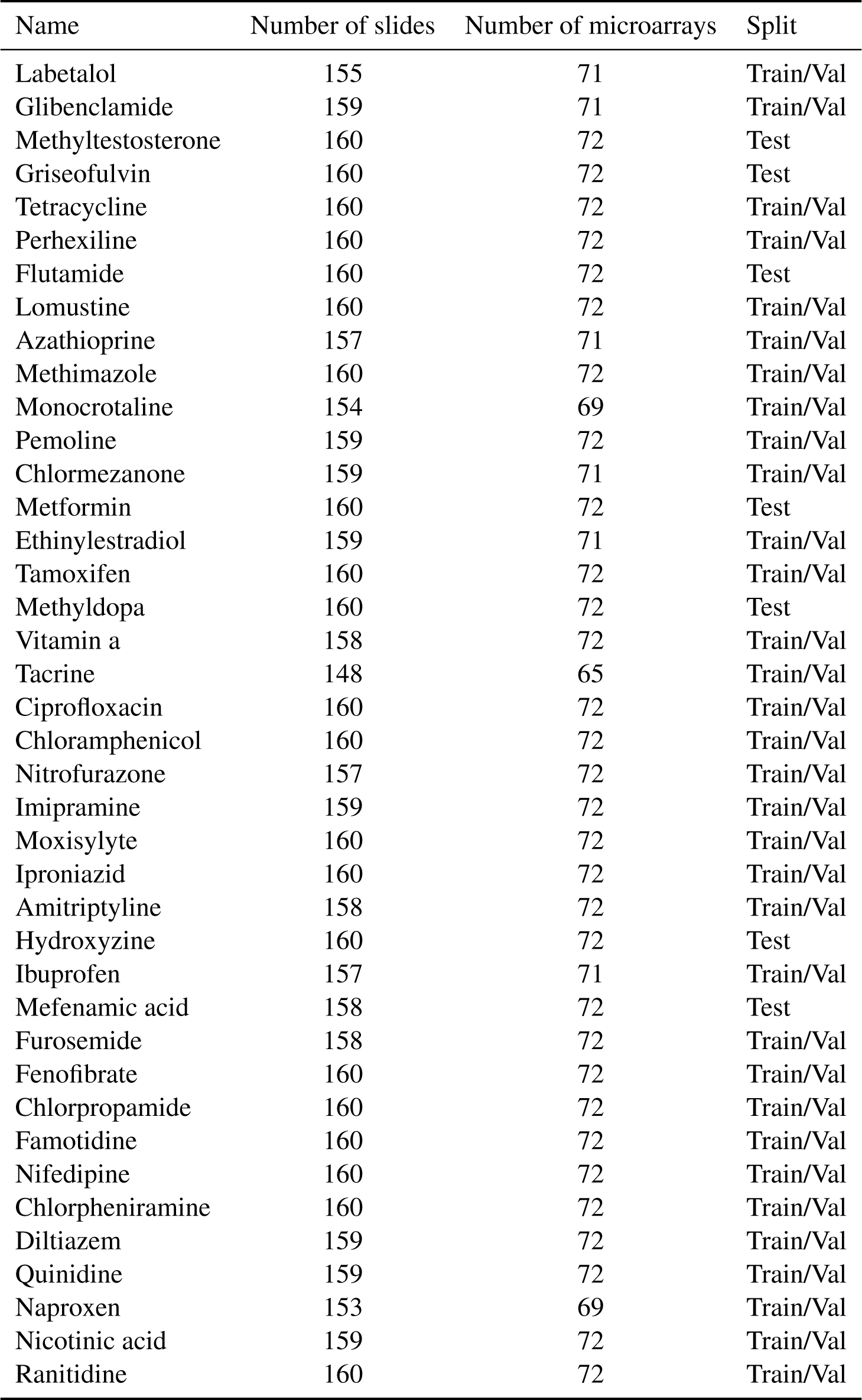
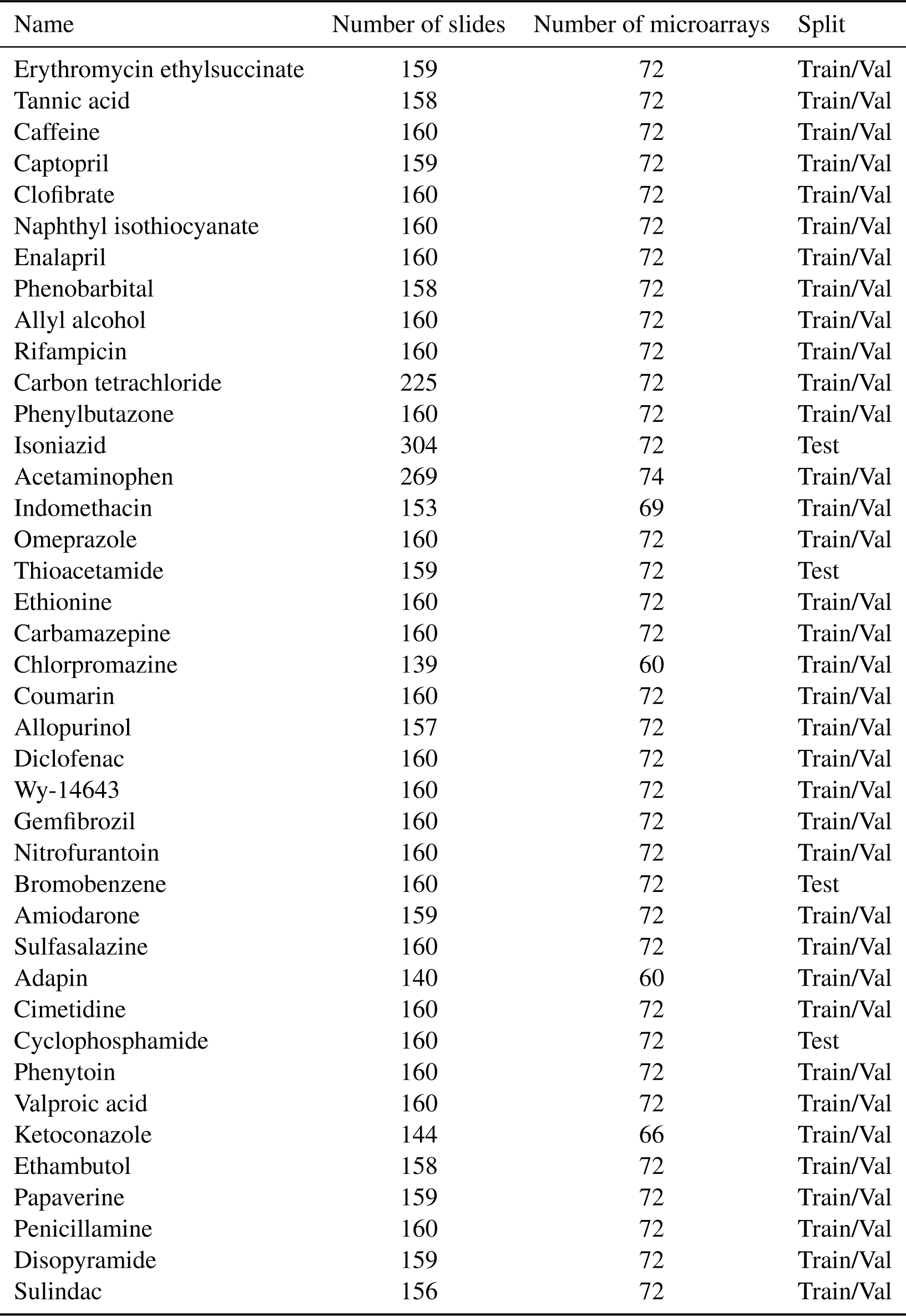
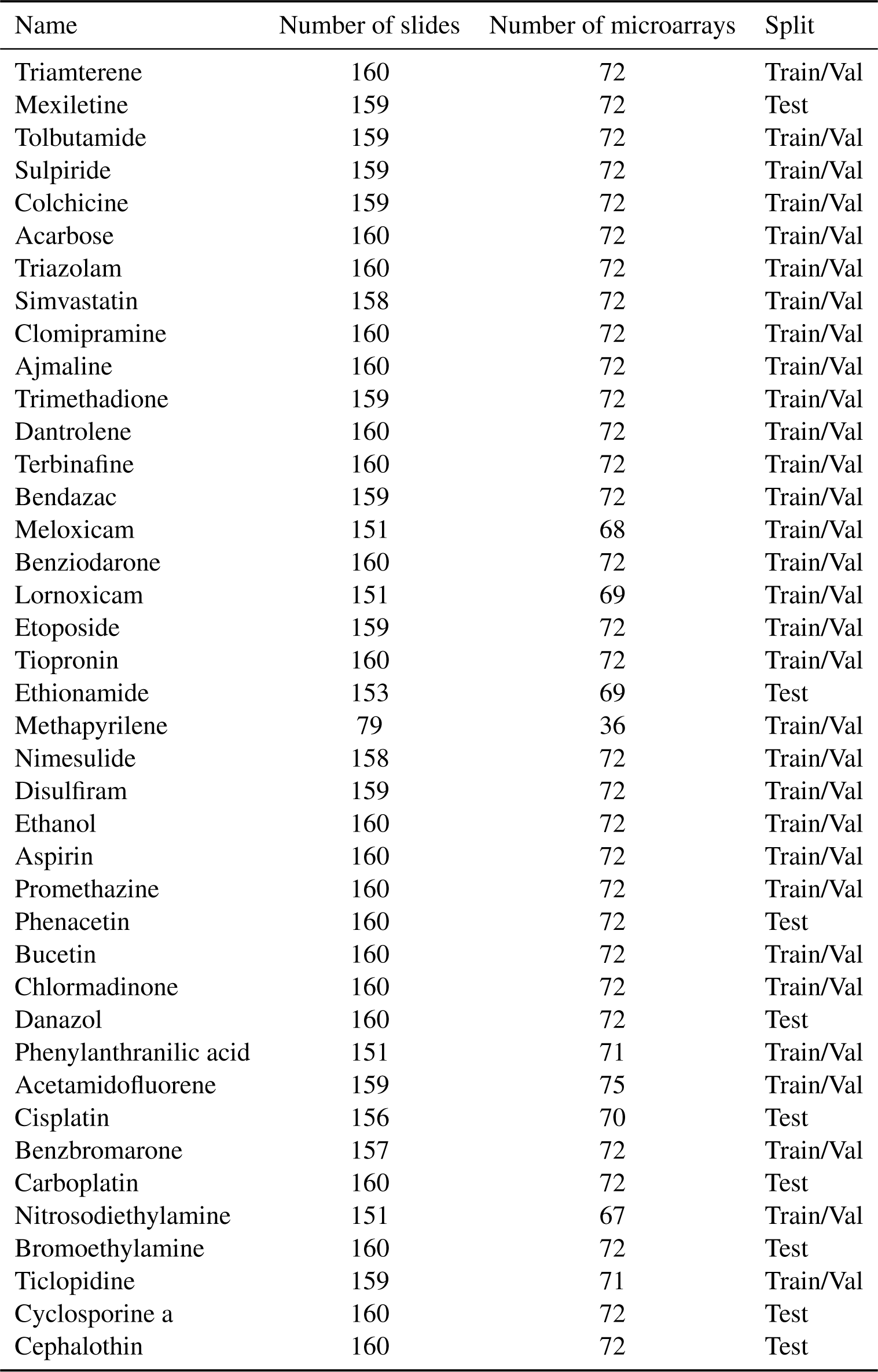
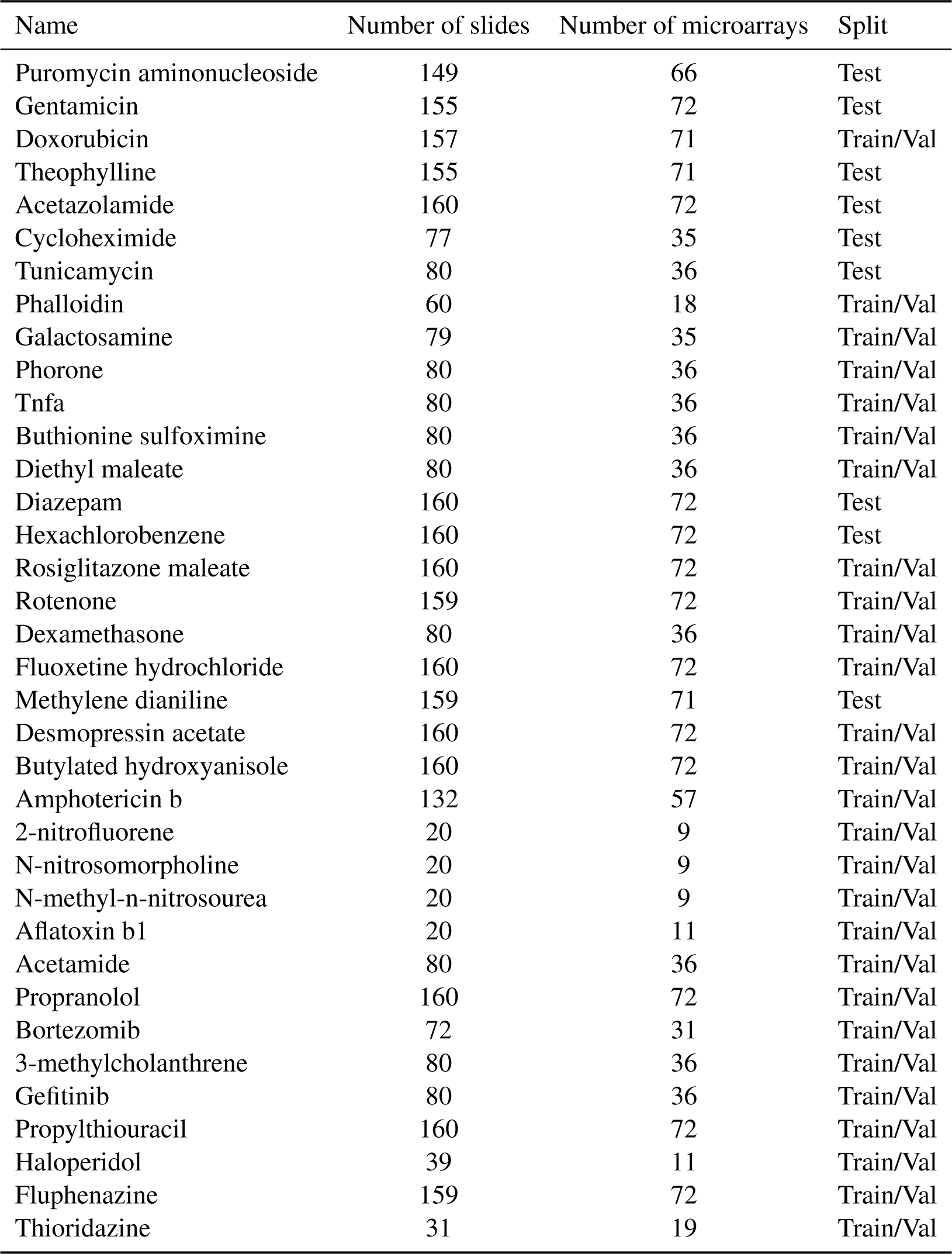
Overview of TG-GATEs compounds. For each compound, we report the number of slides, microarrays (also corresponding to the number of pairs), and whether the study was used for training/validation or testing.

| Name | Number of slides | Number of microarrays | Split |
| --- | --- | --- | --- |
| Labetalol | 155 | 71 | Train/Val |
| Glibenclamide | 159 | 71 | Train/Val |
| Methyltestosterone | 160 | 72 | Test |
| Griseofulvin | 160 | 72 | Test |
| Tetracycline | 160 | 72 | Train/Val |
| Perhexiline | 160 | 72 | Train/Val |
| Flutamide | 160 | 72 | Test |
| Lomustine | 160 | 72 | Train/Val |
| Azathioprine | 157 | 71 | Train/Val |
| Methimazole | 160 | 72 | Train/Val |
| Monocrotaline | 154 | 69 | Train/Val |
| Pemoline | 159 | 72 | Train/Val |
| Chlormezanone | 159 | 71 | Train/Val |
| Metformin | 160 | 72 | Test |
| Ethinylestradiol | 159 | 71 | Train/Val |
| Tamoxifen | 160 | 72 | Train/Val |
| Methyldopa | 160 | 72 | Test |
| Vitamin a | 158 | 72 | Train/Val |
| Tacrine | 148 | 65 | Train/Val |
| Ciprofloxacin | 160 | 72 | Train/Val |
| Chloramphenicol | 160 | 72 | Train/Val |
| Nitrofurazone | 157 | 72 | Train/Val |
| Imipramine | 159 | 72 | Train/Val |
| Moxisylyte | 160 | 72 | Train/Val |
| Iproniazid | 160 | 72 | Train/Val |
| Amitriptyline | 158 | 72 | Train/Val |
| Hydroxyzine | 160 | 72 | Test |
| Ibuprofen | 157 | 71 | Train/Val |
| Mefenamic acid | 158 | 72 | Test |
| Furosemide | 158 | 72 | Train/Val |
| Fenofibrate | 160 | 72 | Train/Val |
| Chlorpropamide | 160 | 72 | Train/Val |
| Famotidine | 160 | 72 | Train/Val |
| Nifedipine | 160 | 72 | Train/Val |
| Chlorpheniramine | 160 | 72 | Train/Val |
| Diltiazem | 159 | 72 | Train/Val |
| Quinidine | 159 | 72 | Train/Val |
| Naproxen | 153 | 69 | Train/Val |
| Nicotinic acid | 159 | 72 | Train/Val |
| Ranitidine | 160 | 72 | Train/Val |
| Erythromycin ethylsuccinate | 159 | 72 | Train/Val |
| Tannic acid | 158 | 72 | Train/Val |
| Caffeine | 160 | 72 | Train/Val |
| Captopril | 159 | 72 | Train/Val |
| Clofibrate | 160 | 72 | Train/Val |
| Naphthyl isothiocyanate | 160 | 72 | Train/Val |
| Enalapril | 160 | 72 | Train/Val |
| Phenobarbital | 158 | 72 | Train/Val |
| Allyl alcohol | 160 | 72 | Train/Val |
| Rifampicin | 160 | 72 | Train/Val |
| Carbon tetrachloride | 225 | 72 | Train/Val |
| Phenylbutazone | 160 | 72 | Train/Val |
| Isoniazid | 304 | 72 | Test |
| Acetaminophen | 269 | 74 | Train/Val |
| Indomethacin | 153 | 69 | Train/Val |
| Omeprazole | 160 | 72 | Train/Val |
| Thioacetamide | 159 | 72 | Test |
| Ethionine | 160 | 72 | Train/Val |
| Carbamazepine | 160 | 72 | Train/Val |
| Chlorpromazine | 139 | 60 | Train/Val |
| Coumarin | 160 | 72 | Train/Val |
| Allopurinol | 157 | 72 | Train/Val |
| Diclofenac | 160 | 72 | Train/Val |
| Wy-14643 | 160 | 72 | Train/Val |
| Gemfibrozil | 160 | 72 | Train/Val |
| Nitrofurantoin | 160 | 72 | Train/Val |
| Bromobenzene | 160 | 72 | Test |
| Amiodarone | 159 | 72 | Train/Val |
| Sulfasalazine | 160 | 72 | Train/Val |
| Adapin | 140 | 60 | Train/Val |
| Cimetidine | 160 | 72 | Train/Val |
| Cyclophosphamide | 160 | 72 | Test |
| Phenytoin | 160 | 72 | Train/Val |
| Valproic acid | 160 | 72 | Train/Val |
| Ketoconazole | 144 | 66 | Train/Val |
| Ethambutol | 158 | 72 | Train/Val |
| Papaverine | 159 | 72 | Train/Val |
| Penicillamine | 160 | 72 | Train/Val |
| Disopyramide | 159 | 72 | Train/Val |
| Sulindac | 156 | 72 | Train/Val |
| Triamterene | 160 | 72 | Train/Val |
| Mexiletine | 159 | 72 | Test |
| Tolbutamide | 159 | 72 | Train/Val |
| Sulpiride | 159 | 72 | Train/Val |
| Colchicine | 159 | 72 | Train/Val |
| Acarbose | 160 | 72 | Train/Val |
| Triazolam | 160 | 72 | Train/Val |
| Simvastatin | 158 | 72 | Train/Val |
| Clomipramine | 160 | 72 | Train/Val |
| Ajmaline | 160 | 72 | Train/Val |
| Trimethadione | 159 | 72 | Train/Val |
| Dantrolene | 160 | 72 | Train/Val |
| Terbinafine | 160 | 72 | Train/Val |
| Bendazac | 159 | 72 | Train/Val |
| Meloxicam | 151 | 68 | Train/Val |
| Benziodarone | 160 | 72 | Train/Val |
| Lornoxicam | 151 | 69 | Train/Val |
| Etoposide | 159 | 72 | Train/Val |
| Tiopronin | 160 | 72 | Train/Val |
| Ethionamide | 153 | 69 | Test |
| Methapyrilene | 79 | 36 | Train/Val |
| Nimesulide | 158 | 72 | Train/Val |
| Disulfiram | 159 | 72 | Train/Val |
| Ethanol | 160 | 72 | Train/Val |
| Aspirin | 160 | 72 | Train/Val |
| Promethazine | 160 | 72 | Train/Val |
| Phenacetin | 160 | 72 | Test |
| Bucetin | 160 | 72 | Train/Val |
| Chlormadinone | 160 | 72 | Train/Val |
| Danazol | 160 | 72 | Test |
| Phenylanthranilic acid | 151 | 71 | Train/Val |
| Acetamidofluorene | 159 | 75 | Train/Val |
| Cisplatin | 156 | 70 | Test |
| Benzbromarone | 157 | 72 | Train/Val |
| Carboplatin | 160 | 72 | Test |
| Nitrosodiethylamine | 151 | 67 | Train/Val |
| Bromoethylamine | 160 | 72 | Test |
| Ticlopidine | 159 | 71 | Train/Val |
| Cyclosporine a | 160 | 72 | Test |
| Cephalothin | 160 | 72 | Test |
| Puromycin aminonucleoside | 149 | 66 | Test |
| Gentamicin | 155 | 72 | Test |
| Doxorubicin | 157 | 71 | Train/Val |
| Theophylline | 155 | 71 | Test |
| Acetazolamide | 160 | 72 | Test |
| Cycloheximide | 77 | 35 | Test |
| Tunicamycin | 80 | 36 | Test |
| Phalloidin | 60 | 18 | Train/Val |
| Galactosamine | 79 | 35 | Train/Val |
| Phorone | 80 | 36 | Train/Val |
| Tnfa | 80 | 36 | Train/Val |
| Buthionine sulfoximine | 80 | 36 | Train/Val |
| Diethyl maleate | 80 | 36 | Train/Val |
| Diazepam | 160 | 72 | Test |
| Hexachlorobenzene | 160 | 72 | Test |
| Rosiglitazone maleate | 160 | 72 | Train/Val |
| Rotenone | 159 | 72 | Train/Val |
| Dexamethasone | 80 | 36 | Train/Val |
| Fluoxetine hydrochloride | 160 | 72 | Train/Val |
| Methylene dianiline | 159 | 71 | Test |
| Desmopressin acetate | 160 | 72 | Train/Val |
| Butylated hydroxyanisole | 160 | 72 | Train/Val |
| Amphotericin b | 132 | 57 | Train/Val |
| 2-nitrofluorene | 20 | 9 | Train/Val |
| N-nitrosomorpholine | 20 | 9 | Train/Val |
| N-methyl-n-nitrosourea | 20 | 9 | Train/Val |
| Aflatoxin b1 | 20 | 11 | Train/Val |
| Acetamide | 80 | 36 | Train/Val |
| Propranolol | 160 | 72 | Train/Val |
| Bortezomib | 72 | 31 | Train/Val |
| 3-methylcholanthrene | 80 | 36 | Train/Val |
| Gefitinib | 80 | 36 | Train/Val |
| Propylthiouracil | 160 | 72 | Train/Val |
| Haloperidol | 39 | 11 | Train/Val |
| Fluphenazine | 159 | 72 | Train/Val |
| Thioridazine | 31 | 19 | Train/Val |

**Table S2:**
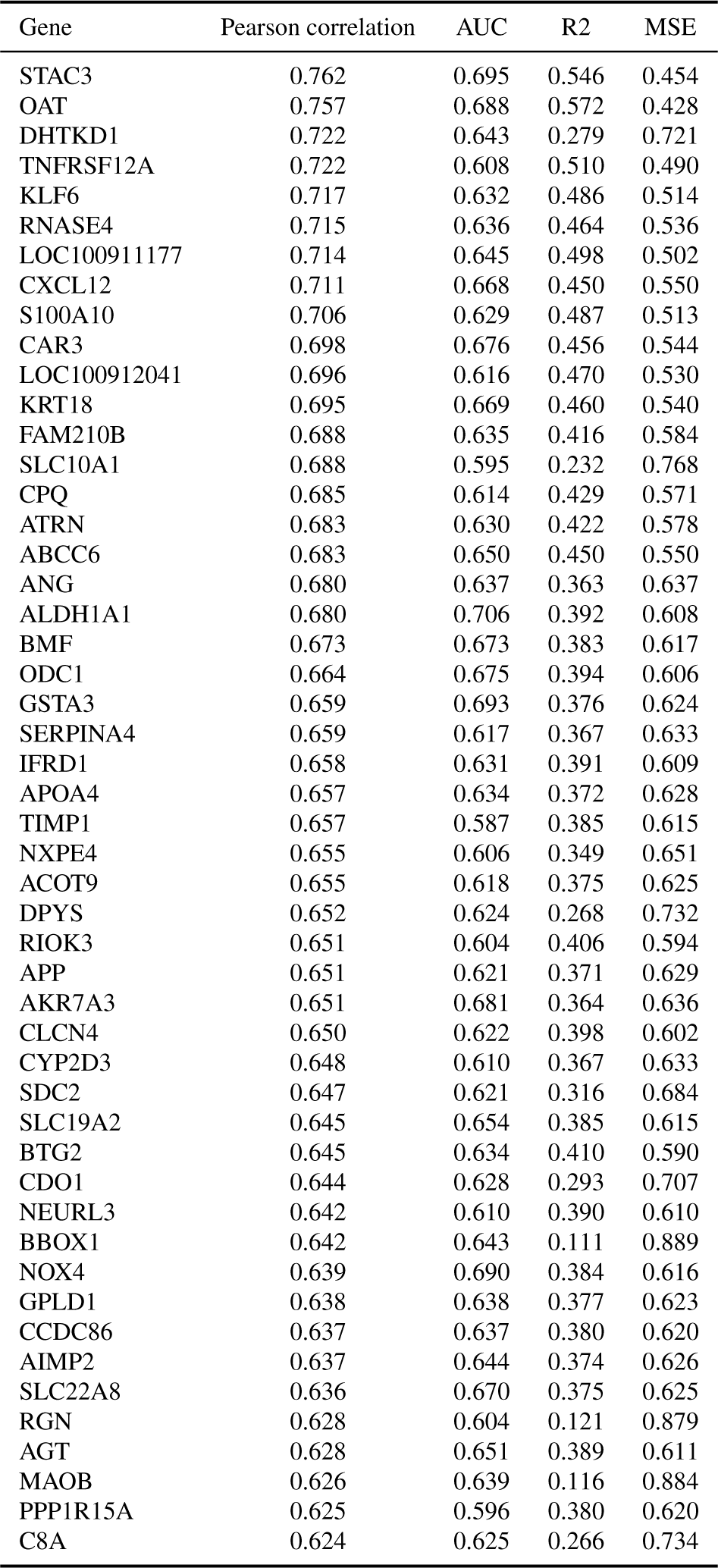
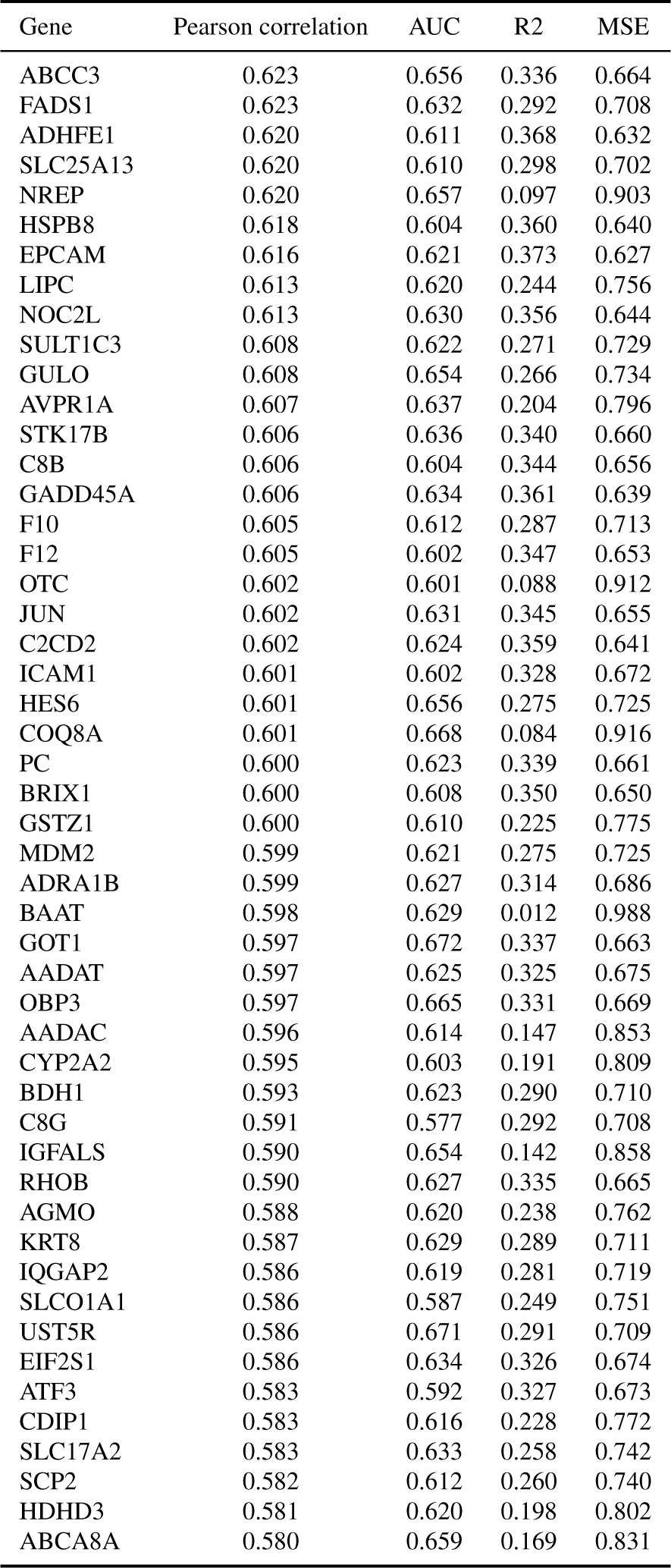
Gene expression prediction performance of top-100 best predicted genes. We report Pearson correlation, AUC, R2, and MSE. Each metric is described in the **Online methods**, section **Metrics**.

| Gene | Pearson correlation | AUC | R2 | MSE |
| --- | --- | --- | --- | --- |
| STAC3 | 0.762 | 0.695 | 0.546 | 0.454 |
| OAT | 0.757 | 0.688 | 0.572 | 0.428 |
| DHTKD1 | 0.722 | 0.643 | 0.279 | 0.721 |
| TNFRSF12A | 0.722 | 0.608 | 0.510 | 0.490 |
| KLF6 | 0.717 | 0.632 | 0.486 | 0.514 |
| RNASE4 | 0.715 | 0.636 | 0.464 | 0.536 |
| LOC100911177 | 0.714 | 0.645 | 0.498 | 0.502 |
| CXCL12 | 0.711 | 0.668 | 0.450 | 0.550 |
| S100A10 | 0.706 | 0.629 | 0.487 | 0.513 |
| CAR3 | 0.698 | 0.676 | 0.456 | 0.544 |
| LOC100912041 | 0.696 | 0.616 | 0.470 | 0.530 |
| KRT18 | 0.695 | 0.669 | 0.460 | 0.540 |
| FAM210B | 0.688 | 0.635 | 0.416 | 0.584 |
| SLC10A1 | 0.688 | 0.595 | 0.232 | 0.768 |
| CPQ | 0.685 | 0.614 | 0.429 | 0.571 |
| ATRN | 0.683 | 0.630 | 0.422 | 0.578 |
| ABCC6 | 0.683 | 0.650 | 0.450 | 0.550 |
| ANG | 0.680 | 0.637 | 0.363 | 0.637 |
| ALDH1A1 | 0.680 | 0.706 | 0.392 | 0.608 |
| BMF | 0.673 | 0.673 | 0.383 | 0.617 |
| ODC1 | 0.664 | 0.675 | 0.394 | 0.606 |
| GSTA3 | 0.659 | 0.693 | 0.376 | 0.624 |
| SERPINA4 | 0.659 | 0.617 | 0.367 | 0.633 |
| IFRD1 | 0.658 | 0.631 | 0.391 | 0.609 |
| APOA4 | 0.657 | 0.634 | 0.372 | 0.628 |
| TIMP1 | 0.657 | 0.587 | 0.385 | 0.615 |
| NXPE4 | 0.655 | 0.606 | 0.349 | 0.651 |
| ACOT9 | 0.655 | 0.618 | 0.375 | 0.625 |
| DPYS | 0.652 | 0.624 | 0.268 | 0.732 |
| RIOK3 | 0.651 | 0.604 | 0.406 | 0.594 |
| APP | 0.651 | 0.621 | 0.371 | 0.629 |
| AKR7A3 | 0.651 | 0.681 | 0.364 | 0.636 |
| CLCN4 | 0.650 | 0.622 | 0.398 | 0.602 |
| CYP2D3 | 0.648 | 0.610 | 0.367 | 0.633 |
| SDC2 | 0.647 | 0.621 | 0.316 | 0.684 |
| SLC19A2 | 0.645 | 0.654 | 0.385 | 0.615 |
| BTG2 | 0.645 | 0.634 | 0.410 | 0.590 |
| CDO1 | 0.644 | 0.628 | 0.293 | 0.707 |
| NEURL3 | 0.642 | 0.610 | 0.390 | 0.610 |
| BBOX1 | 0.642 | 0.643 | 0.111 | 0.889 |
| NOX4 | 0.639 | 0.690 | 0.384 | 0.616 |
| GPLD1 | 0.638 | 0.638 | 0.377 | 0.623 |
| CCDC86 | 0.637 | 0.637 | 0.380 | 0.620 |
| AIMP2 | 0.637 | 0.644 | 0.374 | 0.626 |
| SLC22A8 | 0.636 | 0.670 | 0.375 | 0.625 |
| RGN | 0.628 | 0.604 | 0.121 | 0.879 |
| AGT | 0.628 | 0.651 | 0.389 | 0.611 |
| MAOB | 0.626 | 0.639 | 0.116 | 0.884 |
| PPP1R15A | 0.625 | 0.596 | 0.380 | 0.620 |
| C8A | 0.624 | 0.625 | 0.266 | 0.734 |
| ABCC3 | 0.623 | 0.656 | 0.336 | 0.664 |
| FADS1 | 0.623 | 0.632 | 0.292 | 0.708 |
| ADHFE1 | 0.620 | 0.611 | 0.368 | 0.632 |
| SLC25A13 | 0.620 | 0.610 | 0.298 | 0.702 |
| NREP | 0.620 | 0.657 | 0.097 | 0.903 |
| HSPB8 | 0.618 | 0.604 | 0.360 | 0.640 |
| EPCAM | 0.616 | 0.621 | 0.373 | 0.627 |
| LIPC | 0.613 | 0.620 | 0.244 | 0.756 |
| NOC2L | 0.613 | 0.630 | 0.356 | 0.644 |
| SULT1C3 | 0.608 | 0.622 | 0.271 | 0.729 |
| GULO | 0.608 | 0.654 | 0.266 | 0.734 |
| AVPR1A | 0.607 | 0.637 | 0.204 | 0.796 |
| STK17B | 0.606 | 0.636 | 0.340 | 0.660 |
| C8B | 0.606 | 0.604 | 0.344 | 0.656 |
| GADD45A | 0.606 | 0.634 | 0.361 | 0.639 |
| F10 | 0.605 | 0.612 | 0.287 | 0.713 |
| F12 | 0.605 | 0.602 | 0.347 | 0.653 |
| OTC | 0.602 | 0.601 | 0.088 | 0.912 |
| JUN | 0.602 | 0.631 | 0.345 | 0.655 |
| C2CD2 | 0.602 | 0.624 | 0.359 | 0.641 |
| ICAM1 | 0.601 | 0.602 | 0.328 | 0.672 |
| HES6 | 0.601 | 0.656 | 0.275 | 0.725 |
| COQ8A | 0.601 | 0.668 | 0.084 | 0.916 |
| PC | 0.600 | 0.623 | 0.339 | 0.661 |
| BRIX1 | 0.600 | 0.608 | 0.350 | 0.650 |
| GSTZ1 | 0.600 | 0.610 | 0.225 | 0.775 |
| MDM2 | 0.599 | 0.621 | 0.275 | 0.725 |
| ADRA1B | 0.599 | 0.627 | 0.314 | 0.686 |
| BAAT | 0.598 | 0.629 | 0.012 | 0.988 |
| GOT1 | 0.597 | 0.672 | 0.337 | 0.663 |
| AADAT | 0.597 | 0.625 | 0.325 | 0.675 |
| OBP3 | 0.597 | 0.665 | 0.331 | 0.669 |
| AADAC | 0.596 | 0.614 | 0.147 | 0.853 |
| CYP2A2 | 0.595 | 0.603 | 0.191 | 0.809 |
| BDH1 | 0.593 | 0.623 | 0.290 | 0.710 |
| C8G | 0.591 | 0.577 | 0.292 | 0.708 |
| IGFALS | 0.590 | 0.654 | 0.142 | 0.858 |
| RHOB | 0.590 | 0.627 | 0.335 | 0.665 |
| AGMO | 0.588 | 0.620 | 0.238 | 0.762 |
| KRT8 | 0.587 | 0.629 | 0.289 | 0.711 |
| IQGAP2 | 0.586 | 0.619 | 0.281 | 0.719 |
| SLCO1A1 | 0.586 | 0.587 | 0.249 | 0.751 |
| UST5R | 0.586 | 0.671 | 0.291 | 0.709 |
| EIF2S1 | 0.586 | 0.634 | 0.326 | 0.674 |
| ATF3 | 0.583 | 0.592 | 0.327 | 0.673 |
| CDIP1 | 0.583 | 0.616 | 0.228 | 0.772 |
| SLC17A2 | 0.583 | 0.633 | 0.258 | 0.742 |
| SCP2 | 0.582 | 0.612 | 0.260 | 0.740 |
| HDHD3 | 0.581 | 0.620 | 0.198 | 0.802 |
| ABCA8A | 0.580 | 0.659 | 0.169 | 0.831 |

**Table S3:**
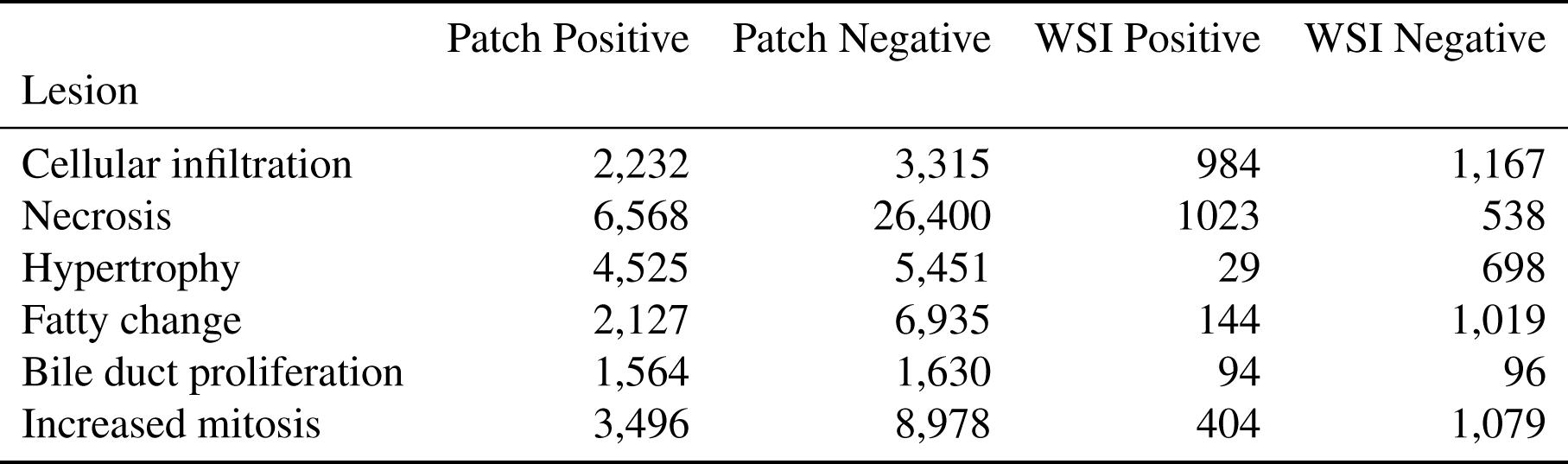
Number of patch-level annotations per lesion. Patch positive refers to the number of patches containing the lesion, and Patch negative refers to the number of patches that do not contain it (can be a normal patch or a patch containing another lesion). A description of each lesion is provided in **table S4**.

**Table S4:**
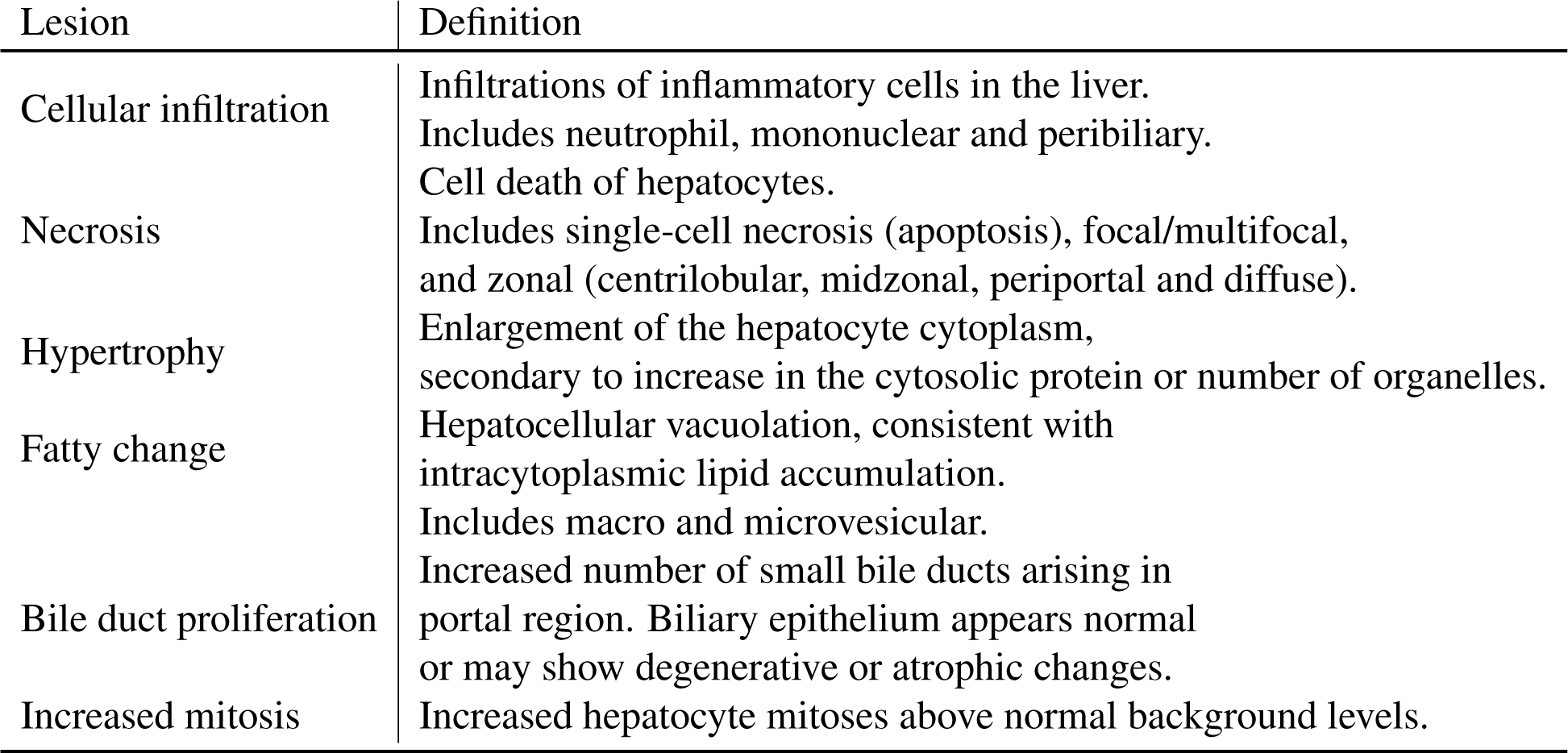
Morphological lesion definition. Morphological characterization of the lesions considered in this work are based on INHAND guidelines^78^, the International Harmonization of Nomenclature and Diagnostic Criteria, a publicly accessible resource that defines guidelines to diagnose lesions found in rodent toxicity and carcinogenicity studies.

**Table S5:** List of compounds used for inferring morphomolecular signatures for each lesion.

| Lesion | Compound |
| --- | --- |
| Necrosis | thioacetamide, ethionamide, methylene-dianiline, bromobenzene |
| Cellular Infiltration | ethionamide, thioacetamide, cyclophosphamide, methylene-dianiline |
| Bile duct proliferation | thioacetamide, methylene-dianiline |
| Fatty change | ethionamide, puromycin-aminonucleoside, cycloheximide, diazepam |
| Hypertrophy | hexachlorobenzene, diazepam, thioacetamide, hydroxyzine, ethionamide, methylene-dianiline, flutamide, phenacetin, bromobenzene, griseofulvin, methyltestosterone |
| Increased mitosis | methyltestosterone, griseofulvin, flutamide, ethionamide, puromycin aminonucleoside, danazol, hexachlorobenzene, methylene-dianiline |

**Table S6:**
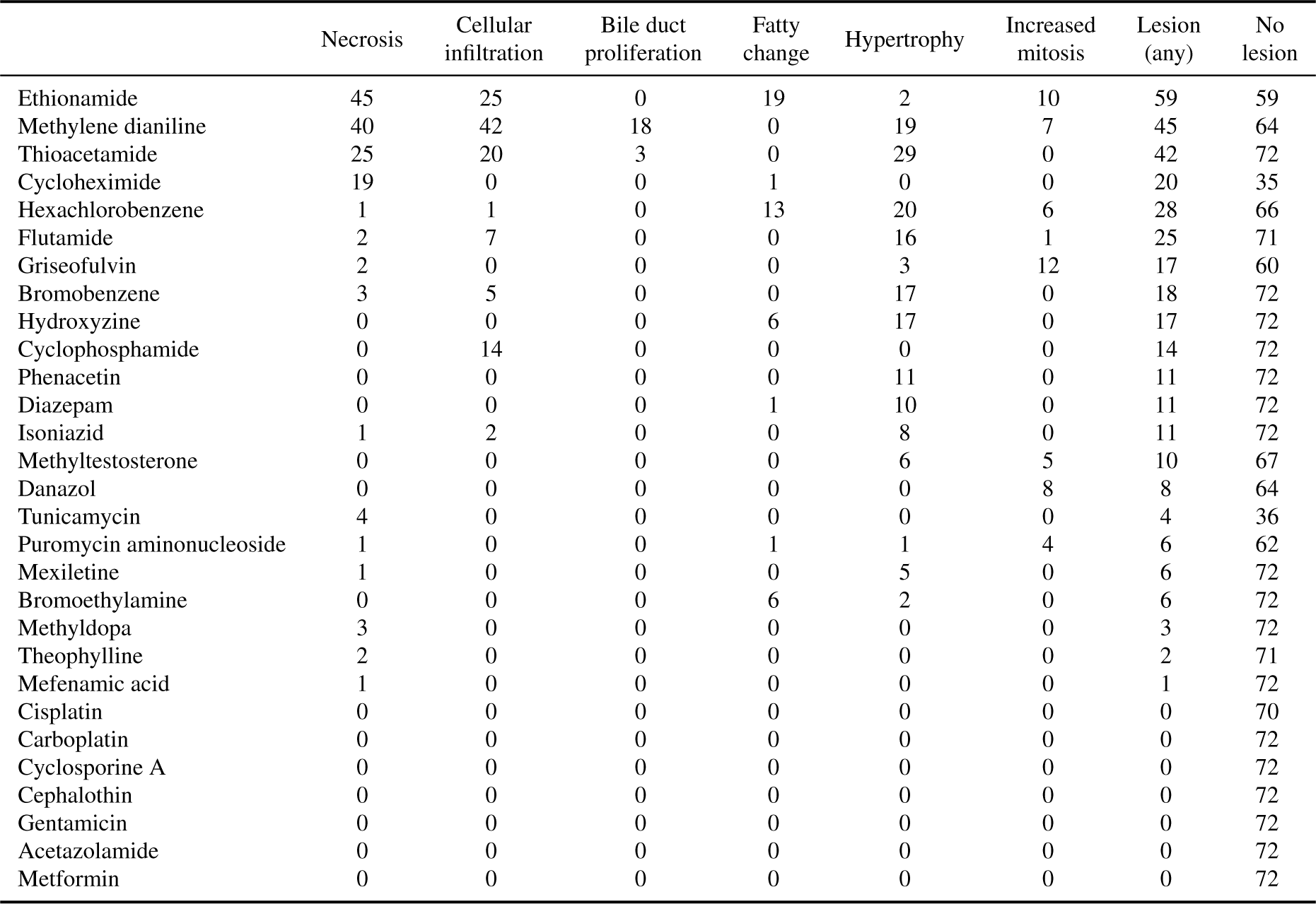
Distribution of lesions in TG-GATEs test set stratified by compound. Compounds are sorted by their percentage of slides with lesions.

**Table S7:** Number of patches with lesions per test study. Compounds are sorted by their percentage of slides with lesions.

|  | Necrosis | Cellular<br>infiltration | Bile duct<br>proliferation | Hypertrophy | Fatty<br>change | Increased<br>mitosis | Lesion<br>(any) | No<br>lesion |
| --- | --- | --- | --- | --- | --- | --- | --- | --- |
| Hexachlorobenzene | 2.53e+03 | 6.10e+03 | 3.28e+02 | 5.35e+05 | 5.01e+03 | 1.82e+04 | 5.60e+05 | 8.38e+05 |
| Thioacetamide | 2.24e+04 | 9.50e+03 | 4.85e+02 | 1.78e+05 | 1.83e+02 | 1.17e+04 | 2.14e+05 | 8.52e+05 |
| Diazepam | 6.35e+02 | 1.47e+03 | 7.90e+01 | 2.20e+05 | 8.88e+02 | 7.29e+03 | 2.29e+05 | 9.67e+05 |
| Ethionamide | 2.55e+04 | 9.04e+03 | 3.03e+02 | 9.02e+04 | 3.85e+04 | 3.15e+04 | 1.79e+05 | 9.70e+05 |
| Gentamicin | 5.38e+02 | 1.39e+03 | 2.63e+02 | 1.19e+04 | 1.50e+01 | 8.64e+04 | 1.00e+05 | 6.64e+05 |
| Cephalothin | 4.91e+02 | 1.97e+03 | 4.20e+02 | 1.17e+04 | 2.10e+01 | 8.03e+04 | 9.47e+04 | 6.81e+05 |
| Carboplatin | 2.13e+03 | 2.21e+03 | 1.24e+02 | 7.78e+04 | 1.90e+01 | 1.11e+04 | 9.31e+04 | 6.29e+05 |
| Acetazolamide | 1.89e+03 | 1.38e+03 | 2.30e+02 | 8.27e+04 | 1.18e+02 | 6.34e+03 | 9.24e+04 | 6.42e+05 |
| Puromycin aminonucleoside | 4.57e+03 | 3.69e+03 | 4.71e+02 | 6.22e+04 | 4.44e+02 | 1.22e+04 | 8.28e+04 | 6.28e+05 |
| Hydroxyzine | 4.40e+01 | 1.27e+03 | 3.10e+01 | 9.60e+04 | 3.37e+03 | 2.45e+02 | 9.76e+04 | 9.14e+05 |
| Cyclosporine A | 4.40e+03 | 2.77e+03 | 4.83e+02 | 4.21e+04 | 2.07e+02 | 1.81e+04 | 6.74e+04 | 6.68e+05 |
| Cycloheximide | 1.43e+04 | 2.99e+02 | 6.30e+01 | 2.24e+03 | 5.47e+03 | 1.12e+04 | 3.11e+04 | 3.19e+05 |
| Cisplatin | 1.33e+03 | 1.74e+03 | 3.13e+02 | 4.30e+04 | 1.13e+02 | 7.44e+03 | 5.35e+04 | 6.59e+05 |
| Isoniazid | 5.23e+03 | 6.84e+03 | 6.09e+02 | 5.64e+04 | 7.90e+01 | 6.86e+03 | 7.51e+04 | 1.08e+06 |
| Bromoethylamine | 8.59e+02 | 2.07e+03 | 1.59e+02 | 3.51e+04 | 5.80e+01 | 8.72e+03 | 4.68e+04 | 6.88e+05 |
| Methylene dianiline | 4.23e+03 | 8.30e+03 | 4.22e+03 | 4.25e+04 | 3.32e+03 | 2.92e+03 | 6.07e+04 | 9.91e+05 |
| Mexiletine | 1.75e+03 | 5.86e+02 | 3.30e+01 | 2.32e+04 | 6.00e+00 | 2.33e+04 | 4.88e+04 | 8.08e+05 |
| Flutamide | 2.09e+02 | 6.47e+03 | 2.58e+03 | 2.87e+04 | 1.10e+01 | 1.57e+04 | 5.16e+04 | 1.21e+06 |
| Theophylline | 7.78e+02 | 1.96e+02 | 1.36e+02 | 1.04e+04 | 3.10e+01 | 1.97e+04 | 3.12e+04 | 7.43e+05 |
| Tunicamycin | 7.99e+02 | 6.60e+01 | 6.00e+00 | 2.54e+03 | 1.00e+00 | 1.24e+04 | 1.58e+04 | 3.79e+05 |
| Danazol | 1.39e+02 | 2.56e+02 | 5.60e+01 | 2.26e+03 | 1.90e+01 | 2.64e+04 | 2.91e+04 | 7.47e+05 |
| Bromobenzene | 1.43e+03 | 6.70e+03 | 4.20e+01 | 1.95e+04 | 4.00e+01 | 6.83e+03 | 3.31e+04 | 9.27e+05 |
| Phenacetin | 1.14e+02 | 1.04e+03 | 3.20e+01 | 2.64e+04 | 3.90e+01 | 6.67e+02 | 2.82e+04 | 8.00e+05 |
| Griseofulvin | 1.16e+02 | 5.01e+03 | 1.82e+03 | 6.82e+03 | 8.00e+00 | 2.58e+04 | 3.82e+04 | 1.22e+06 |
| Methyltestosterone | 1.60e+02 | 2.05e+03 | 1.20e+01 | 2.37e+02 | 1.50e+01 | 2.47e+04 | 2.70e+04 | 1.06e+06 |
| Mefenamic acid | 3.31e+02 | 6.39e+03 | 2.68e+03 | 6.74e+03 | 3.00e+00 | 1.70e+04 | 3.10e+04 | 1.24e+06 |
| Cyclophosphamide | 3.07e+02 | 3.58e+02 | 8.00e+00 | 3.52e+03 | 2.00e+00 | 7.19e+03 | 1.14e+04 | 8.15e+05 |
| Methyldopa | 1.76e+02 | 1.81e+03 | 1.02e+02 | 1.80e+01 | 1.67e+02 | 5.56e+03 | 7.69e+03 | 9.84e+05 |
| Metformin | 1.81e+02 | 1.75e+03 | 4.70e+01 | 1.68e+02 | 5.10e+01 | 4.14e+03 | 6.23e+03 | 1.01e+06 |

**Table S8:** Genes indicative of general toxic exposure. Listed genes with the highest mean absolute predicted gene expression across patches with lesions. Analysis was conducted using five studies that report a large range of lesions: thioacetamide, methylene dianiline, ethionamide, bromobenzene, hexachlorobenzene.

|  | Necrosis | Cellular<br>infiltration | Bile duct<br>proliferation | Fatty<br>change | Hypertrophy | Increased<br>mitosis | Macro average |
| --- | --- | --- | --- | --- | --- | --- | --- |
| ABCC3 | 3.20 | 3.00 | 3.28 | 4.39 | 5.17 | 2.20 | 3.54 |
| GSTM3 | 2.64 | 2.71 | 2.93 | 3.10 | 3.12 | 1.29 | 2.63 |
| GPX2 | 3.60 | 2.77 | 3.11 | 2.42 | 2.75 | 1.00 | 2.61 |
| GSTA3 | 1.50 | 1.71 | 2.15 | 3.00 | 4.02 | 1.73 | 2.35 |
| ALDH1A1 | 0.84 | 1.43 | 2.28 | 2.58 | 3.64 | 1.58 | 2.06 |
| ALDH1A7 | 0.99 | 1.35 | 2.15 | 2.32 | 3.31 | 1.36 | 1.91 |
| ADAM8 | 2.48 | 2.22 | 2.81 | 1.57 | 1.37 | 0.40 | 1.81 |
| ATF3 | 3.27 | 2.64 | 1.96 | 1.44 | 0.98 | 0.50 | 1.80 |
| ACOT1 | 2.25 | 1.40 | 2.27 | 1.35 | 1.15 | 1.97 | 1.73 |
| TRIB3 | 2.84 | 2.01 | 1.31 | 1.50 | 1.65 | 0.65 | 1.66 |
| AKR7A3 | 1.26 | 1.30 | 1.02 | 2.19 | 2.56 | 1.59 | 1.66 |
| CCL2 | 3.12 | 2.54 | 2.15 | 1.29 | 0.70 | 0.02 | 1.64 |
| TNFRSF12A | 2.97 | 2.21 | 1.54 | 1.46 | 1.10 | 0.51 | 1.63 |
| AKR1B8 | 2.67 | 1.91 | 1.61 | 1.44 | 1.40 | 0.61 | 1.61 |
| ASNS | 2.33 | 1.76 | 1.87 | 1.29 | 1.69 | 0.59 | 1.59 |
| RND1 | 3.13 | 2.28 | 1.95 | 1.18 | 0.87 | 0.06 | 1.58 |
| PHGDH | 1.85 | 1.38 | 1.49 | 1.57 | 1.99 | 0.98 | 1.54 |
| OLR59 | -2.45 | -1.74 | -1.68 | -1.27 | -1.16 | -0.71 | -1.50 |
| ABCG8 | -2.06 | -1.54 | -1.80 | -1.47 | -1.38 | -0.86 | -1.52 |
| PKLR | -2.42 | -1.75 | -1.91 | -1.34 | -1.40 | -0.78 | -1.60 |
| MTMR7 | -2.09 | -1.76 | -2.05 | -1.50 | -1.78 | -0.69 | -1.64 |
| AKR1C1 | -2.73 | -1.95 | -1.97 | -1.46 | -1.44 | -0.77 | -1.72 |
| NREP | -3.33 | -2.02 | -1.89 | -1.38 | -1.47 | -0.43 | -1.75 |
| THRSP | -2.90 | -2.18 | -2.35 | -1.40 | -1.28 | -0.57 | -1.78 |
| OAT | -2.23 | -1.86 | -2.02 | -1.86 | -2.09 | -1.08 | -1.85 |
| LOX | -1.49 | -1.53 | -2.13 | -2.08 | -3.11 | -0.84 | -1.86 |
| SLC22A8 | -2.43 | -2.07 | -2.52 | -1.71 | -1.65 | -0.82 | -1.87 |
| BMF | -3.01 | -2.17 | -2.21 | -1.66 | -1.56 | -0.68 | -1.88 |
| INMT | -3.13 | -2.63 | -3.16 | -1.77 | -1.57 | -1.22 | -2.25 |
| NOX4 | -3.21 | -2.62 | -2.96 | -2.02 | -2.00 | -0.98 | -2.30 |
| CAR3 | -3.30 | -2.50 | -3.16 | -2.22 | -2.10 | -0.95 | -2.37 |
| OBP3 | -3.02 | -2.70 | -3.66 | -2.21 | -2.15 | -1.26 | -2.50 |
| STAC3 | -5.14 | -4.07 | -4.97 | -3.91 | -4.16 | -2.04 | -4.05 |

**Table S9:** List of 51 genes identified as being linked to increased mitosis. Analysis was conducted using eight compounds that reported increased mitosis. Patch-level average gene expression for patches containing mitosis compared to those identified with any of the five other lesions.

|  | Increased mitosis | Mean expression | Correlation prediction |
| --- | --- | --- | --- |
| CCNA2 | 2.52 | 0.33 | 0.40 |
| KNSTRN | 2.50 | -0.01 | 0.42 |
| CDKN3 | 2.41 | 0.01 | 0.27 |
| ECT2 | 2.32 | 0.31 | 0.51 |
| CENPW | 2.10 | 0.10 | 0.46 |
| CDK1 | 2.09 | 0.21 | 0.44 |
| CCNB1 | 2.05 | 0.20 | 0.45 |
| CDCA3 | 2.05 | 0.17 | 0.51 |
| SKA1 | 2.04 | 0.31 | 0.60 |
| MKI67 | 2.01 | 0.44 | 0.58 |
| TOP2A | 1.90 | 0.17 | 0.46 |
| KIF18B | 1.84 | 0.37 | 0.40 |
| AURKB | 1.81 | 0.39 | 0.59 |
| UBE2C | 1.80 | 0.25 | 0.47 |
| SPC24 | 1.77 | 0.12 | 0.70 |
| BUB1 | 1.73 | 0.27 | 0.57 |
| NUSAP1 | 1.73 | 0.13 | 0.52 |
| PTTG1 | 1.72 | 0.17 | 0.41 |
| IQGAP3 | 1.67 | 0.29 | 0.53 |
| CCNB2 | 1.67 | 0.15 | 0.39 |
| AURKA | 1.66 | 0.27 | 0.57 |
| NCAPH | 1.66 | 0.34 | 0.53 |
| HMMR | 1.52 | 0.26 | 0.53 |
| ARHGAP11A | 1.51 | 0.53 | 0.70 |
| ASPM | 1.49 | 0.22 | 0.56 |
| SKA3 | 1.48 | 0.27 | 0.68 |
| HJURP | 1.45 | 0.46 | 0.61 |
| ANLN | 1.30 | 0.29 | 0.49 |
| CENPF | 1.30 | 0.21 | 0.49 |
| FAM83D | 1.28 | -0.06 | 0.33 |
| BIRC5 | 1.17 | 0.17 | 0.37 |
| CDC20 | 1.07 | -0.09 | 0.61 |
| EXO1 | 0.99 | 0.21 | 0.35 |
| E2F1 | 0.96 | 0.37 | 0.57 |
| CKS2 | 0.93 | 0.24 | 0.53 |
| CDKN2C | 0.91 | 0.00 | 0.56 |
| HAUS4 | 0.88 | 0.10 | 0.49 |
| H2AX | 0.88 | 0.32 | 0.52 |
| RBL1 | 0.87 | 0.36 | 0.70 |
| TUBB5 | 0.84 | 0.41 | 0.75 |
| DCTPP1 | 0.83 | 0.32 | 0.41 |
| SMC4 | 0.75 | 0.14 | 0.33 |
| TK1 | 0.74 | 0.16 | 0.40 |
| MBOAT1 | 0.72 | 0.35 | 0.67 |
| H2AZ1 | 0.70 | 0.18 | 0.47 |
| MCM5 | 0.69 | 0.28 | 0.62 |
| CCNF | 0.68 | -0.01 | 0.51 |
| NETO2 | 0.67 | -0.09 | 0.50 |
| CCNE1 | 0.65 | 0.29 | 0.70 |
| PRIM1 | 0.58 | 0.15 | 0.65 |
| DNAJC9 | 0.54 | 0.13 | 0.47 |

**Table S10:** List of 66 genes identified as being linked to necrosis. Analysis was conducted using five compounds that reported necrosis. Patch-level average gene expression for patches containing necrosis compared to those identified with any of the five other lesions.

|  | necrosis | mean expression lesion | correlation prediction |
| --- | --- | --- | --- |
| ATF3 | 3.37 | 1.40 | 0.65 |
| TRIB3 | 3.23 | 1.47 | 0.62 |
| MAFF | 3.10 | 0.95 | 0.66 |
| TNFRSF12A | 3.09 | 1.30 | 0.79 |
| ZFAND2A | 2.87 | 0.97 | 0.75 |
| IFRD1 | 2.11 | 0.98 | 0.60 |
| MDM2 | 2.01 | 0.99 | 0.71 |
| FGF21 | 2.01 | 0.93 | 0.46 |
| MYC | 1.88 | 0.77 | 0.77 |
| CDKN1A | 1.87 | 0.92 | 0.35 |
| GADD45A | 1.87 | 0.80 | 0.62 |
| HSPB1 | 1.86 | 0.88 | 0.61 |
| SRXN1 | 1.84 | 0.84 | 0.63 |
| HSPA1A | 1.78 | 0.52 | 0.41 |
| SLC3A2 | 1.71 | 0.80 | 0.72 |
| DDIT3 | 1.70 | 0.75 | 0.49 |
| HMOX1 | 1.65 | 0.64 | 0.75 |
| AEN | 1.62 | 0.60 | 0.71 |
| TMBIM1 | 1.62 | 0.61 | 0.73 |
| LOC100911177 | 1.56 | 0.64 | 0.74 |
| LOC100912041 | 1.53 | 0.56 | 0.63 |
| FBXO30 | 1.37 | 0.57 | 0.68 |
| PPP1R15A | 1.36 | 0.46 | 0.74 |
| CCDC86 | 1.28 | 0.46 | 0.56 |
| JUN | 1.25 | 0.62 | 0.65 |
| HSPB8 | 1.24 | 0.45 | 0.75 |
| FOSL1 | 1.21 | 0.02 | 0.59 |
| AIMP2 | 1.20 | 0.53 | 0.80 |
| SLC7A11 | 1.15 | 0.43 | 0.38 |
| G6PD | 1.13 | 0.56 | 0.41 |
| TXNRD1 | 1.13 | 0.51 | 0.57 |
| RIOK3 | 1.07 | 0.36 | 0.50 |
| CXCL2 | 1.01 | 0.05 | 0.47 |
| ALDH4A1 | -1.02 | -0.37 | 0.80 |
| ACACA | -1.03 | -0.32 | 0.66 |
| AQP9 | -1.09 | -0.44 | 0.47 |
| RARB | -1.10 | -0.42 | 0.67 |
| CAR14 | -1.11 | -0.38 | 0.78 |
| FDFT1 | -1.12 | -0.18 | 0.40 |
| BHMT2 | -1.12 | -0.34 | 0.80 |
| IGFBP3 | -1.12 | -0.49 | 0.72 |
| CADPS2 | -1.13 | -0.47 | 0.66 |
| EBP | -1.13 | -0.39 | 0.74 |
| HGD | -1.16 | -0.49 | 0.72 |
| SUOX | -1.18 | -0.48 | 0.83 |
| GK | -1.21 | -0.57 | 0.74 |
| ARHGEF19 | -1.22 | -0.47 | 0.50 |
| ALBFM1 | -1.23 | -0.57 | 0.73 |
| VEPH1 | -1.23 | -0.27 | 0.63 |
| ACKR4 | -1.27 | -0.38 | 0.83 |
| ADAMTS7 | -1.30 | -0.63 | 0.65 |
| CTH | -1.31 | -0.33 | 0.54 |
| HMGCS1 | -1.36 | -0.38 | 0.32 |
| PRLR | -1.41 | -0.29 | 0.33 |
| HDHD3 | -1.43 | -0.58 | 0.38 |
| HES6 | -1.53 | -0.55 | 0.64 |
| ABCA8A | -1.55 | -0.72 | 0.86 |
| ACSS2 | -1.75 | -0.45 | 0.66 |
| COQ8A | -1.86 | -0.74 | 0.82 |
| CXCL12 | -2.04 | -0.91 | 0.79 |
| FADS1 | -2.07 | -0.64 | 0.63 |
| XPNPEP2 | -2.15 | -0.93 | 0.72 |
| ACACB | -2.18 | -0.65 | 0.58 |
| CYP7A1 | -2.36 | -0.67 | 0.66 |
| AVPR1A | -2.52 | -0.93 | 0.83 |
| NREP | -3.30 | -1.45 | 0.80 |

**Table S11:** List of 27 genes identified as being linked to cellular infiltration. Analysis was conducted using five compounds that reported cellular infiltration. Patch-level average gene expression for patches containing cellular infiltration compared to those identified with any of the five other lesions.

|  | cellular infiltration | mean expression lesion | correlation prediction |
| --- | --- | --- | --- |
| CXCL1 | 2.38 | 0.92 | 0.41 |
| BCL2A1 | 1.67 | 0.71 | 0.48 |
| S100A4 | 1.66 | 0.78 | 0.41 |
| CXCL10 | 1.63 | 0.72 | 0.31 |
| S100A8 | 1.49 | 0.56 | 0.50 |
| CHI3L1 | 1.32 | 0.54 | 0.58 |
| S100A9 | 1.23 | 0.38 | 0.36 |
| IL1B | 1.17 | 0.41 | 0.51 |
| CAPG | 1.11 | 0.54 | 0.55 |
| TGFB1 | 1.06 | 0.35 | 0.59 |
| PTPRC | 0.91 | 0.35 | 0.37 |
| ACKR3 | 0.88 | 0.35 | 0.46 |
| FILIP1L | 0.83 | 0.35 | 0.47 |
| C5AR1 | 0.77 | 0.37 | 0.42 |
| CD83 | 0.76 | 0.29 | 0.40 |
| EVI2A | 0.74 | 0.26 | 0.48 |
| LCP1 | 0.73 | 0.29 | 0.55 |
| ICAM1 | 0.72 | 0.33 | 0.56 |
| CYBB | 0.70 | 0.24 | 0.43 |
| IL7 | 0.69 | 0.14 | 0.44 |
| CASP1 | 0.67 | 0.28 | 0.49 |
| PSMB9 | 0.64 | 0.30 | 0.41 |
| GLIPR1 | 0.62 | 0.29 | 0.55 |
| AP1S2 | 0.57 | 0.23 | 0.40 |
| PLVAP | 0.54 | -0.04 | 0.31 |
| SRGN | 0.53 | 0.22 | 0.39 |
| IFITM2 | 0.52 | 0.26 | 0.68 |

**Table S12:**
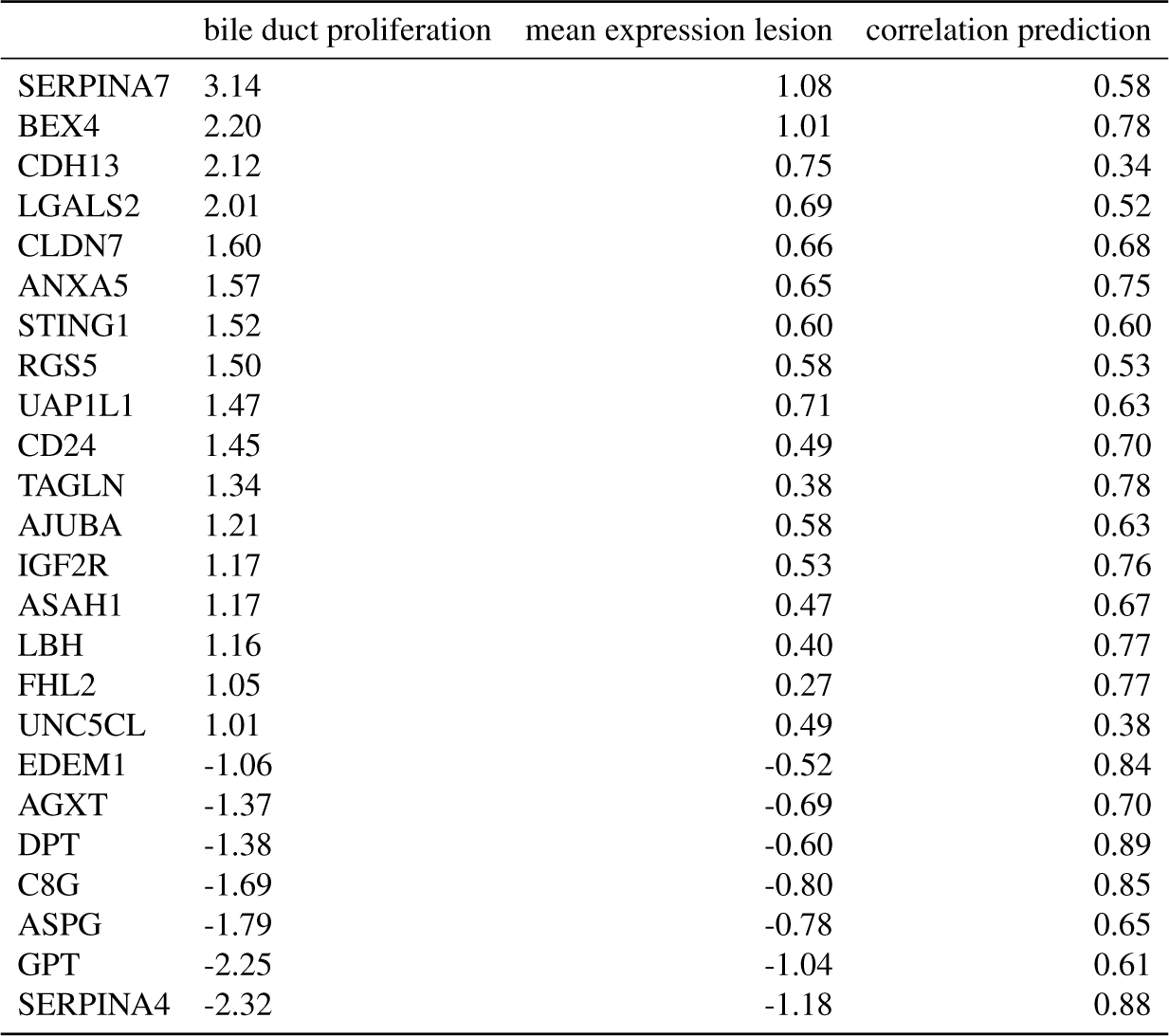
List of 24 genes identified as being linked to bile duct proliferation. Analysis was conducted using two compounds that reported bile duct proliferation. Patch-level average gene expression for patches containing bile duct proliferation compared to those identified with any of the five other lesions.

**Table S13:** List of 14 genes identified as being linked to fatty change. Analysis was conducted using four compounds that reported fatty change. Patch-level average gene expression for patches containing fatty change compared to those identified with any of the five other lesions.

|  | fatty change | mean expression lesion | correlation prediction |
| --- | --- | --- | --- |
| ACOT1 | 1.93 | 0.47 | 0.59 |
| GSTP1 | 1.48 | 0.58 | 0.40 |
| CIDEA | 1.04 | 0.38 | 0.30 |
| ACOT2 | 0.83 | 0.13 | 0.31 |
| TMED3 | 0.75 | 0.35 | 0.53 |
| CYP1A1 | 0.75 | -0.06 | 0.63 |
| HID1 | 0.71 | 0.18 | 0.54 |
| FKBP11 | 0.65 | 0.25 | 0.72 |
| IRAK1 | 0.63 | 0.26 | 0.46 |
| MANBA | 0.56 | 0.17 | 0.65 |
| ACP3 | -0.52 | -0.21 | 0.35 |
| EPSTI1 | -0.58 | -0.15 | 0.42 |
| C5 | -0.61 | -0.22 | 0.30 |
| SLC6A6 | -1.01 | -0.48 | 0.48 |

**Table S14:** List of 23 genes identified as being linked to hypertrophy. Analysis was conducted using eleven compounds that reported hypertrophy. Patch-level average gene expression for patches containing hypertrophy compared to those identified with any of the 5 other lesions.

|  | hypertrophy | mean expression lesion | correlation prediction |
| --- | --- | --- | --- |
| GSTA3 | 3.90 | 1.52 | 0.72 |
| ALDH1A1 | 3.56 | 1.43 | 0.76 |
| CYP1A1 | 3.23 | 0.25 | 0.43 |
| ALDH1A7 | 3.19 | 1.31 | 0.43 |
| ACSM2 | 1.44 | 0.33 | 0.35 |
| RGD1559459 | 1.33 | 0.44 | 0.77 |
| ADGRG2 | 1.32 | 0.45 | 0.59 |
| VNN1 | 1.24 | 0.28 | 0.51 |
| ME1 | 1.18 | -0.26 | 0.43 |
| EPHX1 | 1.04 | 0.31 | 0.46 |
| POR | 0.91 | 0.11 | 0.54 |
| GCLM | 0.71 | 0.00 | 0.53 |
| GCLC | 0.62 | 0.05 | 0.46 |
| GSTA4 | 0.60 | -0.06 | 0.72 |
| TFRC | 0.58 | 0.04 | 0.46 |
| ADH4 | 0.51 | -0.29 | 0.45 |
| MCRIP2 | 0.51 | 0.08 | 0.68 |
| GSTM2 | 0.50 | -0.16 | 0.64 |
| NAPSA | -0.54 | 0.45 | 0.31 |
| PLVAP | -0.85 | 0.23 | 0.37 |
| SLC34A2 | -1.08 | -0.25 | 0.37 |
| SLC6A6 | -1.28 | -0.38 | 0.44 |
| LOX | -2.97 | -1.32 | 0.56 |

**Table S15:** iBOT hyperparameters used SSL pretraining. 4 *×* 80GB NVIDIA A100 GPUs were used for training. Batch size refers to the total batch size across GPUs.

| Hyper-parameter | Value |
| --- | --- |
| Layers | 24 |
| Heads | 16 |
| Patch size | 16 |
| FFN layer | MLP |
| Head activation | GELU |
| Embedding dimension | 1024 |
| Stochastic dropout rate | 0.1 |
| Global crop scale | 0.48, 1.0 |
| Global crop number & size | 2, 224 |
| Local crop scale | 0.16, 0.48 |
| Local crop number & size | 8, 96 |
| Max masking ratio | 0.5 |
| Min masking ratio | 0.1 |
| Gradient clipping max norm | 3.0 |
| Normalize last layer | ✓ |
| Shared head | ✗ |
| AdamW $\beta$ | (0.9, 0.999) |
| Batch size | 3072 |
| Freeze last layer iterations | 1250 |
| Warmup iterations | 12500 |
| Warmup teacher temperature iterations | 37500 |
| High-resolution finetuning iterations | 12500 |
| Max Iterations | 125000 |
| Learning rate schedule | Cosine |
| Learning rate (start) | 0 |
| Learning rate (post warmup) | 2e-3 |
| Learning rate (final) | 1e-6 |
| Teacher temperature (start) | 0.04 |
| Teacher temperature (final) | 0.4 |
| Teacher momentum (start) | 0.992 |
| Teacher momentum (final) | 1.000 |
| Weight decay (start) | 0.04 |
| Weight decay (end) | 0.4 |
| Automatic mixed precision | fp16 |

**Table S16:** Hyperparameters used in GEESE classification. A single 24GB NVIDIA GeForce RTX 3090 GPU was used for each MIL model using weakly-supervised learning and slide-level labels.

| Hyperparameter | Value |
| --- | --- |
| Batch size | 1 |
| Weight decay | 1e-5 |
| AdamW $\beta$ | (0.9, 0.999) |
| Peak learning rate | 1e-4 |
| Learning rate schedule | Cosine |
| Epochs | 20 |

## Footnotes

1 github.com/bytedance/ibot

2 syngoportal.org/convert

3 https://rgd.mcw.edu/rgdweb/enrichment/start.html

4 https://ctdbase.org/

5 https://dbarchive.biosciencedbc.jp/en/open-tggates/desc.html

6 https://zenodo.org/record/7541930

7 https://toxygates.nibiohn.go.jp/toxygates/

